# A bacterial inflammation sensor regulates c-di-GMP signaling, adhesion, and biofilm formation

**DOI:** 10.1101/2020.10.11.335190

**Authors:** Arden Perkins, Dan A. Tudorica, Raphael D. Teixeira, Tilman Schirmer, Lindsay Zumwalt, O. Maduka Ogba, C. Keith Cassidy, Phillip J. Stansfeld, Karen Guillemin

**Author notes:** Corresponding authors: AP, KG.

## Abstract

The reactive oxygen species produced during inflammation through the neutrophilic respiratory burst play profound roles in combating bacterial pathogens and regulating the microbiota. Among these, the neutrophilic oxidant bleach, hypochlorous acid (HOCl), is the most prevalent and strongest oxidizer and kills bacteria through non-specific oxidation of proteins, lipids, and DNA. Thus, HOCl can be viewed as a host-specific cue that conveys important information about what bacterial physiology and lifestyle programs may be required for successful colonization. Nevertheless, bacteria that colonize animals face a molecular challenge in how to achieve highly selective detection of HOCl due to its reactive and transient nature and chemical similarity to more benign and non-host-specific oxidants like hydrogen peroxide (H_2_O_2_). Here, we report that in response to increasing HOCl levels *E. coli* regulates biofilm production via activation of the diguanylate cyclase DgcZ. We show the molecular mechanism of this activation to be specific oxidation of a conserved cysteine that coordinates the zinc of its regulatory chemoreceptor zinc-binding (CZB) domain, forming a zinc-cysteine redox switch 685-fold more sensitive to oxidation by HOCl over H_2_O_2_. Dissection of the signal transduction mechanism through quantum mechanics, molecular dynamics, and biochemical analyses reveal how the cysteine redox state alters the delicate equilibrium of competition for Zn^++^ between the CZB domain and other zinc binders to relay the presence of HOCl through activating the associated GGDEF domain to catalyze c-di-GMP. We find biofilm formation and HOCl-sensing *in vivo* to be regulated by the conserved cysteine, and point mutants that mimic oxidized CZB states increase production of the biofilm matrix polymer poly-N-acetylglucosamine and total biofilm. We observe CZB-regulated diguanylate cyclases and chemoreceptors in phyla in which host-associated bacteria are prevalent and are possessed by pathogens that manipulate host inflammation as part of their colonization strategy. A phylogenetic survey of all known CZB sequences shows these domains to be conserved and widespread across diverse phyla, suggesting CZB origin predates the bacterial last universal common ancestor. The ability of bacteria to use CZB protein domains to perceive and thwart the host neutrophilic respiratory burst has implications for understanding the mechanisms of diseases of chronic inflammation and gut dysbiosis.

## INTRODUCTION

During inflammation neutrophils use the enzyme myeloperoxidase to catalyze the potent reactive oxygen species hypochlorous acid (HOCl) in order to control and eliminate invading bacteria, and sites of inflamed tissue can harbor millimolar concentrations of HOCl^1,2^. This presents a significant obstacle for bacteria that colonize animals, as HOCl is an extremely reactive chemical that acts as a bactericide through the oxidation of a broad spectrum of cellular components, especially sulfur-containing amino acids^3–5^. However, only trace amounts of HOCl form in the absence of enzymatic catalysis, and so HOCl also represents a unique chemical cue that signals the presence of an animal host. The ability to detect HOCl in the environment can allow bacteria to optimize their lifestyle for host colonization and/or enduring host inflammation; but achieving high selectivity and sensitivity of a reactive and labile chemical is not an easy task. A second challenge is that other chemically-similar oxidants exist, such as hydroperoxides (ROOH), that do not necessarily convey the same information about a bacterium’s environment. For example, hydrogen peroxide (H_2_O_2_), which bacteria are well-equipped to eliminate^6,7^, is far less toxic^8^ and is produced through many mechanisms other than immune responses, such as bacterial metabolism^9^, and so its presence is not host-specific per se. Thus, the detection of HOCl requires a sensing apparatus to overcome the oxidant’s propensity for non-specific oxidation and to distinguish between other oxidants that can react with a similar suite of molecular targets. In fact, synthetic molecular probes have only recently overcome this challenge^10,11^.

We recently identified a novel bacterial sensing system in which a chemoreceptor zinc-binding (CZB) protein domain detects the inflammation product HOCl through direct oxidation of a reactive cysteine to form cysteine sulfenic acid (Cys-SOH, Fig. S1)^12,13^. The molecular mechanism is thought to accomplish selective reactivity with HOCl through a conserved zinc-thiolate switch^12,13^, a chemical moiety with enhanced reactivity toward HOCl^5,14,15^. As an example of the biological significance of this mechanism and relevance to human disease, CZB sensing of HOCl was shown to facilitate chemoattraction to HOCl sources for the gastric pathogen *H. pylori* through regulation of the chemoreceptor transducer-like protein D (TlpD), providing an explanation for the bacteria’s persistence in inflamed tissue and tropism for gastric wounds^12,16–18^. Observations of other CZB-containing Gammaproteobacteria that can inhabit inflamed environments, such as *Salmonella enterica*^19^ and *Escherichia coli*^20^, prompted us to investigate whether sensing of host HOCl by CZB proteins could have broad utility in regulating bacterial lifestyles critical for host colonization.

CZB domains remain poorly characterized, but CZB-containing proteins are reported to play roles in sensing exogenous zinc, pH, and oxidants to regulate bacterial chemoreceptors and diguanylate cyclases^12,21–25^. Earlier work has shown CZBs to be approximately 15 kD in size and consist of a four-helix bundle fold with a unique and conserved 3His/1Cys zinc-binding motif^23,26^, with the cysteine of this motif stabilized as a thiolate and activated to be oxidized by HOCl^13^. Prior to this study, it was unknown by what molecular mechanism CZB domains could integrate signals from such divergent ligands as HOCl and Zn^++^ into cellular responses. Cellular zinc homeostasis involves a complex equilibrium between many high affinity zinc binders that compete for Zn^++^ and maintain the cytosolic Zn^++^ concentration near zero^27,28^, and CZB affinity for zinc is thought to be in the sub-femtomolar range^23^. In the highly competitive environment of the cell cytosol even small changes in a protein’s zinc affinity can dramatically shift the binding equilibrium between zinc-bound and zinc-free states^27–30^. Thus, we sought to study whether alterations to the CZB zinc-binding core through HOCl oxidation might regulate the domain through influencing zinc binding. However, a critical barrier to investigating this hypothesis, both *in vitro* and *in vivo*, is the transient nature of the Cys-SOH reaction intermediate, which can spontaneously reduce or become further oxidized to cysteine-sulfinate (Cys-SO_2_^−^) in the presence of excess oxidant^31,32^. So far, *in vivo* studies on the biological roles of CZBs have mostly relied on full-knockouts^12,16,21,23,33^ and the importance of the conserved zinc-binding cysteine has not previously been investigated *in vivo*.

The ubiquitous bacterial signaling molecule bis-(3′-5′)-cyclic dimeric guanosine monophosphate (c-di-GMP) is well-known to play a pivotal role in bacterial decisions of cell adhesion and biofilm formation, that in turn have relevance for the pathogenicity of many bacteria involved in human diseases^34–36^. The existence of CZB-regulated diguanylate cyclases, which catalyze the production of c-di-GMP from guanosine triphosphate (GTP), prompted us to ask if c-di-GMP signaling processes, such as biofilm formation, can be regulated in response to HOCl, and thus, to host inflammation. To address this question, we have for the first time quantified CZB conservation patterns to understand the underlying functional rationale as it pertains to ligand-sensing and signal transduction, determined the prevalence of CZB-containing protein architectures to identify the molecular pathways they regulate, and documented the full biological distribution of these proteins to learn the breadth of environments and organisms in which these proteins operate. These data reveal the *E. coli* diguanylate cyclase Z (DgcZ, previously referred to as YdeH^23,37–39^) as an exemplar CZB-regulated diguanylate cyclase representative of a large subset of CZB-containing proteins, and we have utilized DgcZ as a model system to obtain broad insight into CZB HOCl-sensing and regulation of bacterial biofilm formation.

We find that *E. coli* DgcZ, is highly and preferentially reactive with biologically-relevant concentrations of HOCl and regulates c-di-GMP catalysis and cellular biofilm through direct oxidation of the zinc-binding cysteine. Biofilm and surface attachment is increased by micromolar HOCl through DgcZ and can also be induced by strains harboring a DgcZ point mutant that mimics cysteine oxidation. These data support earlier reports that oxidants can increase *E. coli* biofilm formation^24,40^ and provide new insight into how enteropathogenic, enteroaggregative, and uropathogenic *E. coli* (EPEC, EAEC, UPEC) may respond to host inflammation to favor pathogenicity^41–43^. We propose a new unifying model for how CZB proteins can facilitate both HOCl and zinc-sensing, with relevance for bacterial biology across diverse phyla and human diseases of gut dysbiosis and chronic inflammation. The ability of CZB domains to selectively sense HOCl implicates this family of proteins as bacterial inflammation sensors.

## RESULTS

### Architectures, conservation, and biological distribution of CZB-containing proteins

Protein structure-function relationships can be revealed through analyses of conservation patterns to learn what parts of the protein are indispensable for function across divergent homologues^44–46^. To create a database of all currently-known CZB-containing proteins we performed iterative searches with the Basic Local Alignment Search Tool (BLAST)^47^ for CZB domain sequences in the non-redundant protein database, which resulted in 8,227 sequences that contain the unique 3His/1Cys CZB zinc-binding motif, with 22.7 % pairwise identity across all sequences. Most CZB-containing protein sequences contain multiple protein domains, so to understand the major cellular pathways regulated by CZB domains, we further categorized and quantified sequences according to protein architecture. We found that CZB-containing proteins can be divided into seven subgroups based on domain similarity, with the majority of sequences (85.8 %) involved in two biological outputs, namely chemotaxis or c-di-GMP metabolism (Fig. 1A-B). Some sequences also appear to contain only a CZB domain with no other detectable protein domain sequence signature (12.6 %), although some of these may represent incomplete sequences or annotations. The most common subgroup consists of soluble CZB-regulated chemoreceptors similar in structure to *H. pylori* TlpD (59.4 %), which we refer to here as “TlpD-like.” CZB-regulated nucleotide cyclases, including *E. coli* DgcZ, account for a smaller but widespread fraction of sequences (6.3 %), which we refer to as “DgcZ-like.” Less common, but involved in functionally-related processes, are CZB-regulated chemotaxis W (CheW, 1.8 %) and Glu-Ala-Leu (EAL) (0.6 %) proteins that transduce chemoreceptor signals and degrade c-di-GMP, respectively. Nearly all CZB sequences are predicted to be cytosolic, with only 384 putative periplasmic CZB sequences (approximately 4 %) identified.

**Fig. 1.**
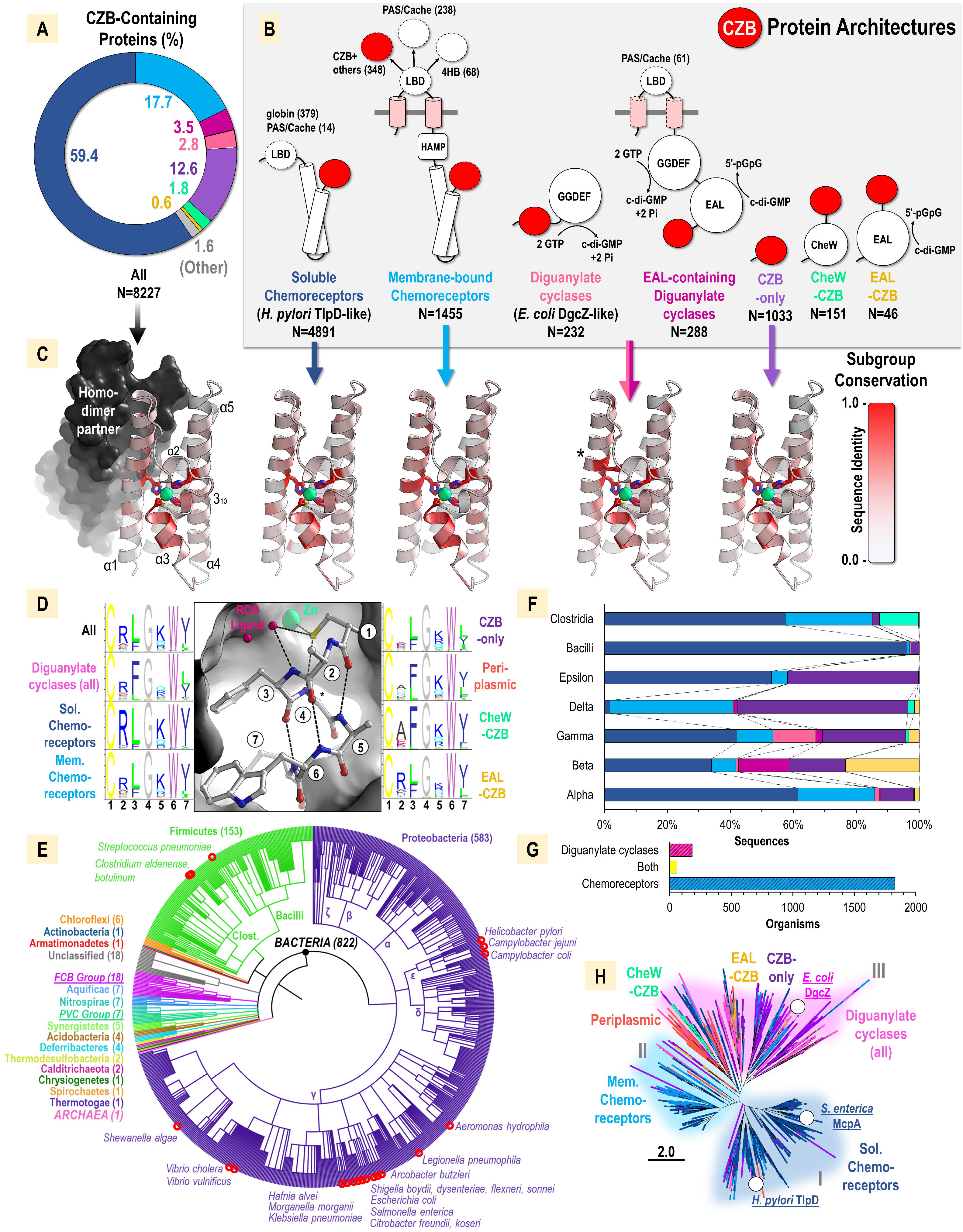
Amino acid conservation and biological distribution of CZB domains. A-B. Architectures of CZB (red circle) domain-containing proteins. Membrane-spanning regions are noted in pink. Structural features found in some, but not all, members of a subgroup are noted with dashed lines, with variations in ligand-binding domain (LBD) quantified in parentheses. Observed domains include four-helix bundle (4HB), globin, Pas/Cache (and tandem PAS/dCache), histidine kinases adenylate cyclases methyl accepting proteins phosphatase (HAMP), methyl accepting chemotaxis proteins (MCP, here referred to as chemoreceptors), CheW-like, GGDEF, and EAL. C. Amino acid conservation of CZB subgroups mapped onto the crystal structure of *E. coli* DgcZ CZB. D. Amino acid conservation of the α3 CZB motif. Conservation for putative periplasmic CZBs from the N-terminal region of membrane-bound chemoreceptors are also shown and noted as “periplasmic.” E. Phylogenetic tree showing the biological distribution of CZB domains colored by phyla (bold), superphyla (bold, italicized, underlined), or kingdom (bold uppercase). Number of organisms identified down to the species level in each group is noted in parentheses. Classes of Firmicutes and Proteobacteria are labeled. Bacteria associated with causing disease in humans are indicated with red circles. F. CZB subgroups found in Firmicutes and Proteobacteria classes. Coloring as in B. G. Quantification of organisms that at the species level contain only a CZB-containing chemoreceptor, only a diguanylate cyclase, or both. H. Relatedness tree of CZB domains from subgroups with clusters I-III highlighted in indigo, cyan, and pink.

Commonalities in amino acid conservation between distantly related CZB sequences could point toward general functions of CZB domains, while differences between subgroups could indicate evolutionary tuning to optimize ligand-sensing and signal transduction in specific settings. To understand these patterns at a structural level, we mapped amino acid conservation onto the crystal structure of the *E. coli* DgcZ CZB domain, which is a homodimer with each monomer composed of five α-helices and one 3_10_-helix (Fig 1C, PDB code: 3t9o)^23,48^. In addition to the ubiquitous 3His/1Cys zinc-binding motif, two regions of global conservation across all CZB domains were revealed (Fig. 1C). First, the N-terminal α1 helix exhibits a modest degree of conservation, with 10 positions that have sequence identity conservation in the range of 20-100 %. This region (residues 1-30) constitutes a large portion of the homodimer interface (384 Å^2^ of 1950 Å^2^ total) that forms a two-fold symmetry axis, with residues packing against their homodimer counterpart. Second, in addition to the universally-conserved zinc-binding Cys, many residues of the α3 region exhibit a high degree of conservation (Fig. 1C). This pattern of conservation in the α1 and α3 regions occurs across all CZB subgroups, suggesting these two regions are of universal importance for CZB function. One additional site of high conservation occurs in the diguanylate cyclase subgroup, where a Trp residue, which resides three sites downstream of the conserved zinc-binding His22, packs into the protein core against the zinc-binding site (Fig. 1C, noted with Asterix).

Oxidation of the conserved zinc-binding Cys, located in α3, was previously reported to induce structural changes in the CZB domain, whereby Cys-SOH formation promotes detachment of the Cys from the zinc core and a local unfolding of the α2-α3 region^12^ (Fig. S1). The structural change upon disruption of the Cys thiolate-zinc interaction is also directly observed in the crystal structure of a DgcZ Cys→Ala mutant (PDB: 4h54)^23^. Therefore, the high conservation of the α3 region could relate to CZB signal transduction. To further investigate the conservation of the α3 region, seqlogo plots were generated for the seven-residue motif containing the conserved Cys for all CZB sequences and CZB architecture subgroups (Fig. 1D). By studying the position and interactions of each residue in the *E. coli* DgcZ CZB structure, putative roles and rationales for conservation were inferred for each amino acid site as follows: Position 1 is approximately 100 % conserved as a Cys, reflecting its absolute requirement for function. The Cys forms part of the zinc-binding core, increases zinc affinity by an order of magnitude^23^, and can serve as a redox-sensor through direct oxidation in some CZB proteins^12^. Positions 2 and 5 are conserved as residues that either contain a hydrophilic side chain or a small hydrophobic side chain that permit exposure to solvent. Positions 3, 6, and 7 are conserved as bulky hydrophobic side chains that are buried in the protein core and provide a thermodynamic driving force for the proper folding of the motif. Position 4 is almost universally conserved as a Gly likely for the reason that a C_β_ atom would clash with the carboxyl oxygen of a position three residues upstream of the Cys in the 3_10_ helix, and the position does not adopt phi-psi angles that are Gly-specific (φ=122.3°, ψ=114.9°). A site of variability between subgroups is Position 2, which is enriched as Arg especially in soluble chemoreceptors but also membrane-bound chemoreceptors, diguanylate cyclases, and EAL-CZBs (Fig. 1D). However, Position 2 is either poorly conserved or conserved as an Ala for the CZB-only, periplasmic, and CheW-CZBs subgroups. If the α3 region exists as a dynamic equilibrium, with the locally-unfolded state serving to relay the signal of ligand-sensing, these amino acid substitutions may be the result of evolutionary tuning to shift that equilibrium to best suit a CZB domain’s signaling role for a given CZB subgroup.

To obtain insight into the biological factors that drive CZB evolution we assessed the phylogenetic distribution of CZB domains. This revealed a total of 822 unique organisms for which phylogenetic classification and annotation were available down to the species level. This analysis shows that CZB domains are found in diverse phyla, and essentially all CZB sequences are bacterial (Fig. 1E). A single eukaryotic sequence from the nematode *Diploscapter pachys* was nearly identical to a sequence from *Pseudomonas* and, therefore, likely contamination. A single archaeon sequence from *Canditatus Woesearchaeota* shows 35 % sequence similarity to sequences from the bacterial genus *Sulfurimonas* and may represent a case of horizontal gene transfer. In total, we report that CZB domains are present in 21 bacterial phyla and 6 candidate phyla (Fig. 1E), which expands on a previous observation of these proteins in 7 phyla^26^. An evolutionary divergence tree for species that contain CZB proteins suggests they may have been present in the bacterial last universal common ancestor (LUCA) more than 4 billion years ago (Fig. S2)^49^.

In terms of currently available sequence data, most CZB proteins are found in Firmicutes and Proteobacteria, phyla that contain many host-associated species, and Gammaproteobacteria account for approximately one-third of all known CZB sequences (Fig. 1E). CZB domains were identified in 24 bacterial species associated with human diseases, including many enteric pathogens such as species from the genera *Vibrio*, *Shewanella*, *Shigella*, *Helicobacter*, *Campylobacter*, *Escherichia*, *Salmonella,* and *Citrobacter*, as well as pathogens associated with nosocomial infections such as *Morganella* and *Klebsiella* (Fig. 1E red circles). There exist considerable differences among the Proteobacteria and Firmicutes classes in the prevalence of CZB protein architectures (Fig. 1F). Regarding subgroups involved in chemotaxis, soluble chemoreceptors make up a sizable fraction in these classes except for Deltaproteobacteria, in which they are nearly absent, and CheW-CZBs are most abundant in Clostridia. Diguanylate cyclases and EAL-CZBs are restricted to Alpha-, Beta-, Gamma-, and Deltaproteobacteria, suggesting CZBs as important regulators of c-di-GMP signaling for these bacteria. At the species level, we were intrigued to find that bacteria seem to contain either CZB-regulated chemoreceptors or CZB-regulated diguanylate cyclases, but not both. This suggesting the signals perceived by CZB domains are exclusively integrated into bacterial regulation of either chemotaxis or c-di-GMP signaling (Fig. 1G).

If CZB domains serve a singular molecular-sensing purpose, and are readily swappable regulatory units, that might result in homogenous amino acid conservation irrespective of subgroup. Alternatively, if CZB domains have become honed over time for more specific functions that could result in discrete clustering for subgroups. Additionally, knowledge of CZB sequence similarity can inform the selection of a specific CZB-containing protein for further study that is representative of a large group of CZBs and can serve as a model system to obtain broad insight into CZB function. A relatedness tree for full-length CZB amino acid sequences shows approximately three main clusters exist: cluster I is dominated by C-terminal soluble chemoreceptor CZBs, including *H. pylori* TlpD and *S. enterica* methyl-accepting chemotaxis protein A (McpA), cluster II by C-terminal membrane-bound chemoreceptor CZBs, and cluster III is more variable and contains diguanylate cyclases, including *E. coli* DgcZ, periplasmic (N-terminal chemoreceptor CZBs), CheW-CZB, EAL-CZB, and CZB-only subgroups (Fig. 1H). From these results, we identified *E. coli* DgcZ as a CZB-containing protein that is representative of a diverse array of CZB subgroups (i.e. cluster III) and that could be leveraged to provide new and broad insight into the mechanism and biological utility of CZB HOCl-sensing, especially as it pertains to regulation of c-di-GMP signaling. *E. coli* DgcZ has several other advantages that led us to choose it as a model system: i) it contains the conserved zinc-binding core and reactive cysteine, ii) the recombinant protein is soluble, stable, permits mutagenesis, and can be produced with high yields for *in vitro* analyses, iii) high-resolution crystal structures have been determined for the full-length protein and the individual CZB domain^23^, iv) its small size facilitates computational analyses for molecular modeling of ligand sensing and signal transduction, and v) the bacterium is genetically-tractable and readily forms biofilms.

### Computational Dissection of HOCl Signal Transduction

Previous analyses suggested *E. coli* DgcZ, and perhaps many other CZB domain-containing proteins, acts as an HOCl sensor via the inherently HOCl-reactive moiety of its Cys-Zn redox switch^12,13^. Since we hypothesized that CZB protein domains may function to relay signals about HOCl concentrations, and hence inflammation, in a bacterium’s environment, we sought to better understand the molecular mechanism by which these signals could be transduced. To this end, we utilized molecular dynamics (MD) simulation to model the CZB domain homodimer from *E. coli* DgcZ^23^ (Fig. 2A) with specific alterations to the conserved zinc-binding cysteine. We examined three redox states, namely Cys52-S^−^ (native unreacted thiolate state when bound to zinc), Cys52-SH, and Cys52-SOH (product of HOCl reaction), as well as two mutations, C52A and C52D. For each state, triplicate simulations were performed for a duration of 1 μs each, allowing six views of the active site (Movies S1-5).

**Fig. 2.**
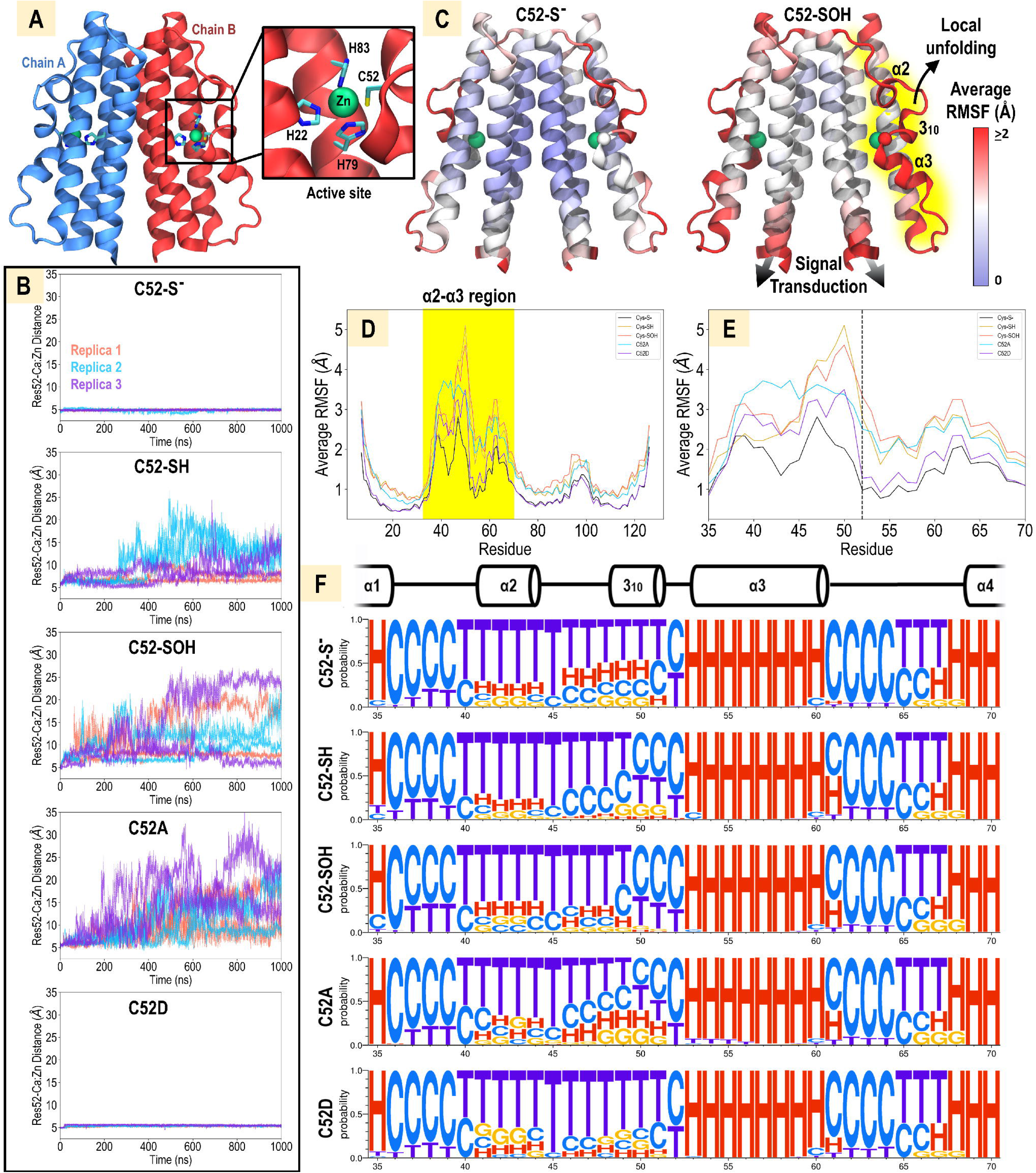
MD simulation of CZB dimer with variable residue 52 moieties. A. Starting model of the *E. coli* CZB domain dimer based on crystal structure 3t9o^23^. B. Local unfolding events in the α2-α3 region for each model as monitored by the distance between Zn and the residue 52 Cα atoms. Data from three independent simulations are shown (red, blue, purple), with independent active sites from each simulation noted with solid and dashed lines. C. RMSFs, averaged across three simulations each, for the C52-S^−^ and C52-SOH models are shown mapped to the CZB structure. The α2-α3 region involved in local unfolding is highlighted in yellow. Black arrows denote structural changes that may be involved in signal transduction. The zinc and residue 52 positions are noted as spheres. D. Average RMSF across three simulations for each model is shown, with the α2-α3 region that undergoes local unfolding highlighted in yellow. E. Close-up view of highlighted region in D. Residue 52 is noted with black dashed line. F. Secondary-structure probability profiles, created using *Weblogo 3*^108^, are shown as determined by Stride^109^ (H: α-helix, red, C: coil, blue, T: turn, purple, G: 3_10_-helix, orange). The secondary structure observed in the crystal structure is noted above.

We first assessed the effects of the above changes at the Cys52 position on the zinc-binding core. Our simulations revealed a loss of coordination between residue 52 and the zinc atom within the Cys-SH, Cys-SOH, and C52A simulations, while the pair maintain tight interactions throughout the C52D and Cys-S^−^ simulations (Fig. 2B). This loss of coordination has two principal effects. First, it leads to increased zinc lability and release from the active site as demonstrated by a corresponding increase in the distance between zinc and the zinc-binding core residue His83 (Fig. S3A-E). Second, it markedly affects the conformational flexibility of the entire α2-α3 segment (approximately residues 39-65) surrounding residue 52, in line with crystallographic and biochemical data suggesting α2-α3 becomes disordered upon disruptions to the zinc-thiolate interaction^12,23^. This effect is illustrated by the stark increase in average root mean square fluctuation (RMSF) of this region for Cys-SOH compared to Cys-S^−^ (Fig. 2C, Fig. S3F). Interestingly, the increased dynamics of the Cys52 position and α2-α3 segment are propagated to the N- and C-termini, which connect to the down and upstream-regulated protein domains, respectively (“signal transduction” arrows, Fig. 2C). Indeed, comparison of average RMSF across all systems reveals that the Cys-SH, C52A, and Cys-SOH states, which disrupt zinc-cysteine interactions, display increased global dynamics (Fig. 2D-E). Thus, changes at the zinc-binding core may be transduced to distant parts of the full-length homodimeric protein such as would be required for signaling in the case of diguanylate cyclases and chemoreceptors.

Consistent with the trends of RMSF, an analysis of average secondary structure probability of the α2-α3 region showed that more random coil is observed at the 3_10_ helix for C52A, Cys-SH, and Cys-SOH, reflecting their increased dynamics and propensity to undergo local unfolding, while less random coil is observed for Cys-S^−^ and C52D, reflecting those models’ stability and generally static nature (Fig. 2F). We note that the α3 region retained high helical content regardless of whether the region as a whole was fully folded or locally unfolded (Fig. 2F). This may help to explain why in our previous study HOCl-induced structural changes were observable by monitoring the fluorescence of the conserved α3 Trp (site 6, Fig. 1D), but not corresponding losses in helicity through circular dichroism at similar HOCl concentrations^12^ (Fig. 2D).

We recently used quantum mechanical (QM) methods to investigate the mechanism and origins of chemoselectivity of CZB domains with HOCl.^13^ Using the core of the CZB domain in *E. coli* DgcZ^23^ as a model system, we showed that the redox process occurs via direct OH transfer from HOCl to the zinc-bound cysteine model in a typical S_N_2 fashion, and that both reactivity and selectivity for HOCl over H_2_O_2_ are governed by minimizing active site geometric strain. To further understand how HOCl oxidation could promote CZB local unfolding and signal transduction, we have now performed additional QM calculations quantifying the energetic tendency for the bound cysteine model to be displaced from the CZB domain by either water or the HOCl ligand. We examined the ligand exchange equilibria at three redox and protonation sulfur states, namely CH_3_S(H), CH_3_SO(H), and CH_3_SO_2_(H) (Table 1).

**Table 1.** Ligand exchange equilibria depicting the energetic tendency for the sulfur ligand across protonation and oxidation states to dissociate from a model Zn^2+^ complex.

| **X** | $\Delta G_{\text{lig.ex}}^a$ | | **XH** | $\Delta G_{\text{lig.ex}}^a$ | |
| --- | --- | --- | --- | --- | --- |
|  | **YH = H<sub>2</sub>O** | **YH = HOCl** |  | **YH = H<sub>2</sub>O** | **YH = HOCl** |
| $\text{CH}_3\text{S}^-$ | 16.9 | 7.2 | $\text{CH}_3\text{SH}$ | 1.6 | -0.3 |
| $\text{CH}_3\text{SO}^-$ | 3.6 | -2.5 | $\text{CH}_3\text{SOH}$ | 1.0 | -0.4 |
| $\text{CH}_3\text{SO}_2^-$ | 9.1 | 8.7 | $\text{CH}_3\text{SO}_2\text{H}$ | -4.2 | -5.8 |

<sup>a</sup> Energies were calculated at 298 K and 1.0 atm and reported in kcal/mol.

Consistent with results from our MD simulations, our QM calculations indicate that in all but one scenario, the protonated sulfur states are more likely to be displaced from the zinc than the deprotonated states. Based on approximate pKa’s of Cys-SH (pKa ∼8.6)^50^, Cys-SOH (pKa ∼6.3-12.5)^51,52^, and Cys-SO_2_H (pKa ∼1.8)^53^, we suggest the following model at neutral cytosolic pH for the relative tendency of discrete cysteine redox states to be displaced from the zinc complex. Prior to oxidation, the deprotonated Cys-S^−^ state is energetically favored to remain associated with the zinc complex (ΔG for displacing CH_3_S^−^ = 16.9 kcal/mol by H_2_O and 7.2 kcal/mol by HOCl). However, upon HOCl oxidation, the protonated Cys-SOH state is more readily displaced from the complex (ΔG for displacing CH_3_SOH = 1.4 kcal/mol by H_2_O and –0.4 kcal/mol by HOCl), and even the deprotonated Cys-SO^−^ state is destabilized compared to the unreacted Cys-S^−^ state (ΔG for displacing CH_3_SO^−^ = 3.6 kcal/mol by H_2_O and –2.5 kcal/mol by HOCl). Following over-oxidization, the Cys-SO_2_^−^ protonation state predominates and is less favored than the Cys-SO^−^ state to dissociate from the complex (ΔG for displacing CH_3_SO_2_^−^ = 9.1 kcal/mol by H_2_O and 8.7 kcal/mol by HOCl).

Taken together, results of the MD and QM analyses indicate the unreacted thiolate plays an important role in protein folding and stability through interactions with zinc, and perturbations to the Cys-Zn bond including protonation, cysteine sulfenic acid formation, or deletion of the Sγ by mutation to alanine, promote local unfolding events (Fig. 2B). These molecular models are consistent with our previous experimental data and proposed mechanism that HOCl oxidation to Cys-SOH promotes a conformational shift of the α2-α3 region^12^. These computational analyses also provide several new insights into CZB conformation change and signal transduction. First, as mentioned above, the structural perturbations induced by alterations to Cys52 stimulate large increases in the dynamics of the termini that link to other protein domains, providing an explanation for how cysteine oxidation by HOCl could promote distant structural changes in diguanylate cyclase c-di-GMP catalysis and chemoreceptor organization^12^. Second, the conformational shifts stimulated by oxidation to Cys-SOH may lower the zinc-binding core’s affinity for zinc and shift the zinc-binding equilibrium toward the zinc-free state, and this behavior is mimicked by a C52A mutant. Thus, we hypothesized this could be a molecular mechanism linking CZB sensing of zinc and HOCl. Indeed, it was shown in earlier work that the C52A mutation in DgcZ stimulates increased α2-α3 disorder in the crystal structure and lowers zinc affinity by 10-fold, though this mutant still had femtomolar zinc affinity and retained the zinc in the crystal structure^23^. Reconstituted chemotaxis signaling assays also showed a similar response for the equivalent mutation in *H. pylori* TlpD and HOCl-treated wild type^12^. Third, by some metrics the C52D mutation was stabilizing to the zinc core, suggesting it could have increased zinc affinity over the native unreacted thiolate form. Earlier work by mass spectrometry showed that, in the absence of reductant, the cysteine is sensitive to overoxidation by HOCl to Cys-SO_2_^− 12^. The C52D mutant, therefore, can also be thought of as a mimic for that oxidation state, which might occur at high HOCl concentrations. Isolating the effects of discrete cysteine oxidation states is experimentally challenging due to the transient nature of reaction intermediates and mixture of species that can occur^31,54^, especially when attempting to learn their roles in signaling events *in vivo*. We recognized the utility of having CZB protein mutants as molecular probes that could mimic these oxidation states and proceeded with characterizing these mutants biochemically.

### DgcZ is selectively reactive with HOCl

The biological distribution of CZB architectures shows that many bacterial species, including some virulent pathogens, possess CZB-regulated diguanylate cyclases that could control c-di-GMP signaling and biofilm formation in response to exogenous HOCl produced through inflammation (Fig. 1, Table 2). To address this possibility, biochemical analyses were conducted with *E. coli* DgcZ to determine if the protein could facilitate HOCl-sensing (Fig. 3A). The reactivity of full-length DgcZ from *E. coli* with various concentrations of HOCl was determined through dimedone adduction and slot blotting analysis to monitor formation of cysteine sulfenic acid (Fig. 3B). The wild type protein, which contains five cysteines, exhibits strong reactivity toward micromolar HOCl, whereas mutation of the conserved Cys52 to Asp (C52D) reduced apparent Cys-SOH formation by approximately 5.7-fold (Fig. 3B). Further characterization using the CZB domain alone recapitulated this behavior, with 10 μM of the wild type protein being half-maximally oxidized by 122 μM HOCl, and the C52D mutant showing little oxidation even at 500 μM HOCl (Fig. 3C). Using HOCl concentrations above 500 μM reduced the amount of detected Cys-SOH, which we interpret as overoxidation of Cys52 to Cys-SO_2_^−^, as was observed to occur for *H. pylori* TlpD at similar concentrations^12^. In contrast, treatment with equivalent concentrations of H_2_O_2_ produced virtually no Cys-SOH, showing that Cys52 possesses the ability to discriminate between H_2_O_2_ and HOCl (Fig. 3D). Pre-oxidized CZB protein was reduced through addition of glutathione disulfide, consistent with the reversibility of the Cys-SOH formation (Fig. 3E). These data show that DgcZ has the capacity to react selectively with HOCl through oxidation at Cys52, can differentiate between HOCl and H_2_O_2_, and can be reduced by reductant systems present in bacteria.

**Fig. 3.**
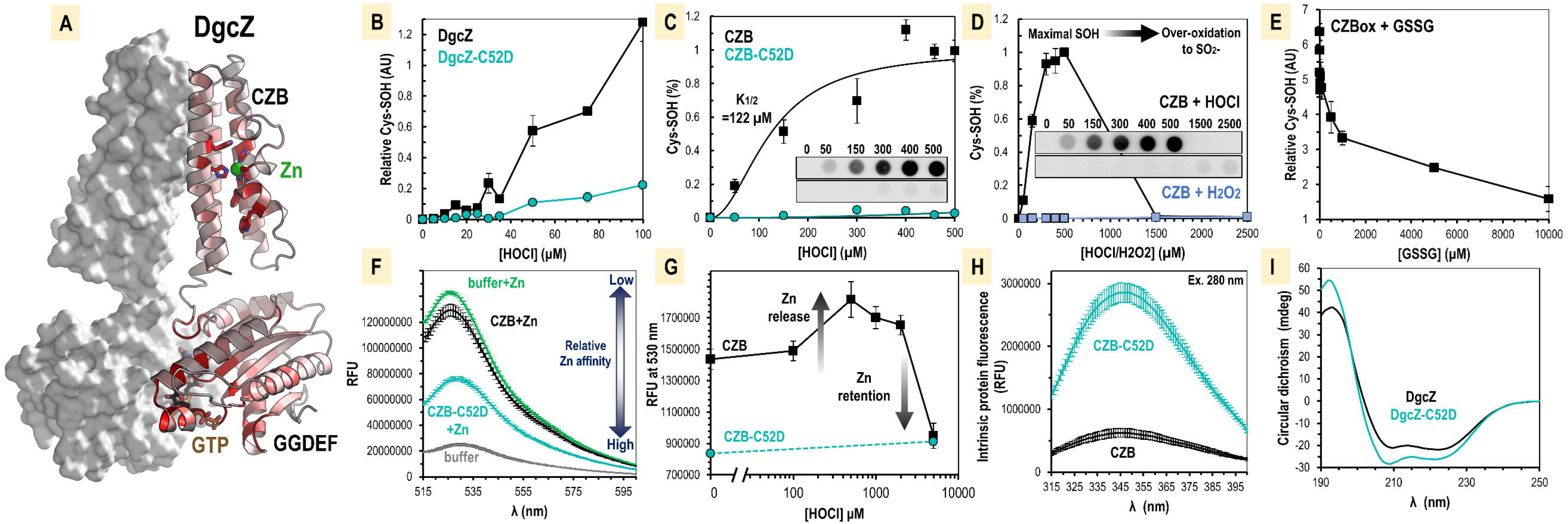
Oxidation of *E. coli* DgcZ by HOCl and influence on protein zinc ligation. A. The DgcZ homodimer is shown, with each monomer containing an N-terminal GGDEF domain, which catalyzes c-di-GMP from GTP, and C-terminal CZB domain that allosterically regulates the GGDEF domain through zinc-mediated inhibition^23^. To highlight regions of functional importance, amino acid conservation across all DgcZ-like homologues containing similar architecture is depicted, colored by conservation as in Fig. 1C. B. Cys52 is required for efficient reactivity with HOCl for full-length DgcZ (wild type shown in black, C52D mutant in teal, n=3). Reactions were performed in PBS at pH 7 with 10 μM protein. C. The CZB domain alone is sufficient to recapitulate Cys52-mediated HOCl-reactivity. Representative western blots are shown for 10 μM wild type CZB (top) and CZB-C52D (bottom), n=8. A fit of the Hill equation to the wild type (black line) and C52D (teal line) data with a Hill coefficient of 2 is shown, and used to approximate the concentration of HOCl required for half-maximal Cys-SOH formation (K_1/2_). D. The CZB domain (10 μM) is preferentially reactive with HOCl over H_2_O_2_ and Cys52 can be over-oxidized to form Cys-SO_2_^−^, n=3. E. Oxidized CZB protein (10 μM) pre-treated with 250 μM HOCl can be reduced by glutathione disulfide, n= 7-8. F. Emission spectra showing competition for zinc between zinpyr-1 probe and CZB proteins, n=3. Reactions contained 10 µM probe, 1 µM protein, +/- 5 µM ZnSO_4_ in PBS buffer pH 7. Zinpyr-1 emission spectra were obtained by excitation at 488 nm. G. Emissions spectra showing Zn^++^ release or retention is regulated by CZB oxidation state in the presence of zinpyr-1. Reactions contained 10 µM protein in PBS pH 7, n=3. Samples were treated with the indicated concentrations of HOCl, then quenched with 1 mM methionine, followed by addition of 10 µM probe. H. Intrinsic fluorescence spectra of 10 μM wild type CZB and CZB-C52D in PBS, n=3, excitation at 280 nm. I. Circular dichroism spectra for 5 µM DgcZ wild type and DgcZ-C52D in PBS. Data points shown are the averages across replicate samples, and error bars are standard error of the mean.

**Table 2.**
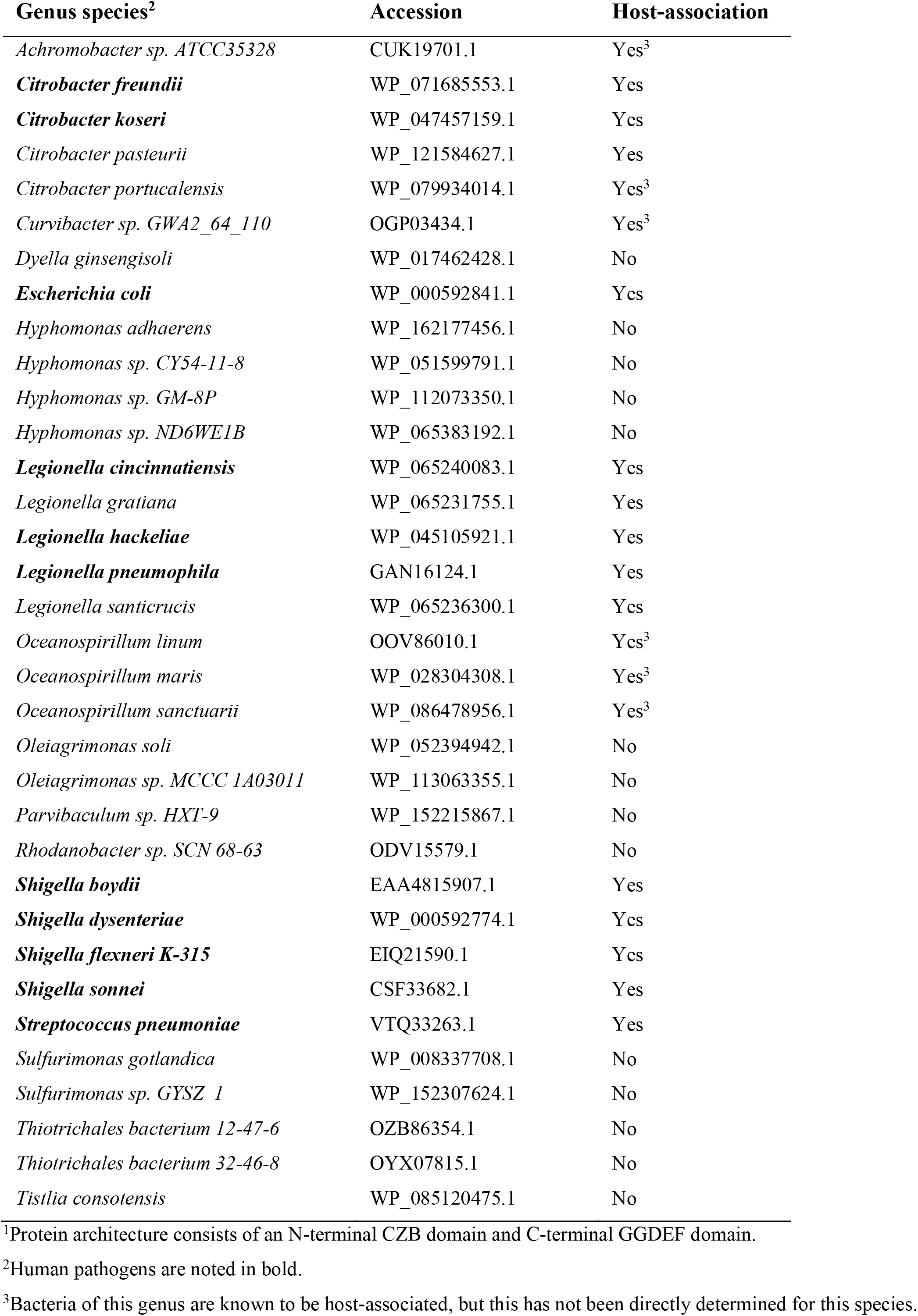
Bacteria that Possess DgcZ-like Proteins

### HOCl oxidation promotes zinc release

In other systems involving Cys-Zn coordination, cysteine oxidation by HOCl is known to promote zinc release^55,56^. Whether this occurs for CZB proteins remains an important open question relevant to understanding the domain’s signaling functions. Due to the protein’s high zinc affinity, even large changes to zinc affinity would not substantially alter the zinc-bound ↔ zinc-free equilibrium in the absence of zinc-binding competitors. Therefore, reactions to test CZB zinc lability were performed in PBS buffer, which contributes to zinc-chelation^30^, and monitored with the fluorescent zinc probe zinpyr-1, which has a relatively high affinity for Zn^++^ and displays an increase in fluorescence when zinc-bound^12,57^. To assay relative zinc affinity, we competed the wild type and mutant proteins against the zinpyr-1 probe in the presence of exogenous zinc and observed that wild type competed poorly and only slightly diminished the zinc available to the probe, while the C52D mutant reduced the probe’s fluorescence signal by about half (Fig. 3F). We next assayed whether HOCl treatment alters CZB zinc binding. HOCl treatment of the wild type protein showed a bimodal response; zinc was increasingly liberated (made available for the probe to bind) by increasing concentrations of HOCl up to 500 μM (50-fold molar HOCl/DgcZ ratio), and higher HOCl concentrations decreased the amount of zinc available to the probe (Fig. 3G). In contrast, the C52D mutant was unresponsive to HOCl and relinquished little zinc even when treated with millimolar HOCl (Fig. 3G). At 2.5 mM HOCl, the wild type retention of zinc was approximately equal to that of the C52D mutant (Fig. 3G). Based on the concentrations of HOCl that result in Cys-SOH and Cys-SO_2_-oxidation states for the wild type protein (Fig. 3C), we interpret these data to indicate that the Cys-SOH state decreases zinc affinity, and the Cys-SO_2_^−^ has increased zinc affinity, and that C52D mimics the increased zinc affinity of the Cys-SO_2_^−^ state (Fig. 3G). Additional biophysical characterizations comparing wild type and C52D forms are consistent with this idea. The C52D mutant has both a stronger alpha-helical circular dichroism signature and higher intrinsic fluorescence, supporting that it adopts less of the locally-unfolded conformation (Fig. 3H-I). Taken together these experimental results support the predictions from molecular modeling that (1) oxidation to Cys-SOH lowers zinc affinity and overoxidation to Cys-SO_2_^−^ increases zinc affinity, and (2) at the molecular level these discrete CZB redox states can be modeled/mimicked by C52A and C52D mutations, respectively.

### HOCl relieves Zn-mediated inhibition of DgcZ diguanylate cyclase activity

Diguanylate cyclases require dimerization for activity, i.e. the productive encounter of two GTP loaded GGDEF domains^58^. The CZB domain of DgcZ is responsible for constitutive dimer formation (Fig. 1C), but has also been shown to allosterically inhibit DgcZ activity when complexed with Zn^++ 23^. Titration of ZnCl_2_ results in a linear decrease of enzymatic activity as measured by the concentration of the c-di-GMP product after a defined incubation time (Fig. 4A). This is consistent with high Zn^++^ affinity, which for DgcZ has been reported to be in the sub-femtomolar range^23^. Note that a super-stoichimetric amount of ZnCl_2_ is needed for complete inhibition, which is most likely due to the loss of a considerable amount of Zn^++^ by complexation with phosphate from the buffer. Phosphate, as opposed to tris, was used to have an inert buffer in the follow-up HOCl experiments.

**Fig. 4.**
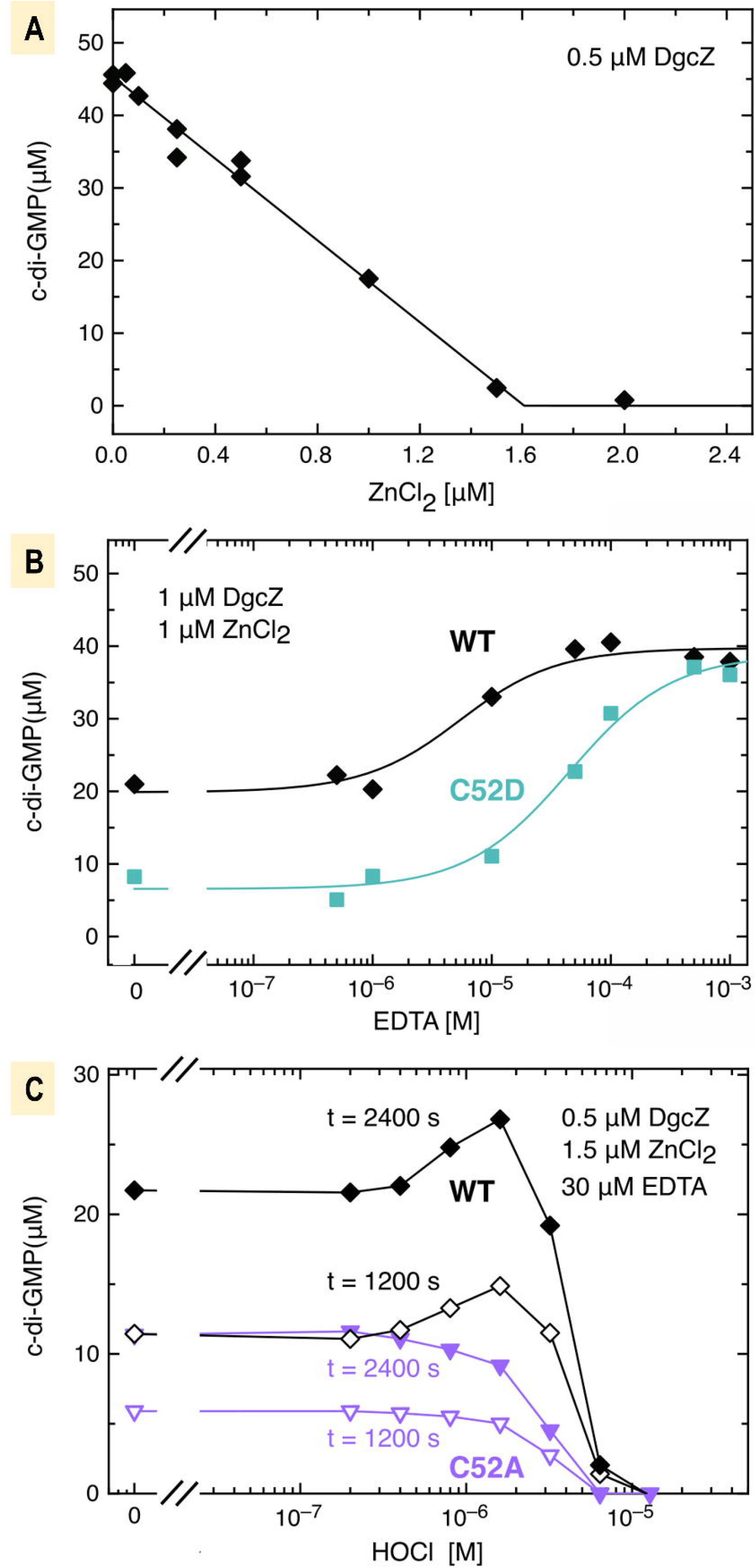
Zinc mediated inhibition of DgcZ activity is counteracted by HOCl. DgcZ catalyzed c-di-GMP synthesis was assayed by FPLC nucleotide concentration determination at saturating GTP (500 μM for panels A-B and 300 μM for C) and indicated enzyme concentrations. Incubation times were 1800 s and 1037 s, for A and B, respectively. A. Zinc titration of wild-type DgcZ demonstrating linear decrease of c-di-GMP production (solid line) consistent with high-affinity binding of the zinc inhibitor. Incubation time 1800 s. Note that more than a stoichiometry amount of ZnCl_2_ is required to fully inhibit DgcZ, because most likely not all Zn^++^ is available for binding due to complexation with the phosphate buffer. B. EDTA titration relieves zinc mediated enzyme inhibition. Solid lines represent the fit with a two-state model. For the C52D mutant about a 10-fold larger EDTA concentration is required for the half-maximal effect indicating tighter zinc binding of the mutant compared to wild type. C. HOCl titration in presence of EDTA competitor shows relief of DgcZ inhibition at low HOCl concentration (≤ 2 μM) for the wild type, but not for the C52A mutant. The C52A mutant demonstrates the effect of dose-dependent inactivation, seen also for the wild type at larger HOCl concentrations. Incubation times were as indicated.

As has been shown before^23^, zinc-mediated inhibition can be relieved by addition of a zinc chelator like EDTA that will compete for Zn^++^. Fig. 4B shows such a titration with EDTA for partially zinc-loaded DgcZ, where the wild type protein in the presence of Zn^++^ is activated by increasing concentrations of EDTA, which can be fit well with a two-state model. Under the same conditions the C52D mutant shows considerably lower activity without EDTA, but converges to the same maximal value at high EDTA (Fig. 4B). Again, the data conform to a two-state model, but with an inflection point at approximately 10-fold higher concentration. These data are in line with the increased Zn^++^ affinity of C52D compared to DgcZ wild-type as deduced above from the theoretical and biophysical results (Table 1, Fig. 3F-I, Fig. S3E, Movie S5).

Pruning C52 of DgcZ by mutagenesis (C52A) results in a moderate decrease of zinc affinity by about one order of magnitude^23^ and a similar effect can be expected for the Cys-SOH state of C52. Thus, in the absence of other zinc competitors C52 modification cannot be expected to alter significantly the complexation state of the CZB domain and, thereby, enzyme activity. Therefore, and also to mimic the presence of high affinity competitors of zinc like occurs *in vivo*^30^, we performed the HOCl titration experiment in the presence of EDTA at conditions where the available Zn^++^ was estimated to be shared about equally between DgcZ and EDTA. HOCl titration clearly increased DgcZ activity (as measured by c-di-GMP production after 20 and 40 min incubation) for HOCl ≤ 2 μM (4-fold molar HOCl/DgcZ ratio), whereas larger concentrations induced a negative effect (Fig. 4C). Comparison with the corresponding results for mutant C52A allowed us to attribute the activating effect to oxidation of C52. At larger HOCl concentrations, DgcZ activity decreases for both wild type and C52A proteins, which we interpret to be due to non-specific protein oxidation. In summary, in the presence of a strong zinc competitor, a clear C52-specific effect of DgcZ activation by HOCl has been demonstrated.

### DgcZ regulates biofilm in response to exogenous HOCl

To test the role of DgcZ in regulating bacterial biofilm, *E. coli* biofilms were grown and quantified under various conditions. For these assays we utilized a previously-engineered *csrA*-deletion strain of MG1655 that mimics *in vivo* DgcZ expression and biofilm formation when in a host under laboratory conditions^23^ (for clarity, we refer to this strain as “wild type” for the remainder of the text). In this background, we first compared the biofilm formation of strains expressing or lacking *dgcZ* (*dgcZ* or *ΔdgcZ*, respectively), in a static microplate when treated with increasing concentrations of HOCl (Fig. 5A). Crystal violet staining was used to quantify biofilm as done previously^23^, with relative biofilm calculated as a ratio of each sample divided by the average of untreated wild type in the same experiment. No difference in biofilm was observed after 24 H for controls containing untreated cells, water-treated, or treatments with PBS buffer at pH 7 (Fig. 5A). Single applications of HOCl diluted in PBS buffer showed a bimodal response for wild type cells, with relative biofilm increasing in response to micromolar HOCl, to a maximum of 1.6-fold at 250 μM, and decreasing at higher HOCl concentrations (Fig. 5A). The *ΔdgcZ* mutant displayed a different trend, showing decreased biofilm in the range of 5-250 μM HOCl, suggesting DgcZ activity may in fact overcome competing and opposite signaling factors in response to HOCl (Fig. 5A). Under these conditions both wild type and *ΔdgcZ* cultures were observed to grow equivalently and did not display growth inhibition (Fig. 5B). Experiments with more established cultures with cells at OD_600_ = 1.0 showed a similar, but smaller, *dgcZ*-dependent increase in biofilm in response to a single HOCl treatment, as well as biofilm inhibition with exogenous zinc treatment, as was previously reported^23^ (Fig. S4A).

**Fig. 5.**
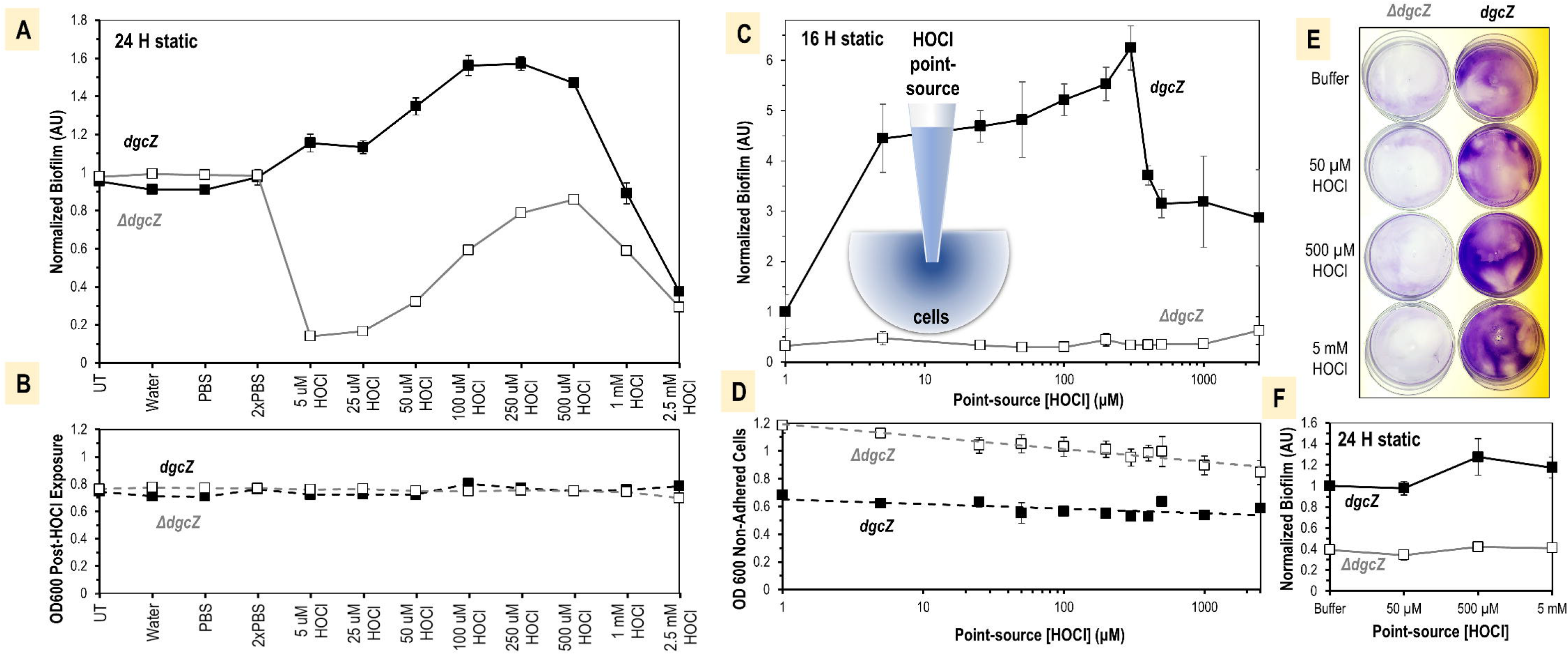
DgcZ regulates biofilm formation in response to HOCl. A. Biofilm after 24 H static growth at 30° C following exposure to control and HOCl treatments. Cell cultures were grown in a 96-well microplate with an initial OD_600_ of 0.5. B. Final OD_600_ of cultures from panel A. C. Biofilm after 16 H at 25° C with exposure to pipettes containing various concentrations of HOCl point sources. A logarithmic scale is shown and for clarity “0” HOCl (treatment with PBS) is shown as 1 µM. D. Final OD_600_ of planktonic fraction of cultures in panel C. E. Representative results for 24 H static biofilm assays with 3 ml of cell culture at 25° C with a central point source treatment, following staining of adhered cells with crystal violet. F. Quantification of biofilm in panel E. For all data shown, points indicate the sample mean, and error bars are standard error of the mean.

Whereas single treatments of HOCl might react quickly and dissipate, we performed a similar version of this static biofilm microplate assay with treatment “point-sources” to model a more sustained exposure to an HOCl microgradient. This was accomplished by using a 96-well Rainin liquidator with pipettes containing HOCl treatments over the range of 5-2500 μM that were submerged in cell cultures for 16-24 H (Fig. 5C). These data showed a DgcZ-dependent maximal increase in relative biofilm of 6.25-fold at 300 μM (Fig. 5C). For these cultures, the OD_600_ of the planktonic population was lower for the wild type than the *ΔdgcZ* mutant, possibly owing to the higher fraction of surface-attached cells for the wild type, and a slight negative correlation was observed between cell density and HOCl concentration (Fig. 5D). We further scaled up these experiments in 15 mm petri dishes, and 3 ml of cell culture, with a central point source treatment and visualized biofilm by crystal violet staining after 24 H. These assays suggested biofilm formation distribution may be altered by the HOCl treatment point sources (Fig. 5E). The total biofilm followed a similar response to HOCl in these assays as experiments with smaller cell volumes (Fig. 5F).

The high conservation of the CZB zinc-binding cysteine has been previously documented^12,23,59^, but its importance has never been investigated *in vivo*. Therefore, we created two new *E. coli* strains through CRISPR gene editing that contain chromosomal point mutants to express *dgcZ^C52A^* or *dgcZ^C52D^* under native promotion in order to mimic oxidized versions of DgcZ and directly test whether the conserved cysteine is important for biofilm formation. As DgcZ is known to regulate poly-n-acetylglucosamine (poly-GlcNAc)-dependent biofilm formation^23,38^, we assessed poly-GlcNAc production through growth on LB agar plates containing the poly-GlcNAc-binding dye Congo red, including a strain expressing an enzymatically-inactivated *dgcZ^E208Q^* mutant^23^ (Fig. 6A). Under these conditions we found that the wild type exhibited a modest amount of Congo red binding, the *dgcZ^C52A^* and *dgcZ^C52D^* mutants showed increased binding, and the *ΔdgcZ* and *dgcZ^E208Q^* mutants showed low dye binding (Fig. 6A). To determine relative poly-GlcNAc production in these assays, images of colonies were color-thresholded and Congo red binding quantified as red pixels (dye-containing cells) as a fraction of red + brown (non-dye containing cells), normalized to the wild type strain (Fig. 6B). For plates inoculated with cells from mid-log exponential cultures the *dgcZ^C52A^* and *dgcZ^C52D^* mutants showed 4.4 and 7.0-fold greater dye binding, respectively, than wild type (Fig. 6B, left column). Plates inoculated with cells from established overnight cultures also showed elevated Congo red binding by *dgcZ^C52A^* and *dgcZ^C52D^* mutants, 2.1 and 2.4-fold higher than wild type, respectively, with wild type showing a moderate amount, and *ΔdgcZ* and *dgcZ^E208Q^* mutants showing lowered binding (Fig. 6B, right column).

**Fig. 6.**
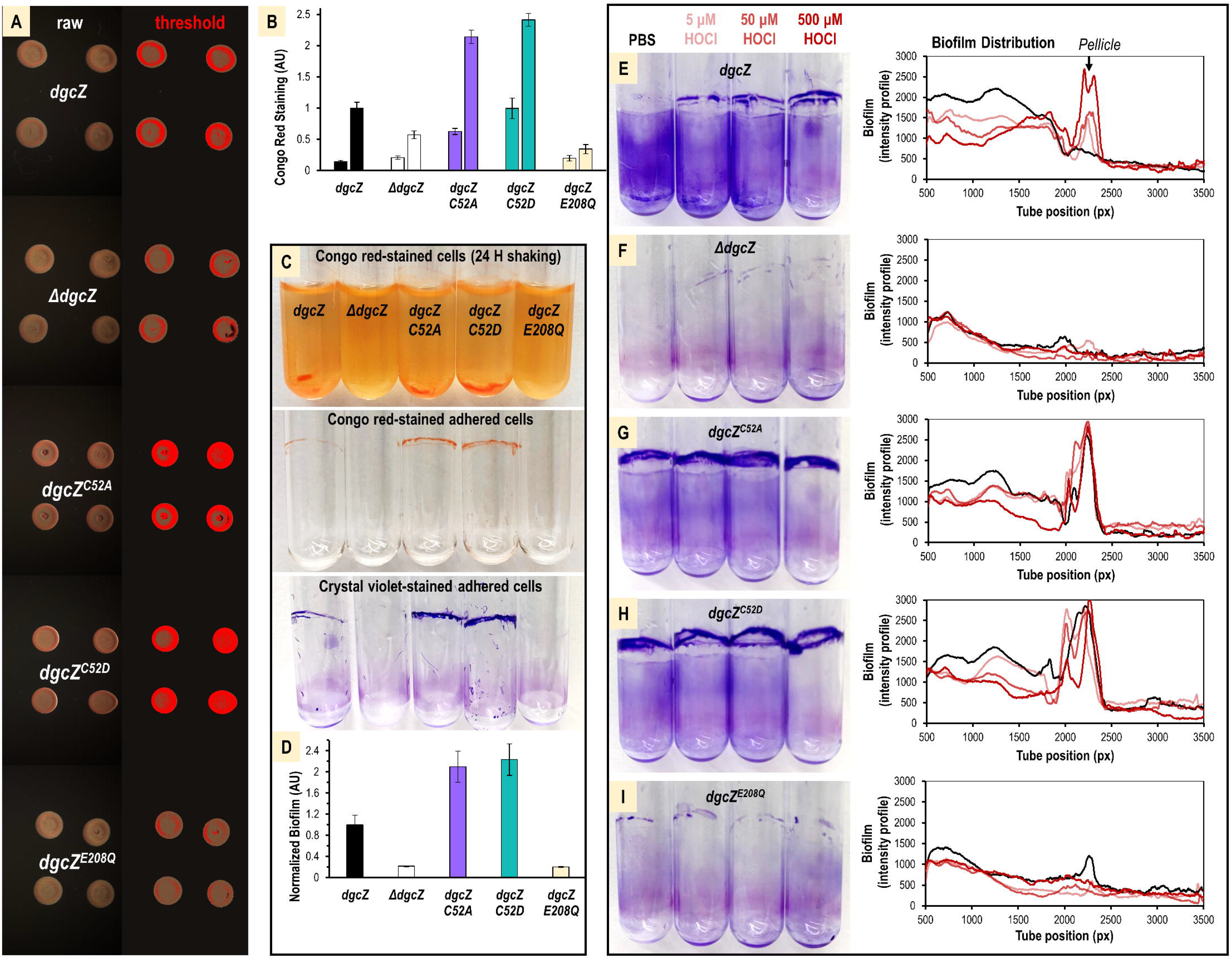
The conserved Cys52 mediates HOCl-sensing and biofilm distribution. A. Representative images showing growth of cells expressing functional DgcZ (*dgcZ*), lacking (*ΔdgcZ*), containing cysteine point mutants (*dgcZ^C52A^*, *dgcZ^C52D^*), or a form unable to catalyze c-di-GMP (*dgcZ^E208Q^*) on Congo red-LB agar plates after 24 H at 25° C. Plates were inoculated with 4 x 2 μl of liquid cultures grown overnight in LB. Raw images are shown on the left, and the color-thresholded versions are shown on the right. Relative dye uptake is visualized by clustering pixels into red (dye-containing cells), brown (non-dye-containing cells), and black (background). B. Quantification of Congo red dye uptake from experiments in A (right column) and equivalent experiments inoculated with mid-log exponential cells at OD_600_ 0.5 (left column), n=12. C. Comparison of Congo red dye uptake and biofilm formation after 24 H static growth at 30° C. D. Quantification of biofilm experiments from C by crystal violet staining, n=10. E-I. Biofilm distribution and pellicle formation is shown for cells treated with PBS (black), 5 μM HOCl (light red), 50 μM HOCl (medium red), or 500 μM HOCl (dark red) and grown for 24 H shaking at 30° C. Representative images are shown of crystal violet-stained tubes on the left, and biofilm density along the vertical length of the tubes are quantified as an intensity profile on the right, n=3.

Based on *in vitro* analyses, we predicted *dgcZ^C52A^* and *dgcZ^C52D^* mutations might stimulate greater and lowered biofilm formation, respectively, due to their differences in zinc-mediated inhibition (Fig. S3, Fig. 4B-C). However, we found when grown in static liquid cultures both *dgcZ^C52A^* and *dgcZ^C52D^* mutations increased biofilm formation (Fig. S4B). In a point-source assay, however, the behavior seen *in vitro* was somewhat recapitulated, with wild type biofilm formation increased by HOCl, *dgcZ^C52A^* unresponsive, and *dgcZ^C52D^* showing decreased biofilm (Fig. S4C).

Lastly, we assayed biofilm production for these *E. coli* strains in rocking liquid cultures. Similar to Congo red staining and biofilm formation in static assays, the wild type showed a moderate degree of poly-GlcNAc production and biofilm formation, the *ΔdgcZ* and *dgcZ^E208Q^* strains showed lower, and the *dgcZ^C52A^* and *dgcZ^C52D^* mutants were elevated (Fig. 6C-D). In these experiments we observed differences in biofilm at the liquid-solid interface (tube bottom) and the formation of a pellicle at the liquid-air interface^60,61^ between the wild type and cysteine mutant strains, suggesting this distribution could be regulated by the conserved C52 (Fig. 6C). Cultures of wild type cells treated with increasing concentrations of HOCl diluted in PBS buffer showed a dose-dependent increase in pellicle formation over PBS treatment alone (Fig. 6E). Equivalent experiments with *dgcZ^C52A^* and *dgcZ^C52D^* mutants showed robust pellicle formation regardless of HOCl treatments, and *ΔdgcZ* and *dgcZ^E208Q^* mutants exhibited low biofilm and little change in pellicle formation (Fig. 6F-I, Fig. S4D). Equivalent biofilm experiments with addition of exogenous zinc partially recapitulated *in vitro* observations of differences in zinc-mediated inhibition of DgcZ c-di-GMP of wild type and cysteine mutants, with addition of 50 μM zinc decreasing biofilm for wild type and *dgcZ^C52D^*, and the *dgcZ^C52A^* mutant less sensitive to zinc inhibition (Fig. S4E).

## DISCUSSION

Learning how bacteria sense and organize in response to host inflammation pathways will shed light on host colonization, diseases of intestinal dysbiosis, and the roles of bacterial pathogenicity in diseases of chronic inflammation. Despite HOCl being the strongest and most abundant reactive oxygen species generated by the neutrophilic oxidative burst during inflammation^1,3,4,62^, relatively little is known about the signaling networks bacteria use to perceive and respond to the presence of this oxidant in their environment^14,63^. Here, we identify bacterial CZB proteins as c-di-GMP and biofilm regulators that optimize bacterial behaviors in response to HOCl, implicating them as important players in bacterial pathogenicity and diseases of chronic inflammation.

### New insights into the molecular mechanism of CZB HOCl-sensing and signal transduction

In this study we leveraged the CZB-containing protein DgcZ from *E. coli* as a model system to obtain broad insight into the molecular mechanism of CZB signaling. Earlier work had suggested reactivity toward HOCl may be a common feature of CZB domains, as representative proteins from *Helicobacter*, *Salmonella*, and *Escherichia* all showed similar propensity for oxidation^12^. However, the HOCl-sensing mechanism had only been investigated in the context of a single non-canonical cytosolic chemoreceptor TlpD from *H. pylori*^12^, and so the degree to which these findings may be relevant to other bacteria, processes outside of chemotaxis, and other CZB subgroups, was unknown. Our biochemical analysis of DgcZ HOCl-sensing supports that CZB-regulated diguanylate cyclases are reactive with physiological concentrations of HOCl through a conserved Cys-Zn redox switch (Fig. 3B-D). Moreover, the CZB redox switch exhibits a high degree of specificity for reaction with HOCl over H_2_O_2_, approximately 685-fold (Fig. 3D), can be reduced by the GSH/GSSG system (Fig. 3E), and oxidation to Cys-SOH results in the relief of the Zn-mediated inhibition of diguanylate cyclase activity (Fig. 4C). The elevated c-di-GMP production activates c-di-GMP signaling pathways controlling aggregation, surface-attachment, and biofilm formation (Fig. 5, Fig. 6) to initiate the transition of bacterial lifestyle from planktonic to sessile.

Our joint experimental and computational analyses of the impacts of HOCl oxidation on zinc affinity give the first insight into the relationship between these two ligands in CZB regulation, and links the mechanisms proposed for *H. pylori* TlpD HOCl-sensing^12^ and *E. coli* DgcZ zinc-sensing^23^. The redox state of the conserved cysteine alters the zinc-binding core’s affinity for zinc, and, when in the presence of other high affinity zinc competitors, as occurs in the cell cytosol, can sufficiently shift the zinc-binding equilibrium to regulate the domain’s structure and dynamics. Specifically, oxidation by a single HOCl molecule to cysteine sulfenic acid lowers zinc affinity, and oxidation by a second HOCl to cysteine sulfinate increases zinc affinity, and these behaviors are well-modeled *in vitro* by C52A^23^ and C52D mutants, which show approximately 10-fold decreases and increases in zinc affinity, respectively (Fig. 3F-G, Fig. 4). The increased zinc affinity exhibited by the Cys-SO_2_^−^ state when treated with high concentrations of HOCl, and mimicry by the C52D mutant (Fig. 3F-G), could relate to an inhibition of DgcZ activity seen at higher HOCl concentrations (Fig. 4C), and decreased biofilm in excess of >1 mM (Fig. 5A). However, it is generally thought for bacteria that Cys-SO_2_^−^ is not a modification used for signaling, and its formation is avoided, because bacteria lack the enzyme sulfiredoxin and the ability to reduce and recover this cysteine redox state^6^. Our *in vitro* studies on Cys-SOH (and C52A) may be most relevant for understanding bacterial responses to physiological concentrations of HOCl, but it remains possible CZB Cys-SO_2_^−^ formation could contribute to CZB signaling in certain instances, such as environments with extremely inflamed tissue, and/or bacteria with poor cellular redox buffering capacity. Nevertheless, the C52D mutant served as a valuable tool *in vitro* for assaying the importance of the conserved C52 and learning the effects of increasing zinc affinity on protein activity.

The signal transduction of CZB domain-containing proteins appears to occur through some reorganization or change in dynamics of their obligate homodimers (Fig. 1B-C)^12,23^ that propagates between structurally-distant parts of the full-length protein. Previous crystallographic work suggested a model in which zinc-binding rigidifies the CZB dimer, and zinc-release increases structural flexibility and dynamics^23^. That the formation of cysteine sulfenic acid through oxidation with HOCl is destabilizing to the zinc-binding core is supported by our QM ligand isomerization calculations showing that displacement of the sulfenic acid is more thermodynamically favored than the thiolate state by approximately 15 kcal/mol, predicting that the added geometric strain shifts the equilibrium of the α2-α3 region from being fully folded toward locally unfolded (Table 1). Consistent with this, we have performed the first molecular dynamics modeling of CZB signal transduction and capture in these simulations a fascinating interplay between cysteine oxidation, local unfolding, zinc lability, and signal transduction (Fig. 2). In these models, we witness the cysteine sulfenic acid stimulate a dramatic increase in dynamics and propensity for local unfolding (Fig. 2B-C) that permits zinc release (Fig. S3), consistent with our *in vitro* data (Fig. 3F-G). Intriguingly, we see that local unfolding of α2-α3 removes packing interactions that stabilize α1 and α5 (Fig. 2C, Fig. S3F), the N- and C-termini, respectively, thereby increasing the dynamics of the CZB regions that connect to regulated protein domains (Fig. 1B). This link between the stabilization of the zinc-binding core and the dynamics of the CZB termini provide an explanation for how CZBs can use the same structural topology to regulate proteins like chemoreceptors, which mostly have C-terminal CZBs, and diguanylate cyclases that have N-terminal CZBs (Fig. 1A-B).

Together, we propose a new and refined model for HOCl-sensing by CZB domains that synthesizes these new findings about HOCl modulation of zinc affinity and protein dynamics and incorporates previous experimental results to provide a unifying molecular mechanism (Fig. 7). CZB domains bound to zinc are stable and relatively rigid, preventing structural flexibility of the proteins they regulate. The Cys-Zn redox switch is highly reactive and specific toward HOCl oxidation to form Cys-SOH, and the molecular signal is transduced through the release of zinc in the presence of Zn-binding competitors, which can be stimulated by Cys-SOH formation. Zinc release promotes local unfolding of the α2-α3 region and destabilization of the CZB termini, leading to greater structural flexibility of the full-length protein. The increased dynamics permit the population of conformations required for signal transduction; in the case of DgcZ-like proteins this allows the GGDEF domain to align productively for c-di-GMP catalysis^23^, and for TlpD-like chemoreceptors disorganization of the coiled-coil domain can inhibit the autophosphorylation activity of the histidine kinase chemotaxis protein A (CheA)^12^.

**Fig. 7.**
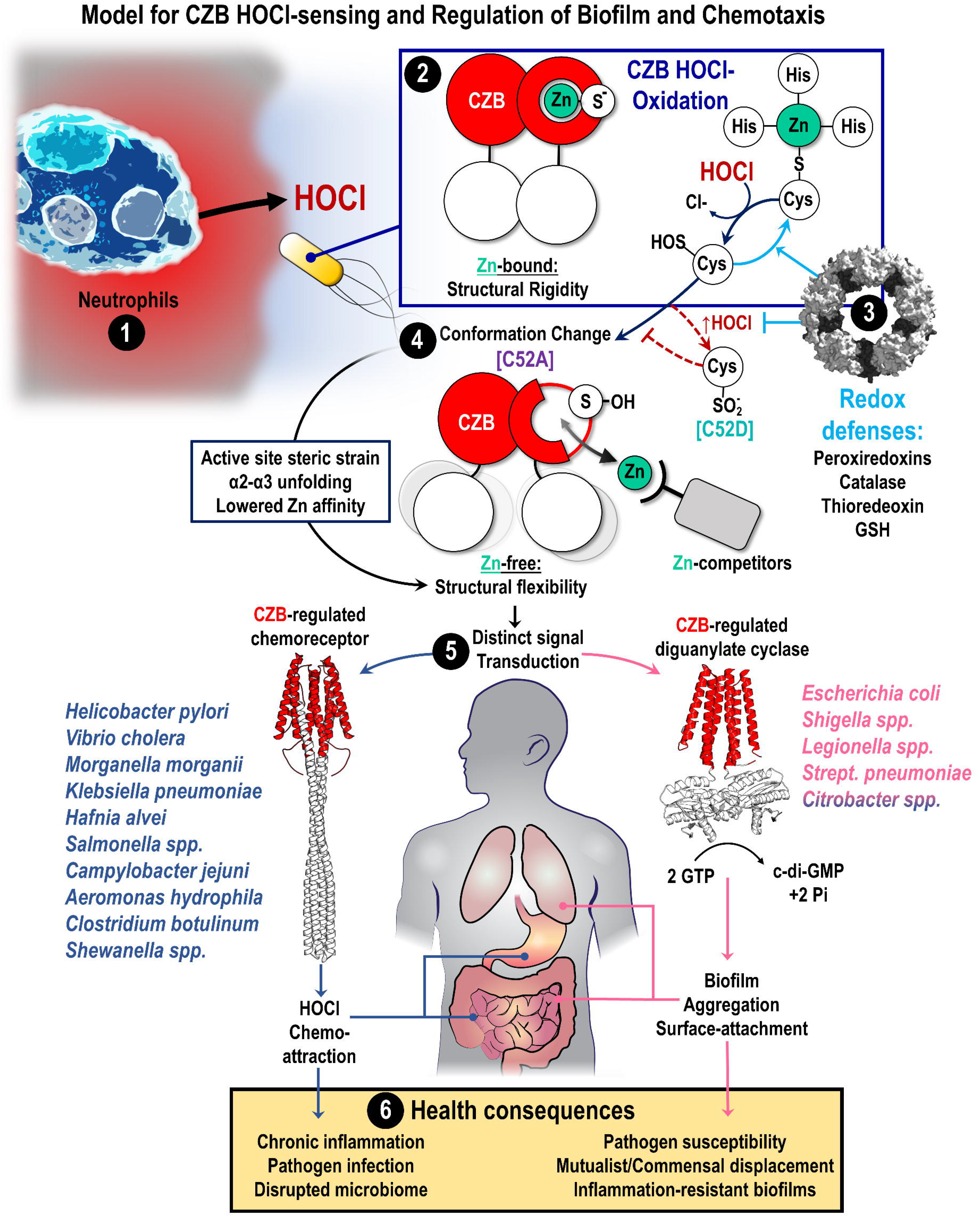
Proposed model for the role of CZB proteins in bacterial sensing of HOCl, and implications for bacterial pathogenesis. (1) Neutrophils produce HOCl as part of innate immunity and inflammation to control bacterial populations and combat pathogens. (2) Bacterial CZB domains (red circles) exist as homodimers that bind zinc (green circle) in the low to sub-femtomolar range. Zinc-binding promotes structural rigidity of the full-length proteins they regulate (white circles). CZB domains sense neutrophilic HOCl through the unique reactivity of their conserved zinc-thiolate complex and direct cysteine oxidation to form cysteine sulfenic acid (Cys-SOH). (3) CZB oxidation is reversed and inhibited by cellular reductants such as glutathione, and possibly antioxidant enzymes. (4) The formation of Cys-SOH, modeled by the C52A mutant (violet) drives a large conformational change in the CZB domain through active site strain induced by the Cys-SOH to promote local unfolding of the α2-α3 region, and this lowers the domain’s zinc-binding affinity. In the presence of cellular chelators that compete for zinc, this shifts the equilibrium toward the protein being zinc-free and promotes structural flexibility and increased dynamics. Alternatively, under conditions of high levels of HOCl (dark red dashed lines), the cysteine can react with a second molecule of HOCl to form cysteine sulfinate (Cys-SO_2_^−^), is modeled by the C52D mutant (teal), which has higher zinc affinity and inhibits signal transduction. (5) Bacterial pathogens and pathobionts typically possess either CZB-regulated chemoreceptors (TlpD-like, blue) or CZB-regulated diguanylate cyclases (DgcZ-like, pink) and thus HOCl-sensing is integrated into chemotaxis or c-di-GMP-signaling processes, respectively. (6) This can contribute to disease in a number of ways, such as initiating positive feedbacks loops that promote chronic inflammation though HOCl chemoattraction (ex: *H. pylori*), altering the tissue environment to the disadvantage of health-promoting native microbiota (ex: *S. enterica*), or initiating virulence and inflammation-resistance pathways in pathobiont communities (ex: *E. coli*).

### CZB domains regulate the switching of biofilm lifestyle in response to HOCl

Bacterial biofilms are well-known to play protective roles against the immune system^35^ and are associated with numerous bacterial diseases^64^, and here we provide evidence that CZB domains perform a biological function as a direct sensor of exogenous HOCl to regulate c-di-GMP signaling, poly-GlcNAc production, surface attachment, and biofilm distribution.

Our *E. coli* model biofilm system shows that DgcZ is required for increased biofilm formation *in vivo* in response to HOCl in the range of 5-500 μM, and that cell growth is not impaired under these conditions (Fig. 5). We also show high concentrations of HOCl (>1 mM) can inhibit biofilm formation, which may explain previous investigations that reported both increased^24^ and decreased^65,66^ *E. coli* biofilm formation in response to oxidative stress. However, directly linking this cellular response to our *in vitro* characterization of DgcZ activity required dissection of the importance of the conserved C52 *in vivo.* To this end, we engineered and utilized the first bacterial strains containing point mutations of the conserved CZB cysteine to verify whether the molecular mechanism of cysteine-mediated HOCl-sensing plays out in the complex process of bacterial c-di-GMP signaling. Guided by our biochemical and modeling data showing that cysteine oxidation states can be mimicked by C52A and C52D mutations to DgcZ, we characterized the biofilm formation and behavior of these mutant strains under various conditions. Indeed, these cysteine point mutants displayed 2-fold greater basal poly-GlcNAc production (Fig. 6A-B) and 1.5-2-fold higher biofilm formation (Fig. 6C-D, Fig. S4B) over the wild type. The inability of the *ΔdgcZ* deletion strain and the catalytically-inactivated *dgcZ^E208Q^* strain to recapitulate these responses suggests these biofilm differences are due to direct regulation of DgcZ c-di-GMP production rather than indirect regulation of c-di-GMP signaling through effector proteins (Fig. 5A-F, Fig. 6E-I, Fig. S4C).

Interestingly, there were some differences between our *in vitro* biochemical analyses of the protein DgcZ and the cellular role of DgcZ in biofilm assays. First, only a modest increase in catalytic activity in response to low micromolar HOCl was observed for DgcZ protein *in vitro* (Fig. 4), whereas 1.5-6.2-fold increases in DgcZ-dependent biofilm occurred in response to HOCl treatments (Fig. 5A-E). Such discrepancy between *in vitro* and *in vivo* responses are not unprecedented and can sometimes be attributed to feedbacks involving c-di-GMP effector proteins that are not present in the *in vitro* analyses^67^. Alternatively, cellular reductants may moderate non-specific oxidation of DgcZ and allow activation responses at concentrations of HOCl higher than what we observed *in vitro*. Second, although *in vitro* the C52D mutant DgcZ protein had 10-fold higher zinc affinity than wild type, and its c-di-GMP catalysis was more readily inhibited by zinc, *in vivo* this mutation performed similarly to *dgcZ^C52A^* in most assays and had increased basal poly-GlcNAc synthesis and biofilm formation (Fig. 6A-D, Fig. S4D). The exception was in the point source assay, in which continual HOCl exposure resulted in decreased biofilm for the *dgcZ^C52D^* strain (Fig. S4C). The explanation for these differences is not clear, but we note that cytosolic zinc-shuffling is complex and can effectively serve as a second messenger for many cellular processes including redox relays and metabolism^27,68^. Overall, the biological role of DgcZ in sensing exogenous Zn^++^ *in vivo* is unclear because zinc is normally present in only trace amounts and highly insoluble at neutral pH^27,30^.

Our work demonstrates that the CZB conserved cysteine is critical for regulating poly-GlcNAc production and determining biofilm distribution in response to HOCl. Treatment with HOCl for wild type *E. coli* show increased formation of pellicle biofilm, and the oxidized *dgcZ^C52A^* and *dgcZ^C52D^* mimics show robust pellicle formation with and without HOCl treatment (Fig. 6E-I, Fig. S4D). Although understanding of the roles of different biofilm organizations remains incomplete and may be species-specific, several previous studies have investigated the conditions that favor robust pellicle formation. Pellicle formation in *E. coli* is linked to the production of poly-GlcNAc^69^, which we see is increased in the *dgcZ^C52A^* and *dgcZ^C52D^* strains (Fig. 5A-C). A majority of clinical isolates of uropathogenic *E. coli* (UPEC) were found to promote virulence through production of poly-GlcNAc^70^, and robust pellicle formation is associated with enteropathogenic *E. coli* (EPEC)^60^. For *Salmonella enterica* Typhimurium, it was discovered that nutrient availability regulates the switching of biofilm distribution, with so-called “bottom” biofilm favored in low nutrient conditions and pellicle formation increased with high nutrient conditions^71^. That *E. coli* may perceive HOCl exposure as indicative of a nutrient-rich environment could relate to observations of metabolic advantages exploited by *E. coli* using nitrate for respiration in the inflamed gut^20^.

### Roles for bacterial CZB HOCl-Sensing in colonization and disease

The relationship between inflammation and bacterial colonization of the human gastrointestinal tract is central to many diseases, as innate immune responses that fail to clear pathogens can manifest into states of chronic inflammation, causing tissue damage and even carcinogenesis^1,4^. Such an example is seen for chronic *Helicobacter pylori* stomach infections and the development of gastric adenocarcinoma^72,73^. In other cases, the inflammation induced by a bacterial pathogen can displace the native microbiota and promote dysbiosis, which is thought to play a contributing role in the development of inflammatory bowel diseases such as Crohn’s disease and Ulcerative Colitis^19,74,75^. Thus, understanding how bacteria sense and respond to molecular cues from host inflammation will advance therapeutic strategies for diseases of chronic inflammation and gut dysbiosis (Fig. 7).

Bacterial populations of healthy human microbiomes that inhabit the low-oxygen environment of the gut are thought to be dominated by obligate anaerobic bacteria from phyla Bacteroidetes and Firmicutes, with Proteobacteria species being less abundant^76^. However in diseases of dysbiosis involving gut inflammation, such as colitis^77^ and Crohn’s disease^78^, a bloom in facultative aerobic Gammaproteobacteria is observed. Similar disturbances to microbiome communities have been shown to act through enteric Gammaproteobacteria pathogens that stimulate host inflammation as part of their infection strategies^19,79^. The HOCl landscapes of healthy and inflamed tissue are vastly different^80^. Neutrophils are the source of myeloperoxidase, and in their activated state use this enzyme to convert approximately 90 % of their available molecular oxygen to HOCl^81^. Thus, the influx of neutrophils in the inflamed gastrointestinal tract generates a dramatic shift in challenges by reactive oxidants and available nutrients that can favor the opportunist. It has been proposed that the expansion of Gammaproteobacteria species in these disease states is linked to their inherent ability to exploit host inflammation by utilizing HOCl-reaction products as nutrients^74^. Tetrathionate and nitrate are two such metabolites that have been identified^20,79,82,83^. Therefore, HOCl could be a signal of opportunity for microbes able to tolerate and exploit inflamed tissue, and CZB proteins are part of a network that detects and relays that signal through chemotaxis and c-di-GMP signaling (Fig. 7).

Indeed, we observe that established cultures of *E. coli* survive HOCl concentrations beyond that thought to be produced physiologically (Fig. 5B), and our previous work showed *H. pylori* remained motile even after treatment with millimolar HOCl^12^. Others have noted the ability of *E. coli* to form and sustain biofilms long-term in the presence of HOCl, as well as *Legionella pneumophila*, an opportunistic pathogen of the lungs and another species that possesses a DgcZ homologue (Table 2)^84,85^. Our phylogenetics analysis shows that CZB proteins are prevalent among Proteobacteria, with the class Gammaproteobacteria accounting for approximately a third of all known sequences (Fig. 1E). This class also shows the greatest diversity of CZB protein architecture, with CZB signaling regulating transmembrane and soluble chemoreceptors, CheW proteins that are components of chemotaxis signaling arrays, c-di-GMP synthesis and degradation, and CZB-only proteins with as-of-yet undetermined functions (Fig. 1F). We also observe that species most often possess either a CZB-regulated chemoreceptor, such as *H. pylori* and *S. enterica*, or a CZB-regulated diguanylate cyclase, such as *E. coli* or *Shigella sonnei*, but not both, indicating CZB domains from these two functional groups may utilize the same chemistry to sense HOCl but promote distinct biological responses (Fig. 1G, Fig. 7).

Adhesion of *E. coli* to intestinal microvilli is the primary mechanism of *E. coli*-induced diarrhea in humans^86^, and kills over 50,000 individuals annually^85^, and an emerging body of evidence suggests that enduring and manipulating host inflammation is a central aspect of *E. coli* pathogenicity. CZB sensing of HOCl and stimulation of biofilm processes could play important roles in enabling the bacterium to thrive in inflamed environments. For instance, the majority of urinary tract infections are caused by UPEC^87^, and a hallmark of the disease in patients is pyuria, stimulated through upregulation of the neutrophil chemokine interleukin-8^43^. Neutrophil infiltration of tissue has been proposed to benefit *E. coli* pathogenicity by stimulating adherence to epithelial cells, which our data would suggest is induced through CZB recognition of neutrophilic HOCl^41^. Relative to healthy individuals, Crohn’s disease colonoscopic biopsies show marked increases of bacteria, and over 50 % of these are *E. coli*^88^. Intriguingly, it has been reported that the anti-inflammatory drug sulfasalazine, used for treatment of ulcerative colitis, is able to bind and directly inhibit DgcZ and *E. coli* biofilm formation^89^, supporting that CZB-regulated proteins may be therapeutic targets for interfering with bacterial co-existence with chronic inflammation.

### CZB conservation, evolution, and origin

Our exhaustive survey of all known CZB sequences provides the first quantification for what proteins and systems are regulated by CZB domains and identifies unique conservation patterns related to functional differences (Fig. 1). We expand on previous observations of the wide biological distribution of CZB domains^59^ to quantitatively report they are found in 21 bacterial phyla and prevalent in Gammaproteobacteria. These include many human gastrointestinal pathogens such as species from the genera *Vibrio*, *Shewanella*, *Shigella*, *Helicobacter*, *Campylobacter*, *Citrobacter*, and *Salmonella*, and pathogens associated with nosocomial infections such as *Legionella*, *Morganella*, *Klebsiella,* and *Streptococcus* (Fig. 1E). The CZB domains of *H. pylori* TlpD and *E. coli* DgcZ share only 25 % sequence identity, yet have the same zinc-binding core and retain similar HOCl-sensing functions^12^. Conservation of the HOCl-sensing machinery suggests the ability to perform this chemistry is inherent to CZB domains and could have widespread utility in bacterial sensing of host inflammation cues.

Though we have focused on CZB sensing of HOCl due to its relevance to chronic inflammation and bacterial pathogenesis, we also confirm earlier work showing the ability to sense zinc^23^. Because we identify CZB protein domains in bacteria from diverse environments that are not host-associated (Fig. 1E), CZB proteins must also have additional functions outside of HOCl sensing. Our results indicate the core mechanism of CZB regulation is the structural rigidity conferred by binding zinc; any effector that alters the equilibrium between zinc-bound and zinc-free forms can induce a regulatory output. We present evidence here that HOCl is one such effector, and others could possibly include exogenous zinc^27^, other chemical species known to have reactivity with thiol-based redox switches such as aldehydes and quinones^63^, or physiological processes that affect cellular zinc homeostasis. These could relate to reports of the involvement of CZB-containing proteins in cellular responses to other oxidants^21^, pH^25^, and energy taxis^33^. Additionally, the ability for a given CZB domain to perceive and relay these signals will depend on how evolution has tuned its relative zinc-binding affinity over time. For example, though they possess the same core zinc-binding residues, zinc affinity seems to be higher for *H. pylori* TlpD than *E. coli* DgcZ — experiments with the fluorescence probe zinpyr-1 readily competed with DgcZ for zinc (Fig. 3F) but could not extract zinc from TlpD^12^, indicating residues other than those of the zinc-binding core impact affinity. Our conservation analyses note distinct patterns in the α3 region that correspond to functional classes of CZB proteins (Fig. 1C-D), suggesting the stability and dynamics of this region could be particularly important for optimizing zinc affinity.

CZB homologs are present in diverse and evolutionarily distant bacteria, including enteric commensals and pathogens, soil-dwelling and marine species, and extremophiles such as the deep-sea genus *Thermatoga*. Thus, a corollary is that CZB domains have an ancient evolutionary heritage. Though difficulties exist in estimating bacterial protein origins due to horizontal gene transfer, a naïve model based on a consensus of evolutionary divergence timelines of species containing CZB domains indicates these proteins to have been present in the bacterial LUCA approximately four billion years ago (Fig. S2)^49^. Because this would have predated animals, which originated approximately 600 million years ago, the zinc-sensing ability of CZB proteins can be viewed as its ancestral molecular function, and HOCl-sensing is a more recent adaptation coopted by some host-associated species to facilitate the colonization of animals.

## ACKNOWLEDGEMENTS

Funding for this work was provided by the National Institutes of Health, NIGMS under award 1P01GM125576 (K.G.), NIDDK under award numbers R01DK1013145 (K.G.) and F32DK115195 (A.P.), and the NIAID under award number 1K99AI148587 (A.P.). The content is solely the responsibility of the authors and does not necessary represent the official views of the National Institutes of Health. Funding was also provided by the U.K. Biotechnology and Biological Sciences Research Council grant BB/S003339/1 (C.K.C. and P.J.S.). L.Z. and O.M.O. acknowledge Chapman University’s research computing for provision and maintenance of the High-Performance Computing system utilized to generate the QM data presented in this manuscript. Molecular dynamics simulations were conducted using the ARCHER UK National Supercomputing Service.

## AUTHOR CONTRIBUTIONS

All authors contributed to, and approved, the final version of the manuscript. A.P. conceived the project, performed phylogenetic, biochemical, and *in vivo* analyses, and wrote the manuscript. D.A.T. performed protein purifications and contributed to HOCl oxidation and zinc fluorescence assays. R. D. T. designed and performed the DgcZ activity assays and analyses. L.Z. and O.M.O. designed and performed the quantum mechanical analyses and contributed to writing the manuscript. C.K.C. designed and performed the molecular dynamics analyses and contributed to writing the manuscript. T.S. contributed to the conceptualization of the DgcZ activity assays, wrote the corresponding paragraph, and critically reviewed the manuscript. P.J.S. provided critical review of the manuscript and feedback on the molecular dynamics sections. K.G. contributed to project conceptualization, interpretation of data, and writing of the manuscript, and provided critical review and revisions of the manuscript. The authors declare no competing interests.

## MATERIALS & METHODS

### Contact for Reagent and Resource Sharing

Further information and requests for resources and reagents should be directed to and will be fulfilled by the Lead Contact, Dr. Arden Perkins.

#### CZB conservation and phylogenetics

Initial BLAST searches of the non-redundant protein database were performed to identify CZB-containing proteins using the CZB domains from *H. pylori* TlpD, *E. coli* DgcZ, and *S. enterica* McpA as search queries and the software Geneious Prime 2020 with default cutoffs. Automated identification of CZB domains was based on four attributes: (1) the presence of the conserved CZB zinc-binding 3His/1Cys core, (2) the conserved Cx[L/F]GxW[Y/L] motif identified in previous studies^23,26^, (3) sequence coverage across the domain, and (4) reasonable alignment to other confirmed CZB sequences. Sequences that did not meet these qualifications were reviewed manually. BLAST searches were continued iteratively with bona fide CZB sequences until no new sequences emerged. Protein sequences were annotated using Interpro^90^ to identify protein domains and putative transmembrane regions. Sequences were aligned using MUSCLE^91^ and relatedness trees constructed with FastTree^92^. Phylogenetic trees of CZB-containing organisms were constructed with phyloT^93^ and divergence trees were constructed with TimeTree^49^.

#### Molecular dynamics simulation

An intact model of the *E. coli* CZB homodimer (residues 7-126) was constructed based on PDB 3t9o^23^. Coordinates for residues 38-51 from chain B were used to fill in the corresponding residues missing in chain A. Chemical alternations to residue 52 were accomplished using the *psfgen* program in VMD^94^. Force-field parameter and topology files were obtained for cysteine sulfenic acid (Cys-SOH) from an independent study^95^. All CZB models were hydrated with TIP3P and neutralized with 150 mM NaCl, producing systems containing ∼40,500 atoms. Each model was subjected to a conjugant-gradient energy minimization (2,000 steps), followed by a series of equilibration simulations with harmonic positional restraints applied successively to backbone+Zn (5 ns), Cα+Zn (5 ns), and Zn only (5 ns). Three independent, all-atom production simulations (1 μs each) provided the data used for subsequent analysis. All molecular dynamics simulations were performed using NAMD 2.13^96^ and the CHARMM36 force field^97^. Simulations were conducted in the NPT ensemble (1 atm; 310 K) using a 2-fs timestep. Short-range, non-bonded interactions were calculated with a cutoff of 12 Å, and long-range electrostatics were evaluated using the particle-mesh-Ewald method. Molecular visualization and basic structural analyses were carried out in VMD. Representative videos of each simulation are presented in Movies S1-S5.

#### Quantum mechanical analyses

All quantum mechanical analyses on the ligand exchange equilibria reported in this manuscript were performed in *Gaussian 16*^98^. All structures were optimized and vibrational frequency calculations performed in vacuum using B3LYP^99^/SDD^100^(Zn)/6-31+G(d,p)^101–103^ level of theory (Data S1). Electronic energies on the B3LYP-optimized geometries were calculated using the Minnesota functional – M06^104^ – and with the identical mixed basis sets described for optimizations. The optimized complexes were verified as ground states through vibrational frequency analysis^105^. All thermal energies (e.g., ΔG) were calculated at 298 K and 1.0 atm. For each entry reported in Table 1, both sides of the equilibrium were modelled as the cationic Zn^2+^ species complexed with the appropriate exogenous ligand; that is, HOCl or H_2_O on the left side, and CH_3_S(H), CH_3_SO(H), or CH_3_SO_2_(H) on the right side depending on the oxidation state of the zinc-bound sulfur state being displaced.

#### Recombinant protein purification

For recombinant protein expression, Rosetta DE3 cells were transformed with pet28a plasmids from previous work^23^ containing sequences for full-length DgcZ, DgcZ-C52A, and CZB containing 6x-His affinity tags. Equivalent constructs of full length DgcZ-C52D and CZB-C52D mutants were obtained commercially from Genscript. Recombinant proteins were expressed and purified as described previously^12,23^. Frozen stocks were used to inoculate 25 mL of LB+Kan cultures, and grown overnight. The following morning, 5 mL of these overnight cultures were added to each of 4 x 1 L cultures of LB+Kan and grown shaking at 37 °C until they reached OD_600_ 0.6-0.8. Protein expression was induced with 1 mM IPTG, and grown for 3 H, and harvested by centrifugation. Cell pellets were resuspended into ice-cold lysis buffer containing 10 mM imidazole, 50 mM HEPES, 10% glycerol, 300 mM NaCl, and 0.5 mM TCEP (pH 7.9) and lysed by sonication. The cell suspension was centrifuged and the soluble portion retained for affinity chromatography. Lysate was applied to a prepacked gravity column of Ni-NTA agarose beads (Qiagen) equilibrated with lysis buffer. Lysate was incubated with the beads for 10 minutes, and then allowed to flow over the column twice. The column was then washed with lysis buffer until no protein was present in the flow through as determined using a Bradford assay. Purified protein was eluted by adding elution buffer to the column containing 300 mM imidazole, 50 mM HEPES, 300 mM NaCl, and 0.5 mM TCEP (pH 7.9), incubating the buffer in the column for 10 minutes, and then collecting the flow through in fractions. Samples were checked for purity by SDS-PAGE. Fractions containing pure protein were pooled and concentrated by a Pall centrifugal device with a 10 k Da cutoff, and flash-frozen in liquid nitrogen. Prior to biochemical experiments proteins were extensively dialyzed into buffers relevant for the reactions using ThermoScientific mini-dialysis tubes with 3-10 k Da MW cutoffs.

#### Cysteine-sulfenic acid quantification

Reactions were prepared with purified protein (10 µM) dialyzed into PBS (pH 7), 500 µM 5,5-dimethyl-1,3-cyclohexanedione (dimedone), and additions of buffer or HOCl/H_2_O_2_ diluted into PBS buffer. Solutions were strictly pH-controlled by monitoring final solution pH with a pH meter and pH strips. Reactions were allowed to proceed for 10 minutes at room temperature, and then quenched with 100 µM L-methionine. Glutathione disulfide (GSSG) treatments were performed similarly, except CZB protein was pre-treated with 250 µM HOCl, and subsequently with varying GSSG treatments, followed by addition of dimedone. GSSG was chosen to test the reversibility of CZB oxidation because it can form mixed disulfides with cysteines in both sulfenic acid and thiol states, and also is oxidized by HOCl, and therefore serves to fully quench the reaction. 20 µL of samples were dispensed into 180 µL of quenching buffer containing 75 mM H_3_PO_4_, 1 M NaCl and drawn by vacuum through a 0.4 micron polyvinylidene fluoride membrane in a 96-well slot blotter. The membrane was blocked in a buffer of 5 % milk in 50 mM Tris, pH 7.5, 150 mM NaCl, 0.1% Tween-201 (TBST) for 10 minutes, and incubated overnight at room temperature with rabbit α-cysteine-dimedone antibody (Kerafast) at a 1:5000 dilution. Subsequently the membrane was washed three times with 20 ml of TBST for a duration of 15 minutes, and then incubated with goat α-rabbit-HRP secondary antibody (1:5,000) for 1 H. Afterward, the membrane was washed three times for 15 minutes with 20 ml of TBST, and then visualized through chemiluminescence using an ECL kit and Licor imaging system.

#### Fluorescence and circular dichroism assays

The zinc-binding probe zinpyr-1 (Abcam) was used to detect available zinc through fluorescence emission. To avoid cross-reactivity of the probe with HOCl, samples were quenched with methionine or GSSG addition prior to probe addition. Samples were analyzed in black clear bottom 96-well plates with excitation at 488 nm and emission spectra collected from 505-600 nm on a Spectramax i3 plate reader. Intrinsic protein fluorescence was collected similarly with excitation at 280 nm. Circular dichroism spectra were collected using a 1 mm quarts cuvette (Sterna Cells) on a Fluoromax-3 spectrofluorometer.

#### DgcZ activity assays

Reactions monitoring the *in vitro* production of c-di-GMP by DgcZ and cysteine point mutants were carried out in PBS buffer (10 mM Na_2_HPO_4_, 1.8 mM KH_2_PO_4_, 2.7 mM KCl, 137 mM NaCl, pH 7) with 5 mM MgCl2 and 300 µM or 500 μM GTP. Protein was pre-incubated in the reaction buffer for 10 min with HOCl, ZnCl_2_, DTT, or EDTA as necessary prior to addition of GTP. Quantification of GTP and c-di-GMP at specific timepoints was performed as done previously^23^ by chromatography using a Resource Q column and AKTA FPLC. A two-state model was fitted to the EDTA titration data with the amount of c-di-GMP produced being equal to:

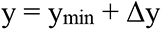

with Δy given by the usual quadratic equation:

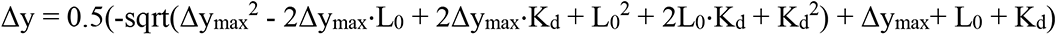

and

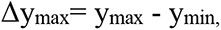

Where L_0_ is ligand concentration. Parameters to be refined were y_min_, y_max_, and K_d_.

#### Bacterial strains

*E. coli* strains, genotypes, relevant phenotypes, and sources are listed in Table S1. All biofilm experiments were performed with MG1655-derivitized strains containing a *csrA* deletion^23,37,38^. CsrA is an RNA-binding protein that directly binds the mRNA of DgcZ (previously called YdeH) and inhibits DgcZ expression^38,106^. The *csrA* deletion background permits a moderate degree of poly-GlcNAc production and biofilm formation under laboratory conditions, similar to that observed *in vivo*, and so represents an *in vitro* mimic of behavior inside a host^38^.

#### Biofilm assays

Unless specified otherwise, cells were prepared for biofilm assays with overnight growth shaking in 5 ml of LB at 37° C, and fresh liquid LB cultures were inoculated in the morning and grown to the desired OD_600_. Static biofilm assays with clear flat bottom microplates were prepared with 200 µl of cells at OD_600_ of 0.5 (Fig. 5A-D, G) or 1.0 (Fig. S4A), covered with parafilm and grown for 16 or 24 H at 25 or 30° C as indicated. Static assays with 15 mm petri dishes were setup similarly but using 3 ml of cell culture at OD_600_ 0.5 and holes drilled into the lids to hold a 20 µl pipette with treatment solution. Treatments were applied with either direct addition (Fig. 5A-B, Fig. 6E-I, Fig. S4A,B,D-E), exposure to 20 µl treatment point-sources with a Rainin 96-well liquidator (Fig. 5C-D, Fig. S4C), or exposure to a 20 µl Rainin pipette tip sealed with parafilm containing 20 µl of treatment solution (Fig. 5E-F). Liquid culture rocking biofilm assays were conducted using 1 ml of OD_600_ 0.5 cells in LB media in glass 10 x 75 mm culture tubes (Fisherbrand) and incubated upright at 30° C for 24 H with near-horizontal rocking (Fig. 6C-I, Fig. S4C-D).

Biofilm was quantified by crystal violet staining as done previously^23^. Non-adhered cells were removed and samples were washed twice with deionized water and stained with 0.1 % crystal violet for 30 minutes. Excess stain was removed and samples were washed twice with deionized water, dried, and then treated with a destain solution containing of 30 % methanol, 10 % acetic acid (equal in volume to cell culture) for 30 minutes. Samples were then quantified by measuring absorbance at 562 nm. Quantification of samples are presented as either raw Abs_562_ values or normalized relative to the average of the wild type untreated control strain in each experiment (Abs_562_ sample/average Abs_562_ of untreated wild type replicates). For quantification of biofilm distribution and pellicle formation high resolution microscopy images were captured of crystal violet-stained culture tubes on a Nikon Z20 dissecting scope and intensity profiles of tubes were collected along the tube length (Fig. 6E-I). Note: images of pellicle formation presented in Fig. 6E-I are illustrative and were not the images used for quantification. Quantification of pellicle formation was performed by integrating the intensity in the range of pixel 2000-2500 (Fig. S4C).

#### Congo red assays

For assaying Congo red dye retention with growth on agar plates, 2 µl of cells either from overnight cultures or mid-log exponential growth cultures at OD_600_ 0.5 were spotted onto LB agar plates containing 25 µg/ml Congo red dye and grown for 24 H at room temperature (Fig. 6A-B). Plates were imaged under identical lighting with a Leica MZ10F scope equipped with an MC190HD camera. Quantification of dye uptake was performed in ImageJ^107^ applying a red hue threshold of 1-14 in HSB color space. Images were then clustered into red, brown, and black (background) color bins using the Color Segmentation plugin and clustering pixels according to the K-Means algorithm. Pixels from color-thresholded and clustered images were then counted using the Color Counter plugin. For visualizing Congo red straining of liquid cultures, cells were treated with 25 μg/ml dye and incubated for 10 minutes (Fig. 6C).

## Notes

### Competing Interest Statement

The authors have declared no competing interest.

