## Supplemental Information for "A bacterial inflammation sensor regulates c-di-GMP signaling, adhesion, and biofilm formation"

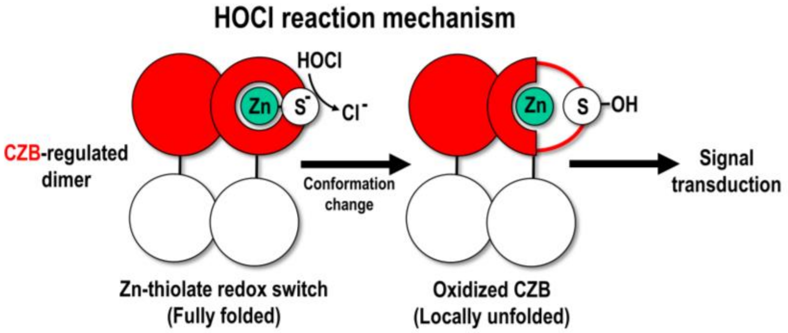


Fig. S1. Mechanism of CZB HOCl sensing proposed in Perkins *et al*. 2019[12]. CZB domains (large red circles) form dimers and regulate the function of other protein domains (large white circles). Each CZB monomer contains a conserved cysteine thiolate ($S^-$) bound to zinc (green). Oxidation of the zinc-bound thiolate by HOCl to cysteine-sulfenic acid (SOH) stimulates a local unfolding of the region containing the cysteine. This conformational change induces downstream signal transduction possibly through alterations in structure or dynamics of regulated proteins or protein-protein interactions. Note, we refine this model later in the study (see Fig. 7).

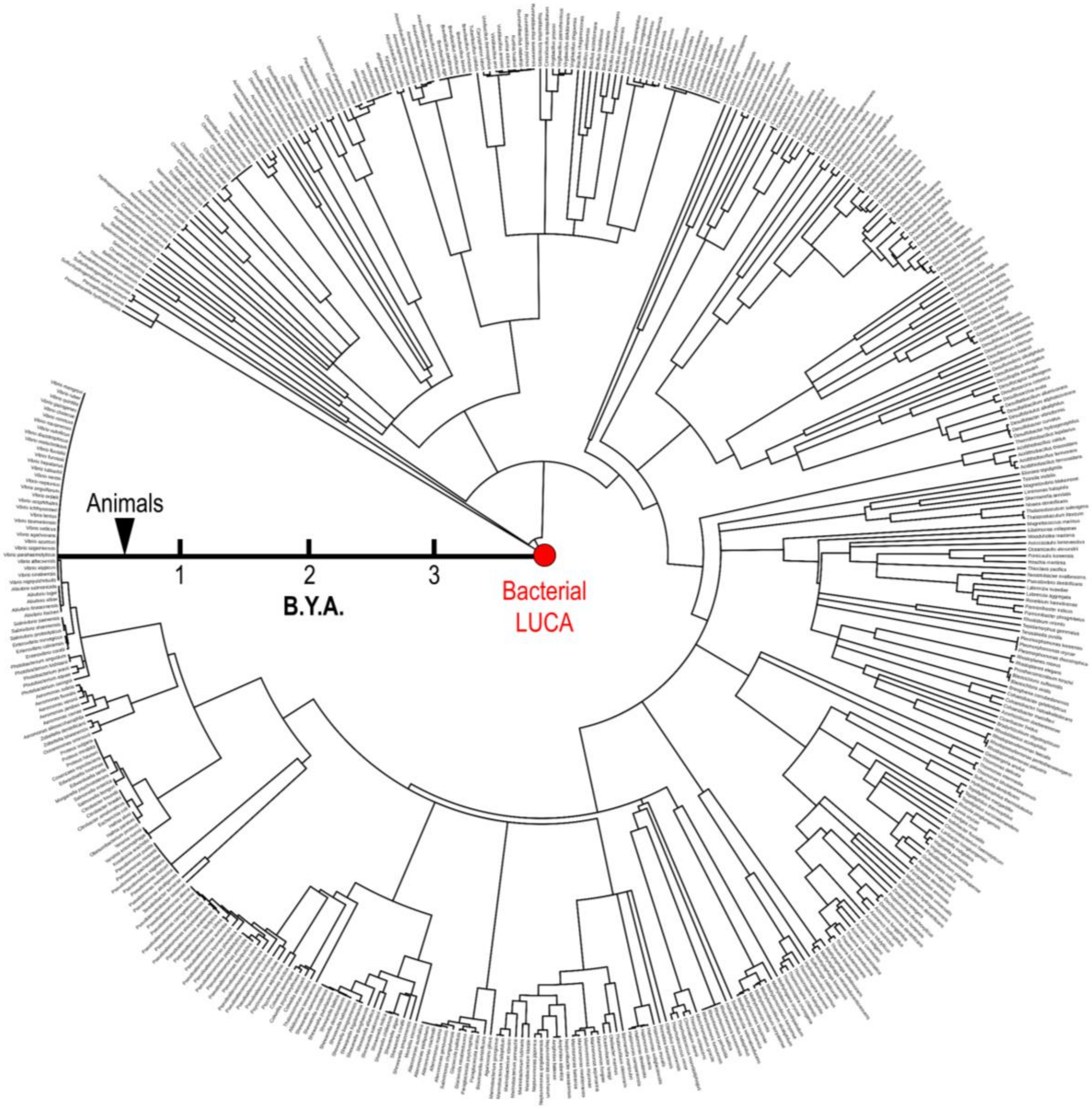


Fig. S2. Divergence tree for CZB-containing species. Scale reflects billions of years ago (B.Y.A.) from present day (circle exterior) dating back to the bacterial last universal common ancestor (LUCA, red circle). The approximate origin of animals is noted (black arrow).

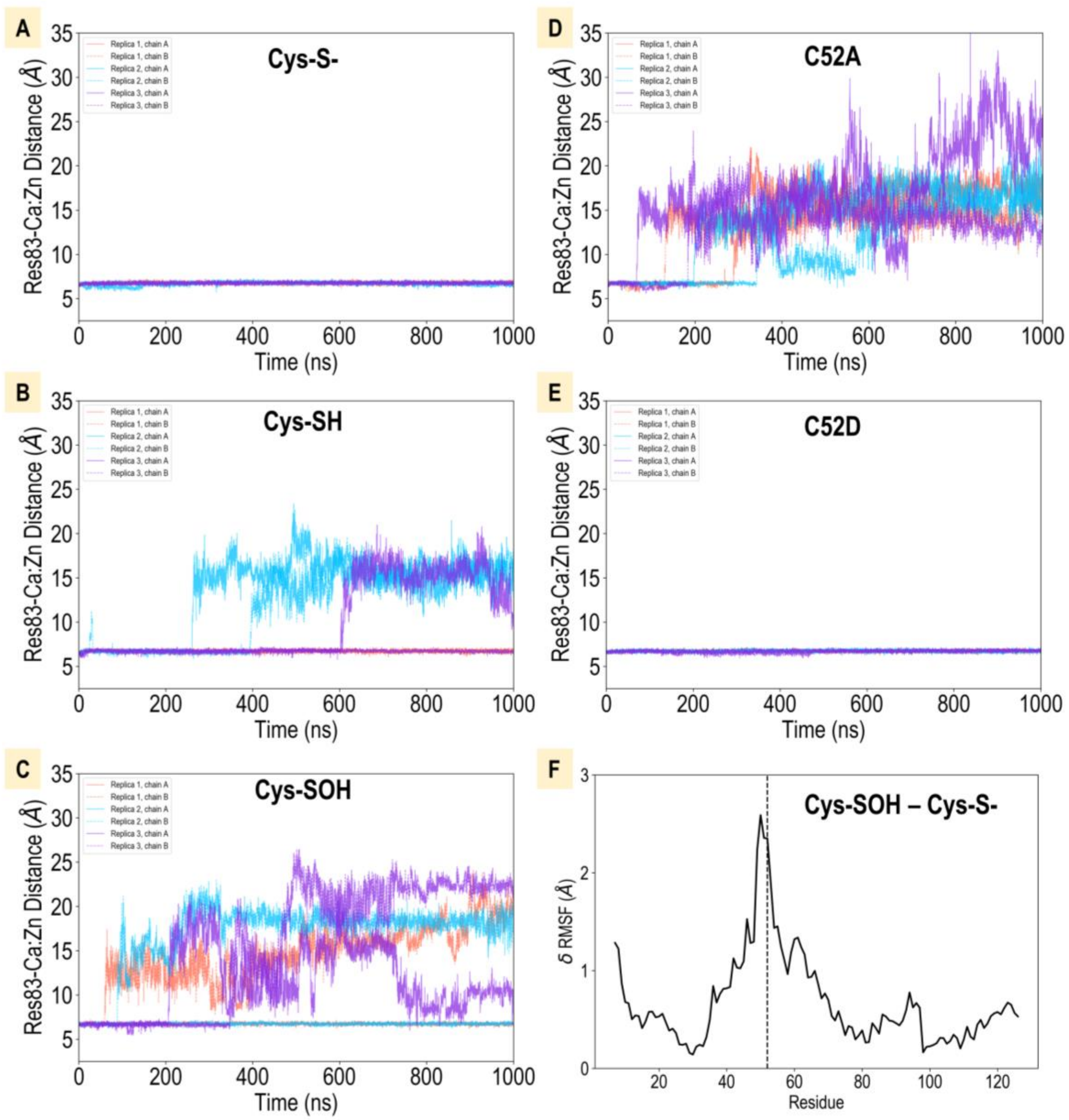


Fig. S3. Structural impacts of C52 redox state and mutation. A-E. Zinc release events monitored by distance between the zinc and zinc-coordinating H83. F. Difference in average RMSF between Cys-SOH and Cys-S$^-$ simulations.

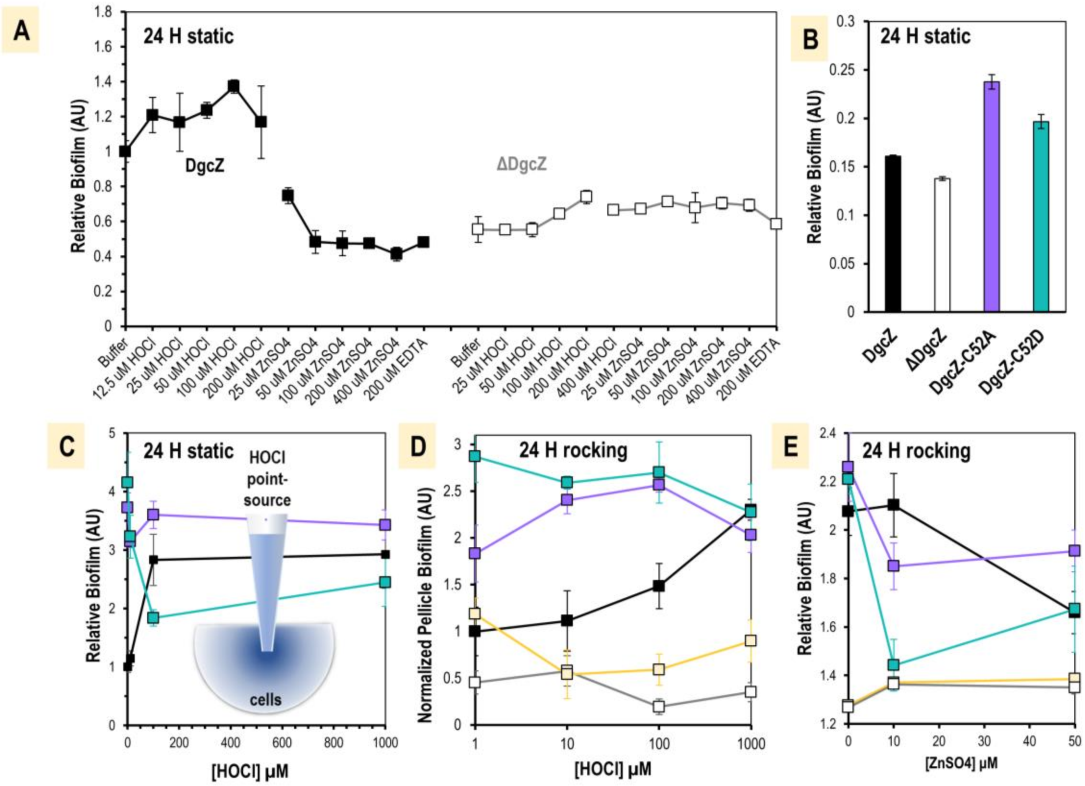


Fig. S4. Biofilm quantification for wildtype and mutant *E. coli* strains. A. Biofilm formation of $OD_{600}$ 1.0 wild type and *ΔdgcZ* cells. Cells were dispensed in a 96-well microplate, treated as indicated, and grown statically for 24 H in LB at 30° C, n=4. B. Biofilm formation of wild type and mutant lines after 24 H static growth in LB at 30° C, n=4. C. Biofilm formation for cells as in B, but exposed to a point source of HOCl, n=6. D. Pellicle formation quantified from Fig. 6E-I for the region spanning pixels 2000-25000 for wild type (black), *ΔdgcZ* (gray), *dgcZ*$^{C52A}$ (violet), *dgcZ*$^{C52D}$ (teal), *dgcZ*$^{E208Q}$ (light yellow), n=3. E. Biofilm formation after treatments with $ZnSO_4$ and incubated with gentle rocking for 24 H at 30° C, n=3. Data points are sample means and error bars are standard error of the mean.

Movie S1: Representative molecular dynamics simulation of wild type CZB in the C52-$S^-$ state. Viewable at: https://youtu.be/eaH6nVFG90s

Movie S2: Representative molecular dynamics simulation of wild type CZB in the C52-SH state. Viewable at: https://youtu.be/m3WnHseUmsI

Movie S3: Representative molecular dynamics simulation of wild type CZB in the C52-SOH state. Viewable at: https://youtu.be/6RXAk-35m7Y

Move S4: Representative molecular dynamics simulation of C52A CZB. Viewable at: https://youtu.be/QkzLckjUUOw

Movie S5: Representative molecular dynamics simulation of C52D CZB. Viewable at: https://youtu.be/v7ke5Ag1UUM

Table S1. *E. coli* bacterial strains

| **Strain** | **Genotype** | **Relevant Phenotype** | **References/source** |
|---|---|---|---|
| AB958 | csrA::Tn5Δ(kan)::Frt | csrA-deletion, expresses wild type *dgcZ* | [23,38] |
| AB959 | csrA::Tn5Δ(kan):: (kan)::Frt ΔydeH::Frt | *ΔdgcZ*, deletion | [23,37] |
| AB1299 | csrA::Tn5Δ(kan):: (kan)::Frt ydeH Frt ydeH-1(E208Q) | $dgcZ^{E208Q}$, catalytically inactive | [37] |
| AP1 | csrA::Tn5Δ(kan)::Frt ydeH (C52A) | $dgcZ^{C52A}$ mutant | This work |
| AP2 | csrA::Tn5Δ(kan)::Frt ydeH (C52D) | $dgcZ^{C52D}$ mutant | This work |
| Rosetta (DE3) | F– ompT $hsdS_B(r_B– m_B–)$ gal dcm (DE3) $pRARE^2$ ($Cam^R$) | Protein expression | Novagen |

Data S1. Data and information relating to quantum mechanics calculations.

Full authorship of Gaussian 16:
Gaussian 16, Revision B.01, M. J. Frisch, G. W. Trucks, H. B. Schlegel, G. E. Scuseria, M. A. Robb, J. R. Cheeseman, G. Scalmani, V. Barone, G. A. Petersson, H. Nakatsuji, X. Li, M. Caricato, A. V. Marenich, J. Bloino, B. G. Janesko, R. Gomperts, B. Mennucci, H. P. Hratchian,

J. V. Ortiz, A. F. Izmaylov, J. L. Sonnenberg, D. Williams-Young, F. Ding, F. Lipparini, F. Egidi, J. Goings, B. Peng, A. Petrone, T. Henderson, D. Ranasinghe, V. G. Zakrzewski, J. Gao, N. Rega, G. Zheng, W. Liang, M. Hada, M. Ehara, K. Toyota, R. Fukuda, J. Hasegawa, M. Ishida, T. Nakajima, Y. Honda, O. Kitao, H. Nakai, T. Vreven, K. Throssell, J. A. Montgomery, Jr., J. E. Peralta, F. Ogliaro, M. J. Bearpark, J. J. Heyd, E. N. Brothers, K. N. Kudin, V. N. Staroverov, T. A. Keith, R. Kobayashi, J. Normand, K. Raghavachari, A. P. Rendell, J. C. Burant, S. S. Iyengar, J. Tomasi, M. Cossi, J. M. Millam, M. Klene, C. Adamo, R. Cammi, J. W. Ochterski, R. L. Martin, K. Morokuma, O. Farkas, J. B. Foresman, and D. J. Fox, Gaussian, Inc., Wallingford CT, 2016.

### Calculation Parameters, Geometries, Energies, and Vibrational Frequencies

**$[Zn(Im)_3(CH_3S)]^+$ Complexed with $H_2O$**

---------------------------------------

Gaussian 16: ES64L-G16RevB.01 20-Dec-2017

---------------------------------------

### B3LYP/GEN PSEUDO=READ gfprint gfinput scf=(direct,tight,maxcycle=300 ,xqc) opt=(maxcycle=250) freq=noraman EDU\13-Jun-2020\0\\# B3LYP/GEN PSEUDO=READ gfprint gfinput scf=(direct ,tight,maxcycle=300,xqc) opt=(maxcycle=250) freq=noraman\\zn_3imid_1th

---------------------------------------

Full point group C1 NOp 1
Stoichiometry C13H23N6OSZn(1+) Framework group C1[X(C13H23N6OSZn)]

---------------------------------------

Num atoms: 45
Charge = 1 Multiplicity = 1

---------------------------------------

SCF = -1538.30598580 | Predicted change in Energy=-9.336652D-09

Optimization completed.
Maximum Force 0.000041 0.000450 YES
RMS Force 0.000003 0.000300 YES
Maximum Displacement 0.001110 0.001800 YES
RMS Displacement 0.000310 0.001200 YES

---------------------------------------

| Atom Type | Coordinates (in Angstroms) X | Y | Z |
|---|---|---|---|
| Zn | 0.241178 | -0.220453 | 0.569254 |
| N | 1.822423 | 0.486178 | -0.561720 |
| C | 1.967967 | 1.720910 | -1.175692 |
| C | 3.204197 | 1.808187 | -1.763628 |
| C | 2.954141 | -0.176174 | -0.767090 |
| N | -1.050791 | 1.409004 | 0.699565 |
| C | -1.786343 | 2.123652 | -0.234317 |
| C | -2.466245 | 3.143149 | 0.385007 |
| C | -1.274417 | 1.982646 | 1.873529 |
| N | -0.720721 | -1.642755 | -0.588213 |
| C | -1.976780 | -1.622620 | -1.174634 |
| C | -2.219373 | -2.818071 | -1.802781 |
| C | -0.198604 | -2.835063 | -0.848815 |
| S | 0.903764 | -0.947606 | 2.626132 |
| C | -0.493677 | -2.032299 | 3.166907 |
| H | -0.211128 | -2.483945 | 4.120141 |
| H | -1.408470 | -1.453647 | 3.316319 |
| H | -0.689759 | -2.828807 | 2.445364 |
| H | 3.173022 | -1.174432 | -0.405854 |
| H | 1.186118 | 2.464527 | -1.147453 |
| H | 0.771310 | -3.184810 | -0.513458 |
| H | -2.626366 | -0.763904 | -1.098595 |
| H | -0.840084 | 1.668214 | 2.812483 |
| H | -1.779800 | 1.866325 | -1.283284 |
| C | -3.390827 | 4.195184 | -0.133197 |
| H | -3.008516 | 5.202854 | 0.065102 |
| H | -3.506010 | 4.088711 | -1.214045 |
| H | -4.386048 | 4.116544 | 0.318785 |
| C | 3.874785 | 2.894466 | -2.538512 |
| H | 4.132291 | 2.569268 | -3.552942 |
| H | 3.208081 | 3.755378 | -2.623724 |
| H | 4.794796 | 3.232067 | -2.047907 |
| C | -3.392081 | -3.329266 | -2.573252 |
| H | -3.120891 | -3.580023 | -3.605075 |
| H | -3.821284 | -4.224567 | -2.109297 |
| H | -4.173137 | -2.566706 | -2.609792 |
| H | 4.738285 | 0.314444 | -1.769242 |
| N | 3.806442 | 0.589369 | -1.489431 |
| H | -0.913320 | -4.511641 | -1.901617 |
| N | -1.073270 | -3.566353 | -1.580368 |
| H | -2.446184 | 3.627407 | 2.470888 |
| N | -2.121494 | 3.028517 | 1.723351 |
| H | 2.191864 | -2.454240 | 1.505710 |
| H | 3.183767 | -3.614790 | 1.161389 |
| O | 2.593189 | -2.963829 | 0.761010 |

---------------------------------------

Statistical Thermodynamic Analysis
Temperature 298.150 Kelvin. Pressure 1.00000 Atm.

---------------------------------------

SCF = -1538.30598580 | Predicted change in Energy=-9.336652D-09
Zero-point correction (ZPE) = -1537.9387898 0.367196
Internal Energy (U) = -1537.9106478 0.395338
Enthalpy (H) = -1537.9097038 0.396282
Gibbs Free Energy (G) = -1538.0015888 0.304397

---------------------------------------

Frequencies
20.2550 26.8233 32.1245
36.0735 40.6855 53.6729
54.9480 58.9418 72.6295
95.7881 99.9315 101.6036
105.7067 116.4920 121.6175
127.0188 128.0459 132.5301
146.5645 148.8251 161.2286
174.3677 180.4455 190.1419
223.3645 229.5147 266.6940
271.0290 273.7737 310.4695
332.3803 343.6308 345.1490
350.0278 447.1385 600.2125
604.0934 605.3391 655.8571
657.6160 657.9206 666.1920
666.5742 667.8031 675.8653
678.1473 680.9764 691.1289
719.9447 819.6590 826.0538
831.6427 859.1108 900.0419
905.8417 966.6479 968.5200
970.1850 971.4355 980.6876
989.8192 991.3114 991.3554
1031.1701 1034.3592 1035.4509
1066.6179 1066.9275 1067.0648

1138.2141 1141.8065 1145.9941
1152.4079 1160.5598 1164.1231
1260.4163 1271.0166 1272.7094
1288.6769 1299.3502 1301.0068
1366.9237 1376.3139 1378.4743
1379.9609 1430.4914 1430.6073
1430.8420 1447.2611 1447.4552
1448.6593 1485.2150 1485.3515
1485.4059 1485.4891 1487.5487
1496.4378 1497.1656 1497.5255
1535.9760 1541.2421 1546.6321
1624.0752 1640.0275 1642.8347
1644.1292 3045.1550 3045.4956
3046.4862 3055.0197 3101.0679
3101.5738 3103.1044 3134.7610
3143.4128 3143.7790 3144.6624
3146.3000 3230.6690 3235.4442
3275.5203 3278.8380 3285.5113
3285.8282 3411.7023 3647.5472
3650.2643 3651.1281 3867.5657

**$[Zn(Im)_3(OH)]^+$ Complexed with $CH_3SH$**

---------------------------------------
Gaussian 16: ES64L-G16RevB.01 20-Dec-2017
---------------------------------------
### B3LYP/GEN PSEUDO=READ gfprint gfinput scf=(direct,tight,maxcycle=300 ,xqc) opt=(maxcycle=250) freq=noraman EDU\12-Jun-2020\0\\# B3LYP/GEN PSEUDO=READ gfprint gfinput scf=(direct ,tight,maxcycle=300,xqc) opt=(maxcycle=250) freq=noraman\\zn_3im_oh_ch
---------------------------------------
Full point group C1 NOp 1
Stoichiometry C13H23N6OSZn(1+) Framework group C1[X(C13H23N6OSZn)]
---------------------------------------
Num atoms: 45
Charge = 1 Multiplicity = 1
---------------------------------------
SCF = -1538.28208814 | Predicted change in Energy=-4.723285D-09

Optimization completed.
Maximum Force 0.000010 0.000450 YES
RMS Force 0.000002 0.000300 YES
Maximum Displacement 0.001391 0.001800 YES
RMS Displacement 0.000340 0.001200 YES

---------------------------------------

| Atom Type | X | Y | Z |
|---|---|---|---|
| Zn | 0.431560 | 0.093621 | -0.379935 |
| N | 0.149879 | 1.985788 | 0.406050 |
| C | 0.409938 | 2.608045 | 1.617408 |
| C | -0.162687 | 3.855210 | 1.633514 |
| C | -0.574231 | 2.836405 | -0.310605 |
| N | 2.452857 | -0.294517 | -0.447768 |
| C | 3.070862 | -1.532024 | -0.542979 |
| C | 4.423442 | -1.368623 | -0.708507 |
| C | 3.411472 | 0.614299 | -0.555581 |
| N | -0.376732 | -1.295316 | 0.902820 |
| C | 0.153124 | -1.948376 | 2.006060 |
| C | -0.801006 | -2.754017 | 2.574538 |
| C | -1.638574 | -1.693282 | 0.795417 |
| H | -0.953468 | 2.634912 | -1.302717 |
| H | 0.978385 | 2.123921 | 2.397545 |
| H | -2.351281 | -1.360941 | 0.049618 |
| H | 1.176018 | -1.800044 | 2.318524 |
| H | 3.275526 | 1.685263 | -0.531178 |
| H | 2.507558 | -2.451813 | -0.494069 |
| C | 5.541021 | -2.345913 | -0.868091 |
| H | 6.054594 | -2.217537 | -1.827516 |
| H | 6.284428 | -2.245219 | -0.069327 |
| H | 5.149546 | -3.364802 | -0.833792 |
| C | -0.199341 | 4.931441 | 2.668153 |
| H | 0.259631 | 5.858355 | 2.305798 |
| H | -1.225976 | 5.160788 | 2.975505 |
| H | 0.350482 | 4.611720 | 3.556103 |
| C | -0.773238 | -3.663919 | 3.758332 |
| H | -0.988465 | -4.700946 | 3.476744 |
| H | 0.216124 | -3.642241 | 4.220542 |
| H | -1.503597 | -3.361730 | 4.517454 |
| H | -1.317188 | 4.765878 | 0.074902 |
| N | -0.779308 | 3.972007 | 0.395826 |
| H | -2.825984 | -3.011605 | 1.922329 |
| N | -1.926554 | -2.569987 | 1.785299 |
| H | 5.497475 | 0.479850 | -0.819778 |
| N | 4.610909 | 0.005367 | -0.711643 |
| H | -2.502396 | 0.395677 | -1.818480 |
| S | -3.794070 | 0.492343 | -1.377418 |
| C | -4.639927 | 0.036434 | -2.943663 |
| H | -4.362441 | -0.968941 | -3.266202 |
| H | -5.712661 | 0.060187 | -2.743360 |
| H | -4.412562 | 0.756880 | -3.731490 |
| O | -0.574800 | 0.282836 | -1.969381 |
| H | -0.253346 | -0.185435 | -2.748001 |

Atom Coordinates (in Angstroms)

---------------------------------------
Statistical Thermodynamic Analysis
Temperature 298.150 Kelvin. Pressure 1.00000 Atm.
---------------------------------------
SCF = -1538.28208814 | Predicted change in Energy=-4.723285D-09
Zero-point correction (ZPE) = -1537.91823814 0.36385
Internal Energy (U) = -1537.88981114 0.392277
Enthalpy (H) = -1537.88886714 0.393221
Gibbs Free Energy (G) = -1537.98486514 0.297223
---------------------------------------
Frequencies
13.9112 18.3992 19.7120
23.0818 28.7039 32.6184
40.8428 44.2765 57.6338
68.3897 72.0408 85.9261
94.5528 104.8840 116.3780
118.6751 127.4639 127.8088
138.7676 151.5789 156.2759
171.4773 188.7991 200.3129
227.3353 231.5302 266.8849
270.2399 271.7821 276.8503

343.4766 345.6078 347.9717
532.4502 548.1951 601.9438
605.4066 607.9735 655.6359
656.2949 658.6395 666.2572
667.7839 668.1906 675.0375
677.3551 679.4492 702.1332
745.2974 814.3592 821.7281
821.7865 836.3486 872.9525
879.8745 886.6026 966.4743
970.7543 972.5641 977.9651
989.6718 990.2176 990.3889
1031.7676 1032.1054 1033.6485
1067.0239 1067.1652 1067.6315
1137.6350 1140.7869 1142.0537
1147.5948 1151.9601 1155.6675
1163.0122 1260.4053 1260.7258
1270.0188 1289.9601 1291.6680
1295.2831 1371.8852 1374.5657
1376.1283 1378.3781 1430.6330
1430.6405 1430.8873 1446.9893
1448.8540 1449.6766 1481.2283
1485.2806 1485.4394 1485.5123
1491.1086 1496.3791 1496.8723
1497.2443 1532.2525 1540.6852
1545.0498 1641.3912 1642.7602
1643.8112 2433.7805 3045.2924
3045.6533 3047.3120 3067.0862
3101.2866 3101.7853 3104.3292
3143.2925 3143.5799 3146.5698
3154.6609 3161.9859 3232.0746
3279.6268 3281.0656 3284.6971
3284.8139 3291.7069 3647.9782
3648.6682 3649.9461 3880.9991

**$[Zn(Im)_3(CH_3S)]^+$ Complexed with HOCl**

---------------------------------------
Gaussian 16: ES64L-G16RevB.01 20-Dec-2017
---------------------------------------
### B3LYP/GEN PSEUDO=READ gfprint gfinput
scf=(direct,tight,maxcycle=300
,xqc) opt=(maxcycle=250) freq=noraman
AN.EDU\28-Jun-2019\0\\# B3LYP/GEN PSEUDO=READ
gfprint gfinput scf=(dir
ect,tight,maxcycle=300,xqc) opt=(maxcycle=250)
freq=noraman\\zn_2im_in
---------------------------------------
Full point group C1 NOp 1
Stoichiometry C13H22ClN6OSZn(1+) Framework group
C1[X(C13H22ClN6OSZn)]
---------------------------------------
Num atoms: 45
Charge = 1 Multiplicity = 1
---------------------------------------
SCF = -1997.82685506 | Predicted change in Energy=-1.102649D-08

Optimization completed.

| | | | |
|---|---|---|---|
| Maximum Force | 0.000026 | 0.000450 | YES |
| RMS Force | 0.000004 | 0.000300 | YES |
| Maximum Displacement | 0.001040 | 0.001800 | YES |
| RMS Displacement | 0.000244 | 0.001200 | YES |

---------------------------------------

| Atom Type | Coordinates (in Angstroms) X | Y | Z |
|---|---|---|---|
| Zn | -0.067419 | 0.118299 | -0.062445 |
| O | -3.466252 | -0.265042 | -0.894276 |
| N | 0.006536 | -1.558694 | -1.267616 |
| C | 1.108274 | -2.328692 | -1.609710 |
| C | 0.712489 | -3.400486 | -2.368910 |
| C | -1.051719 | -2.151248 | -1.807291 |
| N | -0.314721 | 1.763362 | -1.290863 |
| C | 0.562803 | 2.794738 | -1.589484 |
| C | -0.050520 | 3.708859 | -2.408604 |
| C | -1.452815 | 2.041817 | -1.915088 |
| S | -1.658802 | -0.086553 | 1.570816 |
| H | -2.356293 | 1.445965 | -1.872608 |
| H | 1.565348 | 2.822228 | -1.190624 |
| H | -2.075313 | -1.808890 | -1.721585 |
| H | 2.104259 | -2.071495 | -1.282931 |
| C | 1.470332 | -4.527867 | -2.989106 |
| H | 1.356572 | -4.541083 | -4.078994 |
| H | 2.534722 | -4.426801 | -2.766146 |
| H | 1.138621 | -5.498190 | -2.602598 |
| C | 0.421632 | 4.986973 | -3.019769 |
| H | 0.389361 | 4.948193 | -4.114473 |
| H | -0.185319 | 5.839004 | -2.693377 |
| H | 1.454652 | 5.181047 | -2.723110 |
| H | -2.057909 | 3.633830 | -3.150463 |
| N | -1.327585 | 3.203537 | -2.598835 |
| H | -1.284509 | -3.888981 | -2.969238 |
| N | -0.662666 | -3.260092 | -2.478839 |
| C | -1.962527 | 1.656528 | 2.108456 |
| H | -1.080059 | 2.070914 | 2.601226 |
| H | -2.236497 | 2.296331 | 1.266873 |
| H | -2.786990 | 1.642284 | 2.823856 |
| H | 1.272217 | -0.159806 | 2.760119 |
| C | 4.060186 | 0.437934 | 1.154864 |
| C | 3.093922 | 0.485918 | 0.180647 |
| N | 1.846155 | 0.249404 | 0.739049 |
| C | 2.044882 | 0.057266 | 2.035824 |
| N | 3.363806 | 0.163973 | 2.322227 |
| H | -3.129144 | -0.262382 | 0.053767 |
| H | 3.770664 | 0.058402 | 3.242249 |
| H | 3.220393 | 0.671172 | -0.875873 |
| C | 5.541728 | 0.619873 | 1.110963 |
| H | 5.863517 | 1.462607 | 1.733121 |
| H | 6.069232 | -0.276199 | 1.456766 |
| H | 5.861193 | 0.820637 | 0.086073 |
| Cl | -5.169435 | -0.541595 | -0.821042 |

---------------------------------------
Statistical Thermodynamic Analysis
Temperature 298.150 Kelvin. Pressure 1.00000 Atm.
---------------------------------------
SCF = -1997.82685506 | Predicted change in Energy=-1.102649D-08
Zero-point correction (ZPE) = -1997.46959606 0.357259
Internal Energy (U) = -1997.44031606 0.386539
Enthalpy (H) = -1997.43937206 0.387483
Gibbs Free Energy (G) = -1997.53621206 0.290643
---------------------------------------
Frequencies

19.0952 22.0464 27.8407
31.2347 34.2714 40.3056
49.5576 52.4118 53.9321
56.4803 71.8354 93.9032
97.6179 103.4678 106.8610
113.6326 116.4848 126.0880
127.9799 135.9531 145.2627
149.0088 160.7100 175.4330
190.4575 208.6604 225.6950
233.8688 267.2750 271.4129
274.1062 327.6301 343.7450
344.9864 350.5934 604.3645
607.1668 607.4511 656.8724
658.4573 658.7894 666.5617
666.6698 667.7507 676.9988
678.0502 680.1036 690.7620
713.2067 761.6290 818.9885
824.9435 830.2086 855.1362
885.4707 892.9951 967.4896
969.3802 970.7710 973.9674
983.2853 989.9575 991.2419
991.2821 1031.2115 1034.2135
1035.3800 1066.6296 1067.0309
1067.1262 1137.8662 1141.5744
1145.9304 1153.6234 1159.8219
1162.5878 1260.2535 1268.9983
1270.7346 1288.8843 1296.4165
1297.4250 1315.1025 1369.1638
1375.7586 1377.5247 1379.0874
1430.6536 1430.8053 1431.0253
1447.4475 1447.8930 1449.0629
1484.6233 1485.0942 1485.2690
1485.3211 1487.2466 1496.4029
1497.1551 1497.4946 1536.7701
1539.8030 1545.2532 1640.3478
1642.9196 1644.1388 3046.0171
3046.4017 3047.1139 3059.1813
3102.4026 3102.9651 3104.0675
3111.9762 3141.4609 3144.0184
3144.4847 3145.2355 3152.6400
3246.9036 3253.3020 3276.8700
3279.9640 3286.4763 3286.7526
3646.4813 3647.9746 3648.9042

**$[Zn(Im)_3(OCl)]^+$ Complexed with $CH_3SH$**

---------------------------------------
Gaussian 16: ES64L-G16RevB.01 20-Dec-2017
---------------------------------------
### B3LYP/GEN PSEUDO=READ gfprint gfinput scf=(direct,tight,maxcycle=300
,xqc) opt=(maxcycle=250) freq=noraman
AN.EDU\10-Jun-2020\0\\# B3LYP/GEN PSEUDO=READ gfprint gfinput scf=(dir
ect,tight,maxcycle=300,xqc) opt=(maxcycle=250) freq=noraman\\zn_3im_oc
---------------------------------------
Full point group C1 NOp 1
Stoichiometry C13H22ClN6OSZn(1+) Framework group C1[X(C13H22ClN6OSZn)]
---------------------------------------
Num atoms: 45
Charge = 1 Multiplicity = 1
---------------------------------------
SCF = -1997.81624527 | Predicted change in Energy=-5.143094D-09

Optimization completed.

| | | | | |
|---|---|---|---|---|
| Maximum Force | | 0.000032 | 0.000450 | YES |
| RMS | Force | 0.000003 | 0.000300 | YES |
| Maximum Displacement | | 0.001473 | 0.001800 | YES |
| RMS | Displacement | 0.000286 | 0.001200 | YES |

---------------------------------------

| Atom Type | Coordinates (in Angstroms) X | Y | Z |
|---|---|---|---|
| Zn | -0.075288 | 0.964805 | 0.434182 |
| N | -1.460627 | 1.908975 | -0.741209 |
| C | -2.115772 | 1.599146 | -1.922446 |
| C | -2.999972 | 2.598956 | -2.240356 |
| C | -1.936259 | 3.079482 | -0.336951 |
| N | -0.480381 | -1.003924 | 0.663590 |
| C | -0.203058 | -2.073689 | -0.174653 |
| C | -0.623654 | -3.241672 | 0.410722 |
| C | -1.060721 | -1.516016 | 1.745309 |
| N | 1.854980 | 1.321018 | -0.153335 |
| C | 2.590804 | 1.108523 | -1.308348 |
| C | 3.863590 | 1.594284 | -1.145611 |
| C | 2.661846 | 1.930477 | 0.706610 |
| H | -1.636852 | 3.591803 | 0.566249 |
| H | -1.919297 | 0.684879 | -2.461943 |
| H | 2.377862 | 2.245621 | 1.701542 |
| H | 2.165864 | 0.630686 | -2.178222 |
| H | -1.398053 | -0.947225 | 2.599654 |
| H | 0.282766 | -1.941259 | -1.129650 |
| C | -0.570253 | -4.662671 | -0.046481 |
| H | 0.021728 | -5.283245 | 0.635154 |
| H | -1.571319 | -5.102210 | -0.114840 |
| H | -0.110190 | -4.718226 | -1.035583 |
| C | -3.946296 | 2.778110 | -3.381441 |
| H | -4.984029 | 2.854465 | -3.037593 |
| H | -3.714783 | 3.678577 | -3.961572 |
| H | -3.880901 | 1.921265 | -4.055510 |
| C | 5.049528 | 1.628311 | -2.052310 |
| H | 5.897120 | 1.076229 | -1.630656 |
| H | 4.796990 | 1.170073 | -3.011005 |
| H | 5.378535 | 2.654972 | -2.248973 |
| H | -3.378330 | 4.390772 | -1.129251 |
| N | -2.863139 | 3.525014 | -1.216030 |
| H | 4.668282 | 2.552984 | 0.592401 |
| N | 3.878536 | 2.109930 | 0.142451 |
| H | -1.574511 | -3.479551 | 2.329486 |
| N | -1.162812 | -2.856315 | 1.626851 |
| O | -0.004504 | 2.068990 | 2.027745 |
| Cl | -1.284651 | 1.807396 | 3.151842 |
| H | -3.748243 | -4.507700 | 4.212418 |
| C | -1.763060 | -5.002655 | 5.512821 |
| H | -1.674599 | -3.979021 | 5.879052 |
| H | -2.318193 | -5.615773 | 6.223710 |
| H | -0.766240 | -5.427653 | 5.383052 |
| S | -2.570444 | -5.060937 | 3.862128 |

---------------------------------------
Statistical Thermodynamic Analysis
Temperature 298.150 Kelvin. Pressure 1.00000 Atm.

---------------------------------------
SCF = -1997.81624527 |  Predicted change in Energy=-5.143094D-09
Zero-point correction (ZPE) = -1997.46199527    0.35425
Internal Energy (U) = -1997.43202027   0.384225
Enthalpy (H) = -1997.43107627   0.385169
Gibbs Free Energy (G) = -1997.53417827   0.282067
---------------------------------------
Frequencies
7.0194 10.8021 17.0672
19.3303 20.6701 28.0985
31.7096 38.8284 40.2897
42.6727 67.0908 70.2615
79.6267 88.5744 96.1516
111.1168 112.1304 117.3384
122.9664 128.4090 140.6881
148.1885 151.5812 160.4009
180.2940 202.9980 240.2504
249.0005 265.8850 271.5899
273.3096 311.4724 344.1038
346.2352 353.1953 443.2741
602.2531 604.3093 638.4660
654.0629 656.1811 667.0154
668.4769 668.8017 669.6303
674.7824 676.5467 692.0183
735.0071 796.3091 804.2252
816.9283 819.2136 838.9267
855.2453 864.0632 867.0301
969.3282 969.9380 978.0253
986.5913 989.8983 990.4064
993.1282 1031.4608 1032.2090
1033.9254 1067.1228 1067.1989
1067.3717 1101.4182 1140.4622
1141.7556 1147.2429 1153.4830
1154.4029 1190.2849 1261.5010
1263.7938 1271.0258 1291.5803
1292.8355 1299.6087 1375.4674
1376.6577 1377.7297 1378.1056
1430.8068 1430.8912 1432.3185
1448.3839 1449.1149 1468.4671
1479.1323 1484.7030 1485.3130
1485.3844 1489.6971 1496.6032
1497.0625 1501.9609 1533.7308
1537.0450 1545.0292 1642.8204
1643.4447 1647.4754 2690.7465
3045.9366 3046.0105 3049.3778
3076.6224 3102.2723 3102.3475
3106.5953 3143.2128 3144.2744
3144.4609 3171.5723 3174.3975
3276.3248 3280.9251 3282.8168
3283.0557 3285.2751 3288.1834
3377.3964 3650.1079 3650.5571

**[Zn(Im)$_3$(CH$_3$SO)]$^+$ Complexed with H$_2$O**

---------------------------------------
Gaussian 16: ES64L-G16RevB.01 20-Dec-2017
---------------------------------------
### B3LYP/GEN PSEUDO=READ gfprint gfinput scf=(direct,tight,maxcycle=300
,xqc) opt=(maxcycle=250) freq=noraman
EDU\22-Jun-2020\0\\# B3LYP/GEN PSEUDO=READ gfprint gfinput scf=(direct
,tight,maxcycle=300,xqc) opt=(maxcycle=250) freq=noraman\\zn_3imid_so_
---------------------------------------
Full point group C1 NOp 1
Stoichiometry C13H23N6O2SZn(1+) Framework group C1[X(C13H23N6O2SZn)]
---------------------------------------
Num atoms: 46
Charge = 1 Multiplicity = 1
---------------------------------------
SCF = -1613.49930030 |  Predicted change in Energy=-9.547038D-09

Optimization completed.
Maximum Force            0.000012     0.000450     YES
RMS     Force            0.000002     0.000300     YES
Maximum Displacement     0.001274     0.001800     YES
RMS     Displacement     0.000265     0.001200     YES

---------------------------------------
Atom     Coordinates (in Angstroms)
Type       X          Y          Z
---------------------------------------
Zn   0.039779   0.046792   0.410612
N  -1.746754  -0.911578   0.169975
C  -2.232958  -1.559485  -0.958875
C  -3.464093  -2.099953  -0.691006
C  -2.666359  -1.054510   1.119911
N   1.487015  -1.138656  -0.468462
C   1.975905  -1.186951  -1.764061
C   2.935328  -2.163402  -1.866136
C   2.135322  -2.070321   0.215465
N  -0.055628   1.806396  -0.634820
C  -0.991523   2.181758  -1.587953
C  -0.758768   3.473257  -1.987243
C   0.738700   2.853432  -0.450561
O   0.499976   0.185961   2.273098
S   1.518365   1.432675   2.756934
H  -2.589601  -0.692224   2.142681
H  -1.667534  -1.608874  -1.877540
H   1.556753   2.899152   0.254571
H  -1.773644   1.510869  -1.909819
H   1.997659  -2.290210   1.264026
H   1.619288  -0.516069  -2.531274
C   3.777551  -2.629843  -3.007643
H   4.845846  -2.498945  -2.801469
H   3.603637  -3.688088  -3.232929
H   3.538331  -2.054233  -3.904419
C  -4.417714  -2.891359  -1.524244
H  -4.601542  -3.882878  -1.095088
H  -5.383758  -2.383965  -1.626176
H  -4.007528  -3.032937  -2.526607
C  -1.454299   4.364006  -2.963221
H  -0.783965   4.684431  -3.768757
H  -2.292858   3.832421  -3.418152
H  -1.852211   5.261689  -2.476717
H  -4.532003  -2.015075   1.164788
N  -3.710408  -1.762985   0.631754
H   0.782597   4.784153  -1.285413
N   0.345996   3.872660  -1.249393
H   3.630637  -3.458380  -0.309019
N   3.014696  -2.708265  -0.593165

C 2.963032 0.504294 3.349540
H 3.655318 1.242956 3.768092
H 2.667487 -0.191739 4.139143
H 3.461906 -0.032081 2.536953
H -1.680540 0.240984 4.665355
O -1.646374 -0.267818 3.847546
H -0.784063 -0.060062 3.408720
---------------------------------------
Statistical Thermodynamic Analysis
Temperature 298.150 Kelvin. Pressure 1.00000 Atm.
---------------------------------------
SCF = -1613.49930030 | Predicted change in Energy=-9.547038D-09
Zero-point correction (ZPE) = -1613.1292443 0.370056
Internal Energy (U) = -1613.0998283 0.399472
Enthalpy (H) = -1613.0988833 0.400417
Gibbs Free Energy (G) = -1613.1960443 0.303256
---------------------------------------
Frequencies
13.3922 17.5630 21.6104
26.9389 28.4019 34.7417
40.5663 47.6313 58.3653
68.2754 77.5078 95.0059
107.6993 114.3731 121.2455
127.9975 129.1271 135.7150
147.1760 151.7237 158.2553
170.5344 182.4289 194.1557
195.3745 220.0388 231.8894
242.6347 267.2118 269.0852
272.7512 304.0915 344.3865
345.1295 348.6229 472.2259
528.8422 602.7059 604.7893
612.1301 656.0723 657.2169
659.3266 667.0595 667.2682
668.1244 675.5334 676.4928
680.2810 693.8013 765.4890
797.2819 814.2617 823.4597
827.8925 838.7317 852.2617
940.1708 969.8477 969.9949
971.2812 972.4640 974.9153
990.1575 990.7055 991.9743
1031.7191 1033.1964 1036.2329
1066.9620 1067.1863 1067.2296
1139.7163 1140.6430 1145.2818
1153.9444 1155.2568 1172.8996
1262.2737 1262.4557 1275.1843
1290.5439 1292.2441 1311.4099
1350.0857 1374.7128 1377.2154
1378.5321 1430.6058 1430.8072
1431.2958 1447.4764 1448.2701
1451.2241 1452.7560 1484.9352
1485.1863 1485.2777 1485.3957
1496.6429 1496.9512 1497.5828
1536.7713 1540.4228 1552.1911
1641.7614 1643.9295 1646.1793
1664.4672 3037.3229 3045.3459
3046.5225 3046.5761 3101.2820
3103.1347 3103.2012 3123.6994
3134.5251 3142.9398 3144.9946
3145.1624 3176.6450 3274.7625
3279.1071 3282.1130 3285.4761
3290.6987 3373.0304 3648.0415
3648.8712 3649.5011 3893.4358

**$[Zn(Im)_3(OH^-)]^+$ Complexed with $CH_3SOH$**
---------------------------------------
Gaussian 16: ES64L-G16RevB.01 20-Dec-2017
---------------------------------------
### B3LYP/GEN PSEUDO=READ gfprint gfinput scf=(direct,tight,maxcycle=300
,xqc) opt=(maxcycle=250) freq=noraman
EDU\25-Jun-2020\0\\# B3LYP/GEN PSEUDO=READ gfprint gfinput scf=(direct
,tight,maxcycle=300,xqc) opt=(maxcycle=250) freq=noraman\\zn_3_imid_oh
---------------------------------------
Full point group C1 NOp 1
Stoichiometry C13H23N6O2SZn(1+) Framework group C1[X(C13H23N6O2SZn)]
---------------------------------------
Num atoms: 46
Charge = 1 Multiplicity = 1
---------------------------------------
SCF = -1613.49758399 | Predicted change in Energy=-8.687966D-09

Optimization completed on the basis of negligible forces.
Maximum Force 0.000004 0.000450 YES
RMS Force 0.000001 0.000300 YES
Maximum Displacement 0.002965 0.001800 NO
RMS Displacement 0.000671 0.001200 YES

---------------------------------------
Atom Coordinates (in Angstroms)
Type X Y Z
---------------------------------------
Zn 0.218293 -0.502376 -0.078759
O -2.869421 0.600792 -1.848029
N 1.026268 -2.105920 0.933096
C 2.300149 -2.629871 0.776638
C 2.427995 -3.778941 1.516079
C 0.381216 -2.922567 1.754714
N 1.750609 0.659172 -0.822255
C 2.790364 1.396366 -0.277524
C 3.453887 2.071303 -1.271245
C 1.771967 0.879421 -2.130498
S -4.248877 -0.247658 -2.340333
H 1.072715 0.450507 -2.834818
H 2.992450 1.401774 0.783125
H -0.641070 -2.814589 2.085945
H 3.033251 -2.152505 0.144177
C 3.563209 -4.730128 1.706164
H 3.303645 -5.739931 1.369079
H 4.426352 -4.396915 1.126092
H 3.868758 -4.791240 2.756720
C 4.625821 2.996490 -1.248464
H 5.457858 2.611728 -1.849059
H 4.363743 3.989462 -1.631003
H 4.982650 3.118128 -0.223400
H 3.008509 2.051253 -3.366336
N 2.784800 1.723397 -2.436288
H 0.935048 -4.697286 2.745510
N 1.193585 -3.939713 2.127241
C -4.371266 0.276164 -4.075852

H -5.293024 -0.171837 -4.463072
H -4.447854 1.364147 -4.142473
H -3.523644 -0.083816 -4.664375
H -2.544802 1.109400 0.085982
C -1.537648 1.626323 3.169870
C -0.553112 0.841887 2.625924
N -0.862069 0.546747 1.304492
C -2.020797 1.143263 1.039306
N -2.451813 1.799453 2.140970
H -3.311690 2.329197 2.194499
H 0.347550 0.476896 3.096723
C -1.712481 2.223794 4.527288
H -1.757296 3.317932 4.483983
H -2.629173 1.867862 5.011081
H -0.869891 1.949890 5.166071
O -0.719324 -0.876923 -1.697563
H -2.073289 -0.016253 -1.899687
H -0.902148 -1.807278 -1.873736
---------------------------------------
Statistical Thermodynamic Analysis
Temperature 298.150 Kelvin. Pressure 1.00000 Atm.
---------------------------------------
SCF = -1613.49758399 | Predicted change in Energy=-8.687966D-09
Zero-point correction (ZPE) = -1613.12707799 0.370506
Internal Energy (U) = -1613.09813599 0.399448
Enthalpy (H) = -1613.09719099 0.400393
Gibbs Free Energy (G) = -1613.19453699 0.303047
---------------------------------------
Frequencies
8.4988 16.2914 20.8448
24.2334 26.1036 29.8429
39.9268 42.6972 63.4103
71.6182 79.0621 90.1650
96.9872 110.1018 113.1419
121.2317 127.9690 132.1128
147.6172 153.0054 165.9933
174.9009 182.9679 195.1944
227.9638 236.1255 267.7887
269.4149 274.3147 298.8342
307.9639 344.4892 345.1890
349.6198 396.4446 515.3076
603.7615 604.9981 614.2920
655.4528 656.9334 660.0207
666.2903 668.0694 668.3206
675.2059 677.7704 682.3741
690.9474 761.4531 781.5246
812.9238 819.5990 821.5885
837.1253 873.4489 946.1357
966.1428 966.9724 971.0024
972.9203 978.1202 989.3391
989.7656 990.3502 993.8650
1031.6917 1032.4337 1035.6765
1066.9192 1067.1443 1067.5183
1138.6375 1140.0914 1145.2676
1151.0150 1156.2115 1171.0823
1259.4859 1261.1760 1274.9636
1290.5567 1291.2559 1309.0363
1355.3631 1375.2130 1377.0096
1377.9403 1430.5048 1430.6345
1431.1655 1446.8025 1448.1653
1449.9093 1453.7449 1480.7274
1485.1558 1485.2610 1485.3680
1485.5003 1496.3257 1496.9534
1497.7589 1532.2917 1541.9988
1549.2324 1641.8784 1643.0593
1644.8540 3023.7361 3045.2362
3045.5440 3045.7463 3047.5052
3101.5930 3101.8882 3104.6217
3133.0375 3142.9966 3144.1293
3144.1675 3146.7038 3168.5180
3279.6367 3280.6568 3283.6674
3286.6139 3290.1044 3646.9040
3648.0602 3649.8421 3875.1743

**$[Zn(Im)_3(CH_3SO)]^+$ Complexed with HOCl**

---------------------------------------
Gaussian 16: ES64L-G16RevB.01 20-Dec-2017
---------------------------------------
### B3LYP/GEN PSEUDO=READ gfprint gfinput scf=(direct,tight,maxcycle=300 ,xqc) opt=(maxcycle=250) freq=noraman AN.EDU\23-Jun-2020\0\\# B3LYP/GEN PSEUDO=READ gfprint gfinput scf=(dir ect,tight,maxcycle=300,xqc) opt=(maxcycle=250) freq=noraman\\zn_3im_oc
---------------------------------------
Full point group C1 NOp 1
Stoichiometry C13H22ClN6O2SZn(1+) Framework group C1[X(C13H22ClN6O2SZn)]
---------------------------------------
Num atoms: 46
Charge = 1 Multiplicity = 1
---------------------------------------
SCF = -2073.02794243 | Predicted change in Energy=-4.996238D-09

Optimization completed.

| | | | |
|---|---|---|---|
| Maximum Force | 0.000010 | 0.000450 | YES |
| RMS Force | 0.000002 | 0.000300 | YES |
| Maximum Displacement | 0.001747 | 0.001800 | YES |
| RMS Displacement | 0.000323 | 0.001200 | YES |

---------------------------------------
Atom Coordinates (in Angstroms)
Type X Y Z
---------------------------------------
Zn -0.395250 -0.206739 -0.268946
N -0.204993 -1.771443 1.008300
C -1.155886 -2.377103 1.818194
C -0.598687 -3.450589 2.463861
C 0.920808 -2.464232 1.152567
N -2.368977 0.219495 -0.560358
C -3.247708 0.877625 0.288947
C -4.470123 1.013318 -0.318321
C -3.043274 -0.042663 -1.673220
N 0.654263 1.420845 0.373366
C 0.807872 2.005186 1.621881
C 1.724795 3.022744 1.548813
C 1.468146 2.067466 -0.452540
H 1.854647 -2.262067 0.637548
H -2.170420 -2.012625 1.875752
H 1.609251 1.837804 -1.498527

H 0.263615 1.656601 2.486691
H -2.646392 -0.543515 -2.544327
H -2.947313 1.219274 1.268021
C -5.751375 1.634700 0.130744
H -6.050570 2.459713 -0.525410
H -6.568755 0.905201 0.151863
H -5.635710 2.036195 1.139730
C -1.156486 -4.441894 3.431230
H -1.097297 -5.463875 3.040148
H -0.625462 -4.415168 4.389472
H -2.207990 -4.220023 3.626002
C 2.261196 3.967021 2.573489
H 2.044344 5.009668 2.315167
H 1.803865 3.761212 3.543690
H 3.346338 3.864657 2.686754
H 1.416987 -4.156423 2.292082
N 0.716443 -3.478677 2.022367
H 2.818761 3.653721 -0.181812
N 2.123894 3.038880 0.220373
H -5.020216 0.344835 -2.276971
N -4.309378 0.419611 -1.561283
O 0.605666 -0.467886 -1.931797
Cl -0.120316 -1.518599 -3.087318
H 2.126742 -1.056747 -1.482217
O 2.907615 -1.408914 -0.972733
S 4.244228 -0.399307 -1.217165
C 5.002918 -1.172424 -2.675663
H 4.352605 -1.099878 -3.550568
H 5.927313 -0.616036 -2.866092
H 5.250265 -2.215947 -2.467786
---------------------------------------
Statistical Thermodynamic Analysis
Temperature 298.150 Kelvin. Pressure 1.00000 Atm.
---------------------------------------
SCF = -2073.02794243 | Predicted change in Energy=-4.996238D-09
Zero-point correction (ZPE) = -2072.66676043 0.361182
Internal Energy (U) = -2072.63650543 0.391437
Enthalpy (H) = -2072.63556143 0.392381
Gibbs Free Energy (G) = -2072.73575543 0.292187
---------------------------------------
Frequencies
17.0061 20.0241 20.5617
21.7399 27.4792 36.0101
37.9520 44.9557 48.1661
58.7626 65.1414 79.0415
94.1102 96.1904 101.9917
111.6396 121.2776 123.5266
128.2413 135.0620 144.7869
152.9647 158.8944 180.1448
183.0602 200.8411 241.7358
244.4691 266.7773 269.5826
272.8631 277.0048 298.1680
344.0996 347.1591 349.7472
430.3657 606.0710 608.2454
612.9177 654.5853 657.6643
658.5383 667.5133 668.4968
668.7263 674.2855 675.9845
680.2132 690.2423 739.0119
761.5875 814.6718 817.9284
821.9214 853.6123 855.6207
918.9276 919.9089 972.3898
973.1732 973.9154 977.9157
983.0869 990.7838 991.3144
991.6366 1032.7822 1034.1125
1035.4213 1067.3813 1067.4762
1067.5568 1140.6174 1141.7381
1146.4422 1157.6652 1159.2792
1167.1140 1263.9166 1264.4518
1273.4650 1292.8468 1294.6384
1301.3615 1356.4509 1367.6351
1375.5624 1376.3725 1378.1266
1431.0786 1431.2373 1431.4316
1449.0861 1450.2513 1450.5547
1453.1250 1484.2831 1485.1585
1485.2657 1485.3150 1496.9405
1497.2737 1497.5365 1538.5305
1541.1505 1546.8228 1644.0997
1644.8317 1646.0562 3046.3107
3046.5803 3047.2918 3047.6449
3102.8388 3103.3768 3104.8397
3135.9332 3144.3901 3144.6899
3146.1141 3150.9697 3210.6847
3248.0918 3282.5731 3283.0736
3284.2765 3286.6726 3293.3780
3647.2339 3648.5254 3650.1553

**$[Zn(Im)_3(OCl)]^+$ Complexed with $CH_3SOH$**

---------------------------------------
Gaussian 16: ES64L-G16RevB.01 20-Dec-2017
---------------------------------------
### B3LYP/GEN PSEUDO=READ gfprint gfinput scf=(direct,tight,maxcycle=300
,xqc) opt=(maxcycle=250) freq=noraman
AN.EDU\23-Jun-2020\0\\# B3LYP/GEN PSEUDO=READ gfprint gfinput scf=(dir
ect,tight,maxcycle=300,xqc) opt=(maxcycle=250) freq=noraman\\zn_3imid_
---------------------------------------
Full point group C1 NOp 1
Stoichiometry C13H22ClN6O2SZn(1+) Framework group C1[X(C13H22ClN6O2SZn)]
---------------------------------------
Num atoms: 46
Charge = 1 Multiplicity = 1
---------------------------------------
SCF = -2073.02256904 | Predicted change in Energy=-1.256886D-08

Optimization completed.

| | | | |
|---|---|---|---|
| Maximum Force | 0.000011 | 0.000450 | YES |
| RMS Force | 0.000002 | 0.000300 | YES |
| Maximum Displacement | 0.001608 | 0.001800 | YES |
| RMS Displacement | 0.000430 | 0.001200 | YES |

---------------------------------------
Atom Coordinates (in Angstroms)
Type X Y Z
---------------------------------------
Zn -0.134185 0.032151 -0.181289
N 1.158074 0.873738 1.160103
C 0.870842 1.568641 2.328098
C 2.035970 1.947671 2.943903

C 2.482728 0.824828 1.063202
N -1.229184 -1.394419 0.816242
C -2.448088 -1.341149 1.473747
C -2.744547 -2.570258 2.007615
C -0.782518 -2.636652 0.944095
N -1.355877 1.487062 -0.922345
C -1.572977 2.764203 -0.423370
C -2.403304 3.459054 -1.265048
C -2.042165 1.401904 -2.055892
O 0.877492 -0.968942 -1.494623
S 0.314323 -1.101752 -3.078058
H 3.043869 0.314454 0.287442
H -0.142986 1.737526 2.659275
H -2.053977 0.546085 -2.715718
H -1.108655 3.104911 0.489363
H 0.146287 -3.015903 0.542118
H -3.033744 -0.434983 1.514174
C -3.910944 -3.069265 2.795235
H -4.429502 -3.883874 2.277215
H -3.605446 -3.435799 3.781719
H -4.628993 -2.260607 2.947577
C 2.308173 2.700832 4.204171
H 2.886703 2.100607 4.915402
H 2.865158 3.624466 4.010413
H 1.366646 2.973155 4.686106
C -2.945960 4.849325 -1.219018
H -4.041304 4.855508 -1.188117
H -2.583542 5.359855 -0.324230
H -2.628202 5.434298 -2.089388
H 4.033404 1.552484 2.277118
N 3.038551 1.462108 2.118322
H -3.263353 2.760399 -3.097653
N -2.685266 2.568217 -2.290267
H -1.558916 -4.349748 1.889590
N -1.668383 -3.371785 1.656063
C -0.011283 -2.885309 -3.187464
H -0.329133 -3.073493 -4.218893
H 0.902794 -3.452138 -2.992587
H -0.807497 -3.196173 -2.505113
H 2.360851 -1.266977 -1.279913
O 3.343116 -1.358274 -1.002551
Cl 3.380952 -2.788979 -0.046770
---------------------------------------
Statistical Thermodynamic Analysis
Temperature 298.150 Kelvin. Pressure 1.00000 Atm.
---------------------------------------
SCF = -2073.02256904 | Predicted change in Energy=-1.256886D-08
Zero-point correction (ZPE) = -2072.66200204 0.360567
Internal Energy (U) = -2072.63177604 0.390793
Enthalpy (H) = -2072.63083204 0.391737
Gibbs Free Energy (G) = -2072.73187004 0.290699
---------------------------------------
Frequencies
10.8497 13.8182 21.2186
23.6653 25.6609 29.0870
35.1017 40.4020 48.4928
60.0463 65.0292 70.6154
94.8737 102.4876 113.9781
115.6082 124.6514 125.7024
128.3540 141.0614 146.6865
149.5500 159.9170 178.7447
183.3800 199.5376 231.2026
243.8069 250.8957 267.2266
269.3435 274.5528 323.0080
344.6279 345.3029 350.4958
483.5528 604.9335 606.8439
613.2828 656.1495 657.1644
659.8101 667.3289 667.5725
668.0164 675.2859 676.2784
679.5846 693.3347 720.5321
779.4953 812.9580 821.2716
823.1941 844.7629 845.7226
910.1257 971.2344 972.5063
974.0017 974.1967 975.8475
990.3564 990.9671 991.5207
1032.2995 1033.4159 1036.0848
1051.0524 1067.2747 1067.3669
1067.3863 1140.4126 1141.1863
1146.4547 1156.5717 1157.9512
1172.0878 1262.8475 1264.6724
1274.0501 1292.2312 1293.4241
1304.5064 1353.8672 1374.4746
1376.4542 1377.6113 1430.8217
1431.1861 1431.5275 1447.9910
1449.0983 1451.1675 1452.8369
1484.3973 1485.0507 1485.0896
1485.3094 1485.5395 1496.7660
1496.9993 1497.5219 1538.0620
1540.9542 1549.4247 1642.5128
1644.6037 1646.2931 2772.8216
3039.7934 3046.6827 3046.8810
3047.4526 3103.4466 3103.6617
3104.5930 3127.1307 3137.7831
3144.0216 3145.1973 3145.9095
3218.4756 3277.6428 3278.7551
3281.2036 3284.2615 3288.8433
3647.1560 3647.5526 3647.8940

**$[Zn(Im)_3(CH_3SO_2)]^+$ Complexed with $H_2O$**

---------------------------------------
Gaussian 16: ES64L-G16RevB.01 20-Dec-2017
---------------------------------------
### B3LYP/GEN PSEUDO=READ gfprint gfinput scf=(direct,tight,maxcycle=300
,xqc) opt=(maxcycle=250) freq=noraman
19-Jul-2020\0\\# B3LYP/GEN PSEUDO=READ gfprint gfinput scf=(direct,tig
ht,maxcycle=300,xqc) opt=(maxcycle=250) freq=noraman\\zn_3im_1meso2_ct
---------------------------------------
Full point group C1 NOp 1
Stoichiometry C13H23N6O3SZn(1+) Framework group C1[X(C13H23N6O3SZn)]
---------------------------------------
Num atoms: 47
Charge = 1 Multiplicity = 1
---------------------------------------
SCF = -1688.72537512 | Predicted change in Energy=-2.773921D-08

Optimization completed.
Maximum Force 0.000024 0.000450 YES
RMS Force 0.000005 0.000300 YES

Maximum Displacement 0.001519 0.001800 YES
RMS Displacement 0.000373 0.001200 YES

---------------------------------------
Atom Coordinates (in Angstroms)
Type X Y Z
---------------------------------------
Zn 0.079887 0.345954 -0.397100
N 1.347819 1.906330 0.019627
C 2.718119 1.839451 0.222003
C 3.226044 3.100929 0.396779
C 1.017248 3.191263 0.068088
N 1.293768 -1.290428 -0.517315
C 1.907962 -1.973187 0.526332
C 2.691556 -2.981117 0.025661
C 1.698948 -1.869152 -1.641434
N -1.167793 -0.008562 1.214437
C -1.341085 -1.172680 1.949167
C -2.343171 -1.001201 2.869763
C -2.055549 0.864554 1.674615
H 0.017911 3.590999 -0.058945
H 3.246182 0.898258 0.217201
H -2.215218 1.865788 1.292279
H -0.752176 -2.057289 1.761577
H 1.408503 -1.542815 -2.631844
H 1.758404 -1.693595 1.558515
C 3.556032 -4.004414 0.685573
H 4.600296 -3.919376 0.364242
H 3.528639 -3.871573 1.769331
H 3.217099 -5.022874 0.464358
C 4.611973 3.602436 0.636505
H 4.687265 4.149406 1.583223
H 5.308013 2.761934 0.680662
H 4.944602 4.269268 -0.167094
C -2.934215 -1.913205 3.893898
H -2.814028 -1.515897 4.908203
H -4.003667 -2.077203 3.719562
H -2.438154 -2.885500 3.855680
H 2.138461 4.945301 0.368155
N 2.123803 3.936969 0.295821
H -3.519217 0.760518 3.182040
N -2.776216 0.301215 2.672859
H 2.981369 -3.485561 -2.036290
N 2.536832 -2.890212 -1.350253
H -1.948325 2.531981 -1.034022
H -2.785284 3.840750 -0.789817
O -2.110439 3.271223 -0.401471
O -1.262101 1.024613 -1.687016
C -2.060782 -0.275008 -3.816521
H -1.860266 -0.483841 -4.870457
H -1.942333 -1.178777 -3.213320
H -3.062982 0.143367 -3.695871
S -0.832303 0.951952 -3.244496
O 0.492569 0.204009 -3.284890
---------------------------------------
Statistical Thermodynamic Analysis
Temperature 298.150 Kelvin. Pressure 1.00000 Atm.
---------------------------------------
SCF = -1688.72537512 | Predicted change in Energy=-2.773921D-08
Zero-point correction (ZPE) = -1688.35041912 0.374956
Internal Energy (U) = -1688.32064212 0.404733
Enthalpy (H) = -1688.31969812 0.405677
Gibbs Free Energy (G) = -1688.41543912 0.309936
---------------------------------------
Frequencies
19.0761 22.4559 28.0222
31.3696 34.9509 42.4936
49.3681 54.9533 62.8859
78.5706 105.0040 106.8764
114.8315 119.1057 121.0289
127.1684 130.1092 136.8823
141.1815 149.5692 160.1924
171.3359 185.4875 194.4715
202.1662 228.7381 236.0262
266.4241 272.9603 273.4966
277.8983 287.6483 337.8521
345.2558 347.4945 353.6479
413.9950 496.6375 531.0025
601.3540 604.6320 609.8532
655.5070 657.3281 659.8328
661.4196 666.4769 667.4295
668.2444 676.2167 679.3283
680.3611 764.9874 815.4604
825.6199 827.9123 835.5413
897.6835 905.7014 913.5765
943.6936 962.0648 969.7541
971.4595 973.1449 990.5686
991.4514 991.5431 1032.9937
1034.9727 1036.9685 1052.3602
1066.9900 1067.1827 1067.2554
1136.8519 1142.6673 1147.4024
1154.6506 1163.3976 1167.0947
1259.6905 1271.7503 1273.5770
1290.2167 1301.5381 1303.7461
1332.4934 1374.1035 1378.0967
1379.9442 1430.7582 1430.9626
1431.5689 1447.9295 1448.8285
1449.8170 1451.1135 1458.0745
1485.2371 1485.3967 1485.5092
1496.3438 1497.4114 1497.6783
1531.4345 1543.9290 1549.1944
1642.1684 1644.0118 1645.3504
1646.4457 3045.1509 3045.4208
3045.8289 3054.3108 3101.1071
3101.4503 3102.0242 3143.7025
3143.7125 3143.9084 3163.0366
3164.2290 3233.5632 3239.4477
3258.5272 3280.9708 3288.0159
3288.6557 3445.1034 3649.1580
3650.7863 3651.5218 3884.9295

**$[Zn(Im)_3(OH)]^+$ Complexed with $CH_3SO_2H$**

---------------------------------------
Gaussian 16: ES64L-G16RevB.01 20-Dec-2017
---------------------------------------
### B3LYP/GEN PSEUDO=READ gfprint gfinput scf=(direct,tight,maxcycle=300 ,xqc) OPT freq=noraman
17-Jul-2020\0\\# B3LYP/GEN PSEUDO=READ gfprint gfinput scf=(direct,tig ht,maxcycle=300,xqc) OPT freq=noraman\\zn_3im_1oh_ctr_meso2h.xyz\\1,1\
---------------------------------------

Full point group C1 NOp 1
Stoichiometry C13H23N6O3SZn(1+) Framework group C1[X(C13H23N6O3SZn)]
---------------------------------------
Num atoms: 47
Charge = 1 Multiplicity = 1
---------------------------------------
SCF = -1688.71555329 | Predicted change in Energy=-7.726444D-08

Optimization completed.

| | | | |
|---|---|---|---|
| Maximum Force | 0.000073 | 0.000450 | YES |
| RMS Force | 0.000010 | 0.000300 | YES |
| Maximum Displacement | 0.001411 | 0.001800 | YES |
| RMS Displacement | 0.000410 | 0.001200 | YES |

---------------------------------------

| Atom Type | Coordinates (in Angstroms) X | Y | Z |
|---|---|---|---|
| Zn | 0.028014 | 0.091331 | -0.593716 |
| N | -0.902736 | 1.592559 | 0.426031 |
| C | -0.381656 | 2.550207 | 1.286222 |
| C | -1.393031 | 3.339478 | 1.770630 |
| C | -2.217369 | 1.787810 | 0.384721 |
| N | 2.027296 | 0.581057 | -0.753812 |
| C | 3.108376 | -0.276271 | -0.617699 |
| C | 4.269738 | 0.378176 | -0.944301 |
| C | 2.518208 | 1.743623 | -1.161365 |
| N | -0.148370 | -1.618183 | 0.530223 |
| C | 0.475195 | -1.999930 | 1.709505 |
| C | -0.026597 | -3.203829 | 2.134041 |
| C | -1.024462 | -2.571831 | 0.236377 |
| H | -2.943309 | 1.200636 | -0.175797 |
| H | 0.673996 | 2.606346 | 1.506240 |
| H | -1.701452 | -2.570620 | -0.609700 |
| H | 1.231349 | -1.388576 | 2.179103 |
| H | 1.947089 | 2.634471 | -1.377927 |
| H | 2.980820 | -1.301704 | -0.305176 |
| C | 5.696071 | -0.062438 | -0.975803 |
| H | 6.122708 | 0.025100 | -1.981269 |
| H | 6.316912 | 0.526463 | -0.291333 |
| H | 5.766756 | -1.109552 | -0.674042 |
| C | -1.400728 | 4.498536 | 2.712283 |
| H | -1.793408 | 5.404179 | 2.236108 |
| H | -2.009818 | 4.293560 | 3.600025 |
| H | -0.383668 | 4.712354 | 3.048461 |
| C | 0.278927 | -4.058645 | 3.319695 |
| H | 0.642731 | -5.048753 | 3.022346 |
| H | 1.053889 | -3.587091 | 3.927976 |
| H | -0.603745 | -4.199472 | 3.953777 |
| H | -3.482855 | 3.173609 | 1.321192 |
| N | -2.541404 | 2.831337 | 1.182363 |
| H | -1.549869 | -4.372695 | 1.185057 |
| N | -0.975780 | -3.540341 | 1.180318 |
| H | 4.471197 | 2.410569 | -1.585791 |
| N | 3.864105 | 1.659645 | -1.284729 |
| H | -1.971965 | -1.099440 | -2.418820 |
| O | -0.788553 | -0.245235 | -2.291581 |
| H | -0.225724 | -0.190817 | -3.071593 |
| O | -2.834025 | -1.684639 | -2.334631 |
| C | -5.362281 | -2.000490 | -2.386886 |
| H | -6.349311 | -1.535277 | -2.344529 |
| H | -5.239845 | -2.565278 | -3.313340 |
| H | -5.199985 | -2.634614 | -1.512360 |
| S | -4.131497 | -0.663844 | -2.340666 |
| O | -4.272134 | -0.049681 | -0.962031 |

---------------------------------------
Statistical Thermodynamic Analysis
Temperature 298.150 Kelvin. Pressure 1.00000 Atm.
---------------------------------------
SCF = -1688.71555329 | Predicted change in Energy=-7.726444D-08
Zero-point correction (ZPE) = -1688.34165629 0.373897
Internal Energy (U) = -1688.31204429 0.403509
Enthalpy (H) = -1688.31110029 0.404453
Gibbs Free Energy (G) = -1688.40815529 0.307398
---------------------------------------
Frequencies
12.6249 19.8999 21.4890
26.8104 31.0195 42.0202
44.7521 48.9179 67.1805
76.4538 89.7107 96.2137
98.8054 104.2449 116.9865
119.3659 127.9136 131.8498
145.7638 153.0665 158.4278
173.8689 184.9477 189.1436
221.8374 235.3290 240.0554
266.0757 267.9370 273.2310
281.7805 342.1975 346.6834
347.7606 351.9326 383.0130
449.5080 500.0074 604.8342
605.7847 613.6562 656.1269
657.6889 660.7896 666.1408
667.6748 668.7164 676.1350
677.7479 679.7439 682.4165
774.4730 786.7854 810.0875
818.1578 820.1880 838.7045
897.6858 955.1065 963.1184
969.3200 970.7275 972.9813
975.9526 990.5750 990.9893
991.0401 1032.1925 1034.2167
1035.2167 1067.0772 1067.1830
1067.5798 1068.6063 1138.8966
1142.4283 1145.8746 1156.3136
1162.5391 1172.4183 1216.7846
1260.9329 1274.2029 1275.9420
1292.1023 1301.7856 1308.8620
1340.8905 1372.5836 1376.0127
1377.9576 1430.6449 1430.9889
1431.1814 1446.2114 1446.6039
1451.0451 1451.2839 1457.6245
1460.9212 1485.1625 1485.2911
1485.4031 1496.7786 1497.2990
1497.7451 1541.0358 1546.2548
1554.0001 1642.7421 1644.5204
1645.5865 2414.1980 3044.9877
3045.5641 3047.7500 3064.3812
3100.8052 3101.7064 3104.9892
3141.9903 3143.5998 3146.9975
3155.7660 3173.1521 3176.2525
3241.0911 3279.4002 3280.2752
3285.6302 3288.9056 3646.6653
3649.3284 3650.5967 3888.8120

**[Zn(Im)$_3$(CH$_3$SH)]$^{2+}$ Complexed with $H_2O$**

---------------------------------------
Gaussian 16: ES64L-G16RevB.01 20-Dec-2017
---------------------------------------
### B3LYP/GEN PSEUDO=READ gfprint gfinput scf=(direct,tight,maxcycle=300 ,xqc) opt=(maxcycle=250) freq=noraman EDU\23-Jun-2020\0\\# B3LYP/GEN PSEUDO=READ gfprint gfinput scf=(direct ,tight,maxcycle=300,xqc) opt=(maxcycle=250) freq=noraman\\zn_3imid_thi

---------------------------------------
Full point group C1 NOp 1
Stoichiometry C13H24N6OSZn(2+) Framework group C1[X(C13H24N6OSZn)]

---------------------------------------
Num atoms: 46
Charge = 2 Multiplicity = 1

---------------------------------------
SCF = -1538.59956533 | Predicted change in Energy=-1.034109D-08

Optimization completed.

| | | | |
|---|---|---|---|
| Maximum Force | 0.000016 | 0.000450 | YES |
| RMS Force | 0.000002 | 0.000300 | YES |
| Maximum Displacement | 0.001328 | 0.001800 | YES |
| RMS Displacement | 0.000364 | 0.001200 | YES |

---------------------------------------

| Atom Type | Coordinates (in Angstroms) X | Y | Z |
|---|---|---|---|
| Zn | -0.062921 | 0.034114 | -0.291661 |
| N | 0.739985 | 1.810257 | 0.241745 |
| C | 1.805733 | 1.952575 | 1.125272 |
| C | 2.126442 | 3.279820 | 1.255908 |
| C | 0.415105 | 3.037881 | -0.163992 |
| N | 1.345020 | -1.419627 | -0.355272 |
| C | 1.620931 | -2.345101 | 0.646149 |
| C | 2.672881 | -3.140782 | 0.267058 |
| C | 2.220240 | -1.649018 | -1.332260 |
| N | -1.528034 | -0.528325 | 0.985802 |
| C | -2.170261 | -1.761268 | 1.039524 |
| C | -3.091006 | -1.762700 | 2.057367 |
| C | -2.051473 | 0.211903 | 1.960574 |
| S | -0.838286 | 0.245474 | -2.605665 |
| C | -2.374325 | -0.750714 | -2.811919 |
| H | -2.829779 | -0.479061 | -3.764548 |
| H | -2.070594 | -1.797620 | -2.848028 |
| H | -3.067721 | -0.584871 | -1.988138 |
| H | -0.367445 | 3.298845 | -0.865389 |
| H | 2.271973 | 1.101777 | 1.598612 |
| H | -1.775940 | 1.226090 | 2.208488 |
| H | -1.927946 | -2.562545 | 0.357860 |
| H | 2.292606 | -1.111086 | -2.266070 |
| H | 1.050390 | -2.381415 | 1.561752 |
| C | 3.368823 | -4.275196 | 0.943394 |
| H | 4.431499 | -4.058811 | 1.097219 |
| H | 2.919384 | -4.459080 | 1.921189 |
| H | 3.291715 | -5.198246 | 0.358692 |
| C | 3.170556 | 3.986482 | 2.055392 |
| H | 2.724531 | 4.695119 | 2.761592 |
| H | 3.753709 | 3.263757 | 2.629384 |
| H | 3.861987 | 4.539733 | 1.410622 |
| C | -4.039143 | -2.804066 | 2.552537 |
| H | -3.841400 | -3.060745 | 3.598871 |
| H | -5.078890 | -2.467985 | 2.476448 |
| H | -3.937002 | -3.714666 | 1.959008 |
| H | 1.191010 | 4.939099 | 0.282937 |
| N | 1.228627 | 3.937451 | 0.428690 |
| H | -3.531765 | -0.161448 | 3.406334 |
| N | -2.990516 | -0.498897 | 2.619149 |
| H | 3.775333 | -3.043425 | -1.564595 |
| N | 3.027802 | -2.673734 | -0.989236 |
| H | -1.387210 | 1.491095 | -2.593994 |
| H | -1.598891 | 3.908252 | -3.396097 |
| O | -1.875518 | 3.443658 | -2.592261 |
| H | -2.780214 | 3.746049 | -2.425440 |

---------------------------------------
Statistical Thermodynamic Analysis
Temperature 298.150 Kelvin. Pressure 1.00000 Atm.

---------------------------------------
SCF = -1538.59956533 | Predicted change in Energy=-1.034109D-08
Zero-point correction (ZPE) = -1538.22281333 0.376752
Internal Energy (U) = -1538.19381833 0.405747
Enthalpy (H) = -1538.19287433 0.406691
Gibbs Free Energy (G) = -1538.28848833 0.311077

---------------------------------------
Frequencies
14.3361 20.1905 26.4278
29.6068 33.2147 35.1238
40.1422 48.5589 56.9465
77.0457 89.9163 99.8411
102.0467 109.9484 120.2314
123.8123 127.5584 132.9357
141.7188 145.6519 156.1914
164.8831 177.8754 196.7711
234.1811 251.3276 256.1645
260.3909 272.6369 274.4911
282.7643 324.1059 347.5586
349.4001 350.7439 421.9828
628.3778 629.1977 633.6652
656.3456 661.4078 662.4784
664.4730 665.0980 665.7953
666.4180 684.2453 686.8090
687.4594 688.1443 806.7284
812.1163 818.8809 827.2288
832.1467 860.1775 881.5010
975.9181 977.7743 978.3591
991.4876 991.6831 992.4225
994.2819 1032.7064 1033.5692
1035.2025 1066.5344 1066.7668
1066.8843 1123.9116 1133.0697
1135.4984 1141.3509 1166.1942
1167.7177 1171.1692 1257.3239
1259.1684 1266.7496 1293.6714
1294.1475 1299.9586 1367.8218
1369.3060 1370.6445 1377.5738
1432.1440 1432.2993 1432.4292
1446.4007 1446.8349 1447.1385
1477.2364 1480.9758 1483.1038
1483.3980 1483.4669 1495.1483

1495.2781 1495.6250 1542.5596
1545.5434 1550.1107 1643.1503
1644.0841 1645.2634 1645.8685
2531.0863 3053.7953 3054.0587
3054.1838 3086.5004 3114.2555
3114.7681 3115.0033 3152.9317
3153.0472 3153.1005 3186.9438
3194.7985 3251.2585 3286.2656
3287.4365 3287.8934 3290.6652
3293.0453 3628.7261 3629.0907
3631.7107 3781.5380 3880.5448

**$[Zn(Im)_3(H_2O)]^{2+}$ Complexed with $CH_3SH$**

---------------------------------------
Gaussian 16: ES64L-G16RevB.01 20-Dec-2017
---------------------------------------
### B3LYP/GEN PSEUDO=READ gfprint gfinput scf=(direct,tight,maxcycle=300 ,xqc) opt=(maxcycle=250) freq=noraman
EDU\22-Jun-2020\0\\# B3LYP/GEN PSEUDO=READ gfprint gfinput scf=(direct ,tight,maxcycle=300,xqc) opt=(maxcycle=250) freq=noraman\\zn_3imid_h2o
---------------------------------------
Full point group C1 NOp 1
Stoichiometry C13H24N6OSZn(2+) Framework group C1[X(C13H24N6OSZn)]
---------------------------------------
Num atoms: 46
Charge = 2 Multiplicity = 1
---------------------------------------
SCF = -1538.60186892 | Predicted change in Energy=-8.445831D-09

Optimization completed.

| | | | |
|---|---|---|---|
| Maximum Force | 0.000013 | 0.000450 | YES |
| RMS Force | 0.000002 | 0.000300 | YES |
| Maximum Displacement | 0.001420 | 0.001800 | YES |
| RMS Displacement | 0.000343 | 0.001200 | YES |

---------------------------------------
Atom Coordinates (in Angstroms)

| Type | X | Y | Z |
|---|---|---|---|
| Zn | 0.084581 | 0.297406 | -0.067020 |
| N | -0.814295 | 1.900270 | 0.740895 |
| C | -0.875904 | 2.221093 | 2.093183 |
| C | -1.580110 | 3.386643 | 2.261256 |
| C | -1.474104 | 2.858597 | 0.092546 |
| N | 2.002147 | 0.576353 | -0.601427 |
| C | 2.850819 | -0.367197 | -1.173594 |
| C | 4.083234 | 0.192921 | -1.398127 |
| C | 2.711450 | 1.697067 | -0.478126 |
| N | -0.179847 | -1.410642 | 0.948962 |
| C | 0.794876 | -2.039485 | 1.718743 |
| C | 0.263341 | -3.150480 | 2.322845 |
| C | -1.292274 | -2.131017 | 1.084311 |
| H | -1.625512 | 2.916021 | -0.974995 |
| H | -0.421193 | 1.599036 | 2.849343 |
| H | -2.246243 | -1.933933 | 0.617179 |
| H | 1.802365 | -1.657954 | 1.786596 |
| H | 2.361335 | 2.631851 | -0.066133 |
| H | 2.526661 | -1.376042 | -1.380795 |
| C | 5.343674 | -0.353903 | -1.981716 |
| H | 5.640597 | 0.197454 | -2.880381 |
| H | 6.170510 | -0.305553 | -1.264960 |
| H | 5.202706 | -1.399764 | -2.261234 |
| C | -1.946062 | 4.171843 | 3.476967 |
| H | -1.534161 | 5.186041 | 3.438195 |
| H | -3.032657 | 4.250727 | 3.590486 |
| H | -1.550393 | 3.683286 | 4.369481 |
| C | 0.849942 | -4.172158 | 3.239658 |
| H | 0.799699 | -5.176346 | 2.805084 |
| H | 1.899653 | -3.942059 | 3.432005 |
| H | 0.328424 | -4.192208 | 4.202660 |
| H | -2.481085 | 4.587431 | 0.736360 |
| N | -1.943466 | 3.762967 | 0.976573 |
| H | -1.742724 | -3.877465 | 2.160732 |
| N | -1.056793 | -3.179077 | 1.899569 |
| H | 4.694102 | 2.196869 | -0.962104 |
| N | 3.960143 | 1.498431 | -0.946530 |
| H | -4.534374 | -0.518831 | -2.348870 |
| S | -3.586916 | -1.444567 | -2.103194 |
| C | -3.778070 | -2.447860 | -3.635975 |
| H | -3.037196 | -3.246681 | -3.579723 |
| H | -4.776280 | -2.885158 | -3.657684 |
| H | -3.602860 | -1.841446 | -4.524455 |
| O | -0.881262 | 0.192094 | -1.921129 |
| H | -0.367392 | 0.187568 | -2.741216 |
| H | -1.728087 | -0.307143 | -2.089245 |

---------------------------------------
Statistical Thermodynamic Analysis
Temperature 298.150 Kelvin. Pressure 1.00000 Atm.
---------------------------------------
SCF = -1538.60186892 | Predicted change in Energy=-8.445831D-09
Zero-point correction (ZPE) = -1538.22537492 0.376494
Internal Energy (U) = -1538.19657392 0.405295
Enthalpy (H) = -1538.19562892 0.40624
Gibbs Free Energy (G) = -1538.29286092 0.309008
---------------------------------------
Frequencies
11.8420 15.1086 19.7912
25.7923 26.7505 32.4007
35.3989 40.1920 49.3759
60.6652 67.8416 85.2550
100.3093 106.7056 112.6788
120.9534 123.0089 128.0450
135.0776 148.0472 163.2587
182.0019 191.1079 199.0891
260.5346 265.0129 273.4863
274.6695 276.2403 309.5467
330.9985 337.3725 347.8627
349.5163 351.4176 628.9748
629.7760 631.2708 644.8554
660.7090 662.5218 664.3583
665.6466 666.1331 666.4625
681.0249 687.7121 687.9431
688.9425 760.8011 807.3024
809.5764 814.1729 816.9258
829.7825 832.7234 834.3004
977.1360 979.3099 979.8617
990.3363 991.8738 992.0799
992.1945 1032.9774 1033.2424

1034.3610 1066.8613 1066.9586
1067.0304 1101.7881 1132.3995
1135.1023 1140.3889 1166.7382
1168.1434 1169.8591 1256.3875
1257.5707 1260.6447 1294.6672
1295.4106 1296.0384 1366.9943
1369.2967 1369.9048 1377.9353
1432.2310 1432.2963 1432.5197
1446.3790 1447.0651 1447.6032
1475.7957 1483.0611 1483.2836
1483.3504 1486.9220 1495.1930
1495.2053 1495.5161 1543.0377
1544.2789 1549.4452 1644.2729
1644.6596 1645.6901 1649.3658
2696.4312 3054.2339 3054.2712
3054.5544 3085.0289 3115.0236
3115.1655 3115.6507 3153.3733
3153.6470 3153.6676 3183.8082
3191.1191 3227.0994 3284.6233
3285.4013 3286.9944 3289.3648
3292.5584 3294.8651 3627.5412
3628.9799 3629.5560 3838.9170

**$[Zn(Im)_3(CH_3SH)]^{2+}$ Complexed with HOCl**

---------------------------------------
Gaussian 16: ES64L-G16RevB.01 20-Dec-2017
---------------------------------------
### B3LYP/GEN PSEUDO=READ gfprint gfinput
scf=(direct,tight,maxcycle=300
,xqc) opt=(maxcycle=250) freq=noraman
AN.EDU\22-Jun-2020\0\\# B3LYP/GEN PSEUDO=READ
gfprint gfinput scf=(dir
ect,tight,maxcycle=300,xqc) opt=(maxcycle=250)
freq=noraman\\zn_3imid_
---------------------------------------
Full point group C1 NOp 1
Stoichiometry C13H23ClN6OSZn(2+) Framework group
C1[X(C13H23ClN6OSZn)]
---------------------------------------
Num atoms: 46
Charge = 2 Multiplicity = 1
---------------------------------------
SCF = -1998.11095325 | Predicted change in Energy=-
5.922149D-09

Optimization completed.
Maximum Force 0.000009 0.000450 YES
RMS Force 0.000001 0.000300 YES
Maximum Displacement 0.001425 0.001800 YES
RMS Displacement 0.000287 0.001200 YES

---------------------------------------
Atom Coordinates (in Angstroms)
Type X Y Z
---------------------------------------
Zn -0.357871 -0.043291 0.255741
O 4.004198 0.265561 -1.451134
N 0.132458 1.776483 -0.448245
C -0.736334 2.863989 -0.439255
C -0.111624 3.961498 -0.973940
C 1.274747 2.207154 -0.983208
N 0.265162 -1.675227 -0.739471
C -0.487922 -2.839348 -0.864332
C 0.203395 -3.761777 -1.607467
C 1.403809 -1.886736 -1.399004
S 0.460850 -0.144723 2.607300
H 2.227153 -1.193827 -1.507167
H -1.468301 -2.932975 -0.423035
H 2.172110 1.626088 -1.146673
H -1.742590 2.785027 -0.057742
C -0.568050 5.364068 -1.203950
H -0.533644 5.627271 -2.266654
H -1.598378 5.481761 -0.862834
H 0.051334 6.081956 -0.655496
C -0.127377 -5.150047 -2.045512
H -0.146685 -5.231168 -3.137695
H 0.599698 -5.874948 -1.663833
H -1.112455 -5.434898 -1.670949
H 2.142791 -3.530971 -2.478729
N 1.394092 -3.127869 -1.928251
H 1.885813 4.074057 -1.729235
N 1.156912 3.511693 -1.306823
C 1.665353 -1.523995 2.803425
H 1.089021 -2.448413 2.750835
H 2.419769 -1.503322 2.017892
H 2.123652 -1.439097 3.789014
H -2.676518 -0.140890 2.621042
C -4.568842 -0.221440 -0.042644
C -3.295722 -0.175858 -0.553069
N -2.371348 -0.138987 0.487151
C -3.075637 -0.161100 1.617709
N -4.393227 -0.210844 1.332787
H 4.722198 0.370365 -2.101967
H -5.136516 -0.236317 2.020975
H -2.991146 -0.166379 -1.588957
C -5.910540 -0.273143 -0.695047
H -6.457662 -1.179641 -0.414481
H -6.523499 0.591763 -0.419357
H -5.797686 -0.271779 -1.780919
Cl 4.832257 0.103079 0.058352
H 1.320801 0.893199 2.614851
---------------------------------------
Statistical Thermodynamic Analysis
Temperature 298.150 Kelvin. Pressure 1.00000 Atm.
---------------------------------------
SCF = -1998.11095325 | Predicted change in Energy=-
5.922149D-09
Zero-point correction (ZPE) = -1997.74404425 0.366909
Internal Energy (U) = -1997.71367125 0.397282
Enthalpy (H) = -1997.71272725 0.398226
Gibbs Free Energy (G) = -1997.81483825 0.296115
---------------------------------------
Frequencies
8.5147 14.4245 18.7683
25.0049 26.4512 33.3685
37.5084 40.3745 41.6368
52.7157 60.4154 66.9554
81.4647 98.7414 105.0412
105.8896 110.8134 118.8444
124.6763 127.6824 133.8278
140.7757 153.0103 161.0107
178.4909 195.6271 214.2588
256.2493 266.2161 274.3050
276.3007 279.1450 308.7947

348.1459 348.9904 353.1350
481.0823 629.6504 631.8075
634.2697 661.2603 663.0648
664.1469 664.9289 665.1919
665.9717 677.0709 686.6250
687.8219 688.5353 724.6011
803.5347 809.5262 818.8204
821.1151 833.2034 860.5780
867.5882 976.3822 978.9226
979.6175 991.6499 992.2353
992.2824 997.7914 1033.4336
1034.4635 1035.2882 1066.5606
1066.8620 1066.9873 1100.5580
1133.5244 1134.3386 1141.3938
1168.6510 1170.4156 1172.6417
1257.2616 1258.1847 1263.3276
1268.5076 1295.6355 1296.9606
1298.5104 1366.9001 1368.3919
1370.8055 1379.5393 1432.2072
1432.2998 1432.4946 1446.6822
1446.9164 1447.8966 1472.0742
1481.7460 1483.0481 1483.2784
1483.4703 1495.1555 1495.2145
1495.4807 1544.1783 1547.1393
1552.8363 1643.7586 1645.8870
1646.5524 2692.4935 3054.0800
3054.0950 3054.2280 3087.2814
3114.8656 3114.9266 3115.0022
3153.3327 3153.3853 3153.4710
3189.6521 3196.9800 3264.7648
3270.0311 3284.7217 3290.7136
3291.7111 3293.1827 3627.5707
3630.9503 3631.5435 3740.9848

**$[Zn(Im)_3(HOCl)]^{2+}$ Complexed with $CH_3SH$**

---------------------------------------
Gaussian 16: ES64L-G16RevB.01 20-Dec-2017
---------------------------------------
### B3LYP/GEN PSEUDO=READ gfprint gfinput
scf=(direct,tight,maxcycle=300
,xqc) opt=(maxcycle=250) freq=noraman
AN.EDU\23-Jun-2020\0\\# B3LYP/GEN PSEUDO=READ
gfprint gfinput scf=(dir
ect,tight,maxcycle=300,xqc) opt=(maxcycle=250)
freq=noraman\\zn_3imid_
---------------------------------------
Full point group C1 NOp 1
Stoichiometry C13H23ClN6OSZn(2+) Framework group
C1[X(C13H23ClN6OSZn)]
---------------------------------------
Num atoms: 46
Charge = 2 Multiplicity = 1
---------------------------------------
SCF = -1998.11489294 | Predicted change in Energy=-7.124956D-09

Optimization completed.
Maximum Force 0.000007 0.000450 YES
RMS Force 0.000001 0.000300 YES
Maximum Displacement 0.001291 0.001800 YES
RMS Displacement 0.000333 0.001200 YES

---------------------------------------

| Atom Type | Coordinates (in Angstroms) X | Y | Z |
|---|---|---|---|
| Zn | -0.193476 | -0.367488 | -0.059589 |
| N | 0.590893 | -2.207408 | 0.037452 |
| C | 0.776403 | -2.949960 | 1.200911 |
| C | 1.312032 | -4.173852 | 0.888532 |
| C | 1.008822 | -2.971745 | -0.970868 |
| N | -2.119648 | -0.246950 | -0.570551 |
| C | -2.739324 | 0.836360 | -1.185954 |
| C | -4.070260 | 0.562298 | -1.372900 |
| C | -3.062733 | -1.169573 | -0.385239 |
| N | 0.374187 | 0.852467 | 1.418699 |
| C | -0.446598 | 1.765387 | 2.074045 |
| C | 0.273252 | 2.427364 | 3.035830 |
| C | 1.580706 | 0.959098 | 1.974723 |
| H | 1.006592 | -2.704301 | -2.017143 |
| H | 0.514987 | -2.562709 | 2.174258 |
| H | 2.461905 | 0.395571 | 1.706347 |
| H | -1.488887 | 1.879638 | 1.818162 |
| H | -2.928196 | -2.140417 | 0.068177 |
| H | -2.191270 | 1.723762 | -1.463260 |
| C | -5.181505 | 1.351390 | -1.981709 |
| H | -5.607160 | 0.837665 | -2.850438 |
| H | -5.988389 | 1.528402 | -1.262448 |
| H | -4.810133 | 2.322232 | -2.315386 |
| C | 1.704540 | -5.342334 | 1.730576 |
| H | 1.139071 | -6.239857 | 1.457778 |
| H | 2.771260 | -5.569099 | 1.628419 |
| H | 1.506038 | -5.128111 | 2.782495 |
| C | -0.101771 | 3.484930 | 4.020303 |
| H | 0.490696 | 4.394872 | 3.876427 |
| H | -1.154689 | 3.747859 | 3.902018 |
| H | 0.045470 | 3.142638 | 5.050304 |
| H | 1.816788 | -4.908574 | -1.056907 |
| N | 1.446542 | -4.153463 | -0.491614 |
| H | 2.341679 | 2.162915 | 3.518341 |
| N | 1.550808 | 1.894274 | 2.944710 |
| H | -5.114154 | -1.226283 | -0.832663 |
| N | -4.240870 | -0.712988 | -0.854642 |
| H | 4.036285 | 1.627320 | -2.510084 |
| S | 3.204293 | 2.323278 | -1.709309 |
| C | 2.920542 | 3.771451 | -2.811618 |
| H | 2.220983 | 4.425874 | -2.289831 |
| H | 3.864089 | 4.297812 | -2.956800 |
| H | 2.498995 | 3.461847 | -3.767634 |
| O | 0.684919 | 0.600265 | -1.756079 |
| H | 1.533680 | 1.172230 | -1.794765 |
| Cl | 0.455606 | -0.035086 | -3.345077 |

---------------------------------------
Statistical Thermodynamic Analysis
Temperature 298.150 Kelvin. Pressure 1.00000 Atm.
---------------------------------------
SCF = -1998.11489294 | Predicted change in Energy=-7.124956D-09
Zero-point correction (ZPE) = -1997.74868894 0.366204
Internal Energy (U) = -1997.71872494 0.396168
Enthalpy (H) = -1997.71778094 0.397112
Gibbs Free Energy (G) = -1997.81948794 0.295405
---------------------------------------
Frequencies

11.1601 16.7521 18.3273
23.5658 24.8646 27.3568
30.6901 33.1894 39.6131
45.9478 55.1400 67.1214
77.5466 99.0135 101.8302
109.1836 112.7220 123.6068
128.3147 129.7576 133.9357
150.1022 156.0132 164.3213
180.2436 201.8574 268.2147
269.1836 273.4427 275.1794
278.0713 335.1499 348.5969
348.7631 351.7270 395.0910
629.0292 629.8227 632.1103
660.3043 661.0227 661.9572
665.8407 666.0277 666.9353
677.8444 686.4937 688.1950
689.4471 729.7122 792.8524
806.8295 808.5045 813.8069
827.6285 831.8934 834.2388
847.2734 977.6257 980.0126
980.3728 991.9070 992.1963
992.3851 995.0417 1032.9139
1033.7236 1033.9047 1066.8197
1066.8379 1066.9548 1100.6764
1132.5272 1135.3288 1140.6859
1168.9120 1168.9966 1169.1459
1257.7826 1258.3082 1258.8353
1295.5317 1295.6277 1295.7054
1367.4196 1367.8301 1369.8098
1378.1657 1415.6372 1432.1739
1432.3275 1432.5075 1447.0968
1447.1382 1447.5220 1475.5756
1482.9943 1483.0963 1483.1661
1485.1246 1495.2086 1495.3082
1495.5603 1544.2689 1544.9091
1549.3597 1644.8423 1645.3606
1646.1855 2689.1636 2781.3274
3054.3638 3054.5368 3054.5711
3085.8217 3115.3468 3115.5532
3115.6450 3153.6975 3153.7420
3153.9695 3185.3936 3192.4615
3285.6208 3289.7415 3290.0377
3292.6505 3293.0653 3293.3067
3627.2872 3627.9364 3628.8237

**$[Zn(Im)_3(CH_3SOH)]^{2+}$ Complexed with $H_2O$**

---------------------------------------
Gaussian 16: ES64L-G16RevB.01 20-Dec-2017
---------------------------------------
### B3LYP/GEN PSEUDO=READ gfprint gfinput scf=(direct,tight,maxcycle=300 ,xqc) opt=(maxcycle=250) freq=noraman
EDU\16-Jun-2020\0\\# B3LYP/GEN PSEUDO=READ gfprint gfinput scf=(direct ,tight,maxcycle=300,xqc) opt=(maxcycle=250) freq=noraman\\zn_3imid_soh
---------------------------------------
Full point group C1 NOp 1
Stoichiometry C13H24N6O2SZn(2+) Framework group C1[X(C13H24N6O2SZn)]
---------------------------------------
Num atoms: 47
Charge = 2 Multiplicity = 1
---------------------------------------
SCF = -1613.81099429 | Predicted change in Energy=-7.441317D-09

Optimization completed.

| | | | |
|---|---|---|---|
| Maximum Force | 0.000017 | 0.000450 | YES |
| RMS Force | 0.000002 | 0.000300 | YES |
| Maximum Displacement | 0.001758 | 0.001800 | YES |
| RMS Displacement | 0.000450 | 0.001200 | YES |

---------------------------------------

| Atom Type | Coordinates (in Angstroms) X | Y | Z |
|---|---|---|---|
| Zn | -0.181976 | -0.028055 | 0.364775 |
| N | -2.160499 | 0.320692 | 0.237835 |
| C | -2.955180 | -0.053714 | -0.841725 |
| C | -4.243816 | 0.370422 | -0.638499 |
| C | -2.956321 | 0.965322 | 1.089602 |
| N | 0.341593 | -1.820161 | -0.377942 |
| C | 1.128516 | -2.042699 | -1.502728 |
| C | 1.234724 | -3.390665 | -1.736034 |
| C | -0.028880 | -3.018444 | 0.068159 |
| N | 0.914835 | 1.477974 | -0.386544 |
| C | 0.410234 | 2.535225 | -1.137424 |
| C | 1.431708 | 3.375629 | -1.500546 |
| C | 2.229469 | 1.672495 | -0.294007 |
| O | 0.204804 | -0.173897 | 2.416314 |
| S | 1.836382 | -0.024643 | 2.988603 |
| H | -2.666225 | 1.381882 | 2.043994 |
| H | -2.555299 | -0.599447 | -1.683026 |
| H | 2.930172 | 1.041118 | 0.231618 |
| H | -0.641256 | 2.620548 | -1.365288 |
| H | -0.651970 | -3.215803 | 0.927771 |
| H | 1.559817 | -1.229199 | -2.065829 |
| C | 1.946104 | -4.174806 | -2.788503 |
| H | 2.697801 | -4.841552 | -2.352360 |
| H | 1.249234 | -4.784840 | -3.373300 |
| H | 2.456913 | -3.498108 | -3.476255 |
| C | -5.484611 | 0.240547 | -1.458323 |
| H | -6.262057 | -0.315282 | -0.923063 |
| H | -5.892250 | 1.221114 | -1.726883 |
| H | -5.265860 | -0.296009 | -2.383589 |
| C | 1.458247 | 4.639839 | -2.294047 |
| H | 2.085986 | 4.540043 | -3.186111 |
| H | 0.448821 | 4.895523 | -2.621979 |
| H | 1.841510 | 5.477172 | -1.700921 |
| H | -5.002546 | 1.444372 | 1.048628 |
| N | -4.209847 | 1.009831 | 0.591609 |
| H | 3.508166 | 3.170782 | -1.025254 |
| N | 2.567990 | 2.800895 | -0.950814 |
| H | 0.355458 | -4.975047 | -0.598055 |
| N | 0.491680 | -3.979048 | -0.723604 |
| C | 2.052609 | -1.625578 | 3.807399 |
| H | 3.079710 | -1.601353 | 4.190844 |
| H | 1.368402 | -1.739004 | 4.650264 |
| H | 1.958027 | -2.452318 | 3.101325 |
| H | -0.398050 | 0.256699 | 3.082590 |
| H | -1.046451 | 1.854839 | 4.592746 |
| O | -1.421756 | 1.082616 | 4.143330 |
| H | -1.899063 | 0.587091 | 4.825249 |

---------------------------------------
Statistical Thermodynamic Analysis
Temperature 298.150 Kelvin. Pressure 1.00000 Atm.
---------------------------------------
SCF = -1613.81099429 | Predicted change in Energy=-7.441317D-09
Zero-point correction (ZPE) = -1613.42833929 0.382655
Internal Energy (U) = -1613.39845029 0.412544
Enthalpy (H) = -1613.39750629 0.413488
Gibbs Free Energy (G) = -1613.49554629 0.315448
---------------------------------------
Frequencies
16.8535 19.1241 21.0255
29.2453 30.4231 31.3041
36.8851 48.0451 52.2183
66.6154 82.4483 91.2620
101.1990 101.6733 111.7172
113.7051 124.0381 128.0267
131.4969 155.1333 159.3444
160.3815 181.4708 188.4504
200.4960 243.6443 262.5877
264.8397 273.2094 274.2789
277.6175 288.0753 348.2626
349.2391 350.2144 369.1967
392.8524 453.9575 628.4281
629.6238 630.9936 661.2646
661.6532 662.6345 665.3912
666.0085 666.1852 683.4972
686.5385 686.9579 687.0110
711.4265 808.7633 812.6999
818.4374 829.6921 834.3109
847.9355 946.8735 976.7298
977.2926 978.9720 979.2977
989.3653 991.9611 992.1679
992.5527 1033.2393 1033.6859
1034.8900 1066.8113 1066.9284
1067.0363 1134.7636 1135.9877
1141.2379 1166.6131 1167.0284
1169.1063 1257.9631 1258.6185
1263.4799 1294.1492 1294.5031
1298.0421 1362.7068 1368.7914
1370.0772 1370.5961 1386.4043
1432.0969 1432.2205 1432.4047
1446.6663 1446.8703 1447.1908
1447.9576 1476.5339 1483.2473
1483.3338 1483.4728 1495.2477
1495.3432 1495.5726 1542.5202
1544.7287 1549.4818 1644.5559
1645.0730 1645.5838 1647.6522
3052.8675 3053.8750 3053.9883
3054.1033 3114.4571 3114.6463
3114.8271 3140.4596 3153.0427
3153.2346 3153.2378 3170.9962
3260.1721 3279.3564 3287.2294
3289.2710 3289.8069 3292.7143
3294.2914 3629.6931 3630.1825
3631.1896 3780.1093 3876.7063

**$[Zn(Im)_3(H_2O)]^{2+}$ Complexed with $CH_3SOH$**

---------------------------------------
Gaussian 16: ES64L-G16RevB.01 20-Dec-2017
---------------------------------------
### B3LYP/GEN PSEUDO=READ gfprint gfinput
scf=(direct,tight,maxcycle=300
,xqc) opt=(maxcycle=250) freq=noraman
EDU\16-Jun-2020\0\\# B3LYP/GEN PSEUDO=READ
gfprint gfinput scf=(direct
,tight,maxcycle=300,xqc) opt=(maxcycle=250)
freq=noraman\\zn_3_imid_h2
---------------------------------------
Full point group C1 NOp 1
Stoichiometry C13H24N6O2SZn(2+) Framework group
C1[X(C13H24N6O2SZn)]
---------------------------------------
Num atoms: 47
Charge = 2 Multiplicity = 1
---------------------------------------
SCF = -1613.81116677 | Predicted change in Energy=-2.323674D-09

Optimization completed.

| | | | |
|---|---|---|---|
| Maximum Force | 0.000021 | 0.000450 | YES |
| RMS Force | 0.000002 | 0.000300 | YES |
| Maximum Displacement | 0.001176 | 0.001800 | YES |
| RMS Displacement | 0.000296 | 0.001200 | YES |

---------------------------------------

| Atom Type | Coordinates (in Angstroms) X | Y | Z |
|---|---|---|---|
| Zn | -0.057314 | -0.290317 | 0.153210 |
| O | -1.951599 | 0.678705 | -3.565080 |
| N | -0.607973 | -1.574238 | 1.601930 |
| C | -0.065274 | -2.833634 | 1.833846 |
| C | -0.683106 | -3.414715 | 2.912260 |
| C | -1.545661 | -1.389880 | 2.530438 |
| N | 1.662492 | -0.757338 | -0.767102 |
| C | 2.913628 | -0.212454 | -0.500364 |
| C | 3.853485 | -0.785727 | -1.319090 |
| C | 1.837247 | -1.652058 | -1.737755 |
| S | -2.133906 | 2.392985 | -3.639319 |
| H | 1.067874 | -2.258909 | -2.191357 |
| H | 3.056076 | 0.550874 | 0.249369 |
| H | -2.164779 | -0.512829 | 2.648703 |
| H | 0.730578 | -3.232102 | 1.223512 |
| C | -0.496419 | -4.733364 | 3.586506 |
| H | -1.413561 | -5.331500 | 3.557599 |
| H | 0.289802 | -5.300683 | 3.084714 |
| H | -0.204272 | -4.609117 | 4.634819 |
| C | 5.325301 | -0.578197 | -1.456855 |
| H | 5.880924 | -1.499065 | -1.249329 |
| H | 5.589819 | -0.241269 | -2.464965 |
| H | 5.662118 | 0.182576 | -0.750035 |
| H | 3.531710 | -2.290517 | -2.805604 |
| N | 3.139333 | -1.691755 | -2.088833 |
| H | -2.247959 | -2.583197 | 4.111636 |
| N | -1.614610 | -2.475164 | 3.328207 |
| C | -3.911603 | 2.533148 | -3.969800 |
| H | -4.106434 | 3.611636 | -4.002231 |
| H | -4.166176 | 2.104992 | -4.941256 |
| H | -4.505217 | 2.084969 | -3.171053 |
| H | -1.174524 | 2.623704 | -0.888729 |
| C | 0.077212 | 3.477983 | 2.011715 |
| C | 0.294282 | 2.124685 | 1.954304 |

N  -0.196815   1.619957   0.753163
C  -0.706398   2.654166   0.086507
N  -0.557327   3.779294   0.816173
H  -1.744571   0.341381  -4.451559
H  -0.862636   4.701817   0.530337
H   0.765244   1.491544   2.691355
C   0.395806   4.502955   3.049281
H   1.064162   5.276409   2.655715
H  -0.510651   4.994017   3.419461
H   0.892994   4.031845   3.899438
O  -1.463154  -0.649689  -1.334808
H  -1.620822  -0.141853  -2.179564
H  -2.264328  -1.154067  -1.139507
---------------------------------------
Statistical Thermodynamic Analysis
Temperature 298.150 Kelvin. Pressure 1.00000 Atm.
---------------------------------------
SCF = -1613.81116677 |  Predicted change in Energy=-2.323674D-09
Zero-point correction (ZPE) = -1613.42865777   0.382509
Internal Energy (U) = -1613.39888377   0.412283
Enthalpy (H) = -1613.39793977   0.413227
Gibbs Free Energy (G) = -1613.49732077   0.313846
---------------------------------------
Frequencies
10.9816 17.7288 21.1963
25.2865 26.1262 32.6656
33.9491 37.6066 41.7708
51.7477 68.9563 81.7570
97.3410 100.2325 112.5402
114.9311 123.5085 129.7568
134.5693 151.9550 160.7724
181.0849 185.1295 194.6378
213.5117 259.2124 265.6793
272.9625 275.3751 276.7848
286.4246 296.3229 347.6503
349.0247 351.4672 377.4446
410.4652 628.2951 630.0611
631.4417 653.8156 661.3263
663.3104 665.8031 665.9837
666.5031 667.7901 680.7398
686.1308 686.6056 688.3149
716.0136 807.2246 809.5094
817.0946 828.4278 830.3508
857.8415 886.3398 977.3354
978.0790 979.4080 979.8252
988.3279 991.9280 992.1510
992.2854 1033.3669 1033.4417
1034.7739 1066.8744 1067.0336
1067.1382 1134.6090 1136.2038
1141.4336 1166.8127 1167.1323
1170.1682 1207.9556 1257.2299
1257.8306 1264.0539 1294.4546
1295.9169 1297.7331 1365.8323
1368.2122 1369.1545 1370.7453
1432.0801 1432.2072 1432.5179
1446.3189 1446.3424 1447.5444
1448.8950 1477.0819 1483.2619
1483.3430 1483.4455 1495.2450
1495.3094 1495.5236 1542.8955
1545.1576 1550.6758 1644.7269
1644.9603 1645.7743 1667.1554
3053.1995 3053.9014 3054.1639
3054.4313 3114.5643 3114.9487
3115.4099 3141.6092 3152.9056
3153.4620 3153.6728 3170.8808
3212.2695 3254.7787 3284.1289
3288.5138 3288.7830 3292.3095
3293.6168 3628.5411 3630.4025
3631.9704 3774.3226 3854.0272

##### $[Zn(Im)_3(CH_3SOH)]^{2+}$ Complexed with HOCl

---------------------------------------
Gaussian 16: ES64L-G16RevB.01 20-Dec-2017
---------------------------------------
### B3LYP/GEN PSEUDO=READ gfprint gfinput scf=(direct,tight,maxcycle=300 ,xqc) opt=(maxcycle=250) freq=noraman
AN.EDU\21-Jun-2020\0\\# B3LYP/GEN PSEUDO=READ gfprint gfinput scf=(dir ect,tight,maxcycle=300,xqc) opt=(maxcycle=250) freq=noraman\\zn_3imid_
---------------------------------------
Full point group C1 NOp 1
Stoichiometry C13H23ClN6O2SZn(2+) Framework group C1[X(C13H23ClN6O2SZn)]
---------------------------------------
Num atoms: 47
Charge = 2 Multiplicity = 1
---------------------------------------
SCF = -2073.32248191 |  Predicted change in Energy=-3.887317D-09

Optimization completed.
Maximum Force   0.000025   0.000450   YES
RMS   Force   0.000002   0.000300   YES
Maximum Displacement   0.001419   0.001800   YES
RMS   Displacement   0.000346   0.001200   YES

---------------------------------------
Atom   Coordinates (in Angstroms)
Type   X   Y   Z
---------------------------------------
Zn  -0.012940   0.008259   0.101918
N  -1.811104  -0.892422   0.112684
C  -2.194024  -1.894785  -0.773833
C  -3.490701  -2.265677  -0.522939
C  -2.863480  -0.656229   0.894683
N   1.456935  -1.178304  -0.574671
C   2.160539  -1.014637  -1.763267
C   3.059632  -2.039947  -1.915140
C   1.920790  -2.291112  -0.009464
N  -0.042630   1.818497  -0.759289
C  -1.080945   2.311662  -1.544043
C  -0.767642   3.570393  -1.989860
C   0.892851   2.766759  -0.728767
O   0.440807   0.243558   2.143867
S   1.613982   1.411291   2.674346
H  -2.912004   0.055921   1.705447
H  -1.521040  -2.278790  -1.525416
H   1.832762   2.716299  -0.199745
H  -1.972882   1.735140  -1.736170
H   1.591669  -2.717363   0.926580
H   1.975807  -0.181806  -2.424558

C 4.051898 -2.355187 -2.984991
H 5.074078 -2.370628 -2.591648
H 3.849879 -3.328750 -3.444448
H 4.006287 -1.598405 -3.770505
C -4.387724 -3.274295 -1.160500
H -4.707872 -4.036632 -0.442052
H -5.284173 -2.804793 -1.579670
H -3.862751 -3.779183 -1.973711
C -1.504805 4.544240 -2.848171
H -0.941187 4.784933 -3.755998
H -2.464455 4.121701 -3.152089
H -1.703067 5.479372 -2.313349
H -4.793535 -1.481828 0.982493
N -3.883244 -1.463900 0.538236
H 1.016755 4.680553 -1.586129
N 0.486378 3.827266 -1.455765
H 3.392127 -3.679429 -0.580941
N 2.882387 -2.828747 -0.788193
C 2.801259 0.333666 3.516751
H 3.576224 1.016902 3.884850
H 2.346629 -0.170841 4.371284
H 3.256634 -0.378220 2.826286
H -0.277056 0.205573 2.821296
H -1.657706 0.951712 4.652301
O -1.622911 0.265314 3.960782
Cl -1.913431 -1.219098 4.801514
---------------------------------------
Statistical Thermodynamic Analysis
Temperature 298.150 Kelvin. Pressure 1.00000 Atm.
---------------------------------------
SCF = -2073.32248191 | Predicted change in Energy=-3.887317D-09
Zero-point correction (ZPE) = -2072.94968191 0.3728
Internal Energy (U) = -2072.91856191 0.40392
Enthalpy (H) = -2072.91761791 0.404864
Gibbs Free Energy (G) = -2073.02163791 0.300844
---------------------------------------
Frequencies
9.5750 14.2081 16.1424
23.1110 26.6472 28.2845
33.8188 38.3722 44.6121
48.0510 51.0694 69.0641
83.2357 91.0584 99.1551
102.0310 110.1906 115.1359
124.2828 127.9195 130.8222
153.0138 158.1997 159.8845
179.3890 188.1748 198.8211
263.4873 266.3303 273.2120
274.1046 274.7795 285.9080
348.3212 349.0496 350.0580
353.7645 374.4565 628.8375
628.9757 630.3946 661.0093
661.3351 661.8843 665.3287
666.0537 666.1798 681.0805
686.3615 687.4336 687.8834
701.8828 728.9340 806.8684
808.7330 810.5697 815.0749
829.7700 831.6690 833.1342
976.9151 977.1644 979.2632
979.4917 988.4877 991.9993
992.2199 992.3823 1033.1524
1033.6974 1034.3252 1066.9392
1066.9701 1067.0380 1134.2143
1134.9358 1140.3031 1166.8892
1167.4960 1169.0588 1246.4711
1257.8146 1258.3425 1261.1124
1294.4100 1294.6030 1296.2271
1353.0007 1364.3164 1368.5085
1369.4327 1369.9581 1432.0817
1432.3072 1432.4493 1446.5486
1447.0669 1447.2529 1447.5193
1476.3342 1483.2078 1483.2898
1483.4156 1495.2890 1495.3622
1495.5677 1542.8468 1544.6265
1549.2516 1644.5679 1645.1850
1645.6731 3052.3455 3054.1219
3054.1440 3054.2328 3114.9102
3114.9327 3115.0626 3140.0794
3153.2344 3153.4149 3153.4197
3172.1566 3286.7771 3288.8718
3289.8352 3289.9700 3292.4878
3294.3046 3424.7502 3629.1022
3629.5844 3630.1810 3738.0442

**$[Zn(Im)_3(HOCl)]^{2+}$ Complexed with $CH_3SOH$**

---------------------------------------
Gaussian 16: ES64L-G16RevB.01 20-Dec-2017
---------------------------------------
### B3LYP/GEN PSEUDO=READ gfprint gfinput scf=(direct,tight,maxcycle=300
,xqc) opt=(maxcycle=250) freq=noraman
AN.EDU\23-Jun-2020\0\\# B3LYP/GEN PSEUDO=READ gfprint gfinput scf=(dir
ect,tight,maxcycle=300,xqc) opt=(maxcycle=250) freq=noraman\\zn_3im_ho
---------------------------------------
Full point group C1 NOp 1
Stoichiometry C13H23ClN6O2SZn(2+) Framework group C1[X(C13H23ClN6O2SZn)]
---------------------------------------
Num atoms: 47
Charge = 2 Multiplicity = 1
---------------------------------------
SCF = -2073.32343335 | Predicted change in Energy=-1.097327D-08

Optimization completed.
Maximum Force 0.000016 0.000450 YES
RMS Force 0.000003 0.000300 YES
Maximum Displacement 0.001788 0.001800 YES
RMS Displacement 0.000453 0.001200 YES

---------------------------------------
Atom Coordinates (in Angstroms)
Type X Y Z
---------------------------------------
Zn -0.103202 -0.544673 0.422033
N -0.302952 -1.976089 1.802140
C -1.413423 -2.190695 2.613013
C -1.184673 -3.264799 3.435018
C 0.590905 -2.909055 2.125921
N -1.747204 -0.167913 -0.651443
C -2.458236 1.029038 -0.645171
C -3.558746 0.918290 -1.456269

C -2.409094 -0.997599 -1.456668
N 0.893896 1.086945 1.003710
C 1.283738 1.365703 2.310123
C 1.928428 2.575558 2.352245
C 1.297602 2.115269 0.258033
H 1.566685 -3.037061 1.682026
H -2.293822 -1.568824 2.553191
H 1.170224 2.220896 -0.809647
H 1.073733 0.688387 3.123769
H -2.133197 -2.014542 -1.691511
H -2.140567 1.878354 -0.059695
C -4.647912 1.876803 -1.807484
H -4.674791 2.075758 -2.884272
H -5.629512 1.493963 -1.508121
H -4.488657 2.827125 -1.294293
C -2.019945 -3.922767 4.482684
H -2.217285 -4.971755 4.236781
H -1.532444 -3.889398 5.462984
H -2.980870 -3.411855 4.568789
C 2.537845 3.349675 3.473696
H 2.054485 4.324995 3.595659
H 2.424743 2.800467 4.410357
H 3.607608 3.518233 3.309531
H 0.574061 -4.481455 3.518448
N 0.091175 -3.694830 3.100699
H 2.320988 3.892844 0.712275
N 1.920940 3.021049 1.038409
H -4.163457 -0.791393 -2.592612
N -3.496183 -0.374713 -1.954094
O 1.346494 -1.499757 -0.831652
Cl 0.886846 -2.886451 -1.748825
H 2.119769 -1.073986 -1.344381
O 3.306381 -0.475006 -2.124300
S 3.264601 1.020958 -2.987600
C 3.193150 0.426574 -4.698337
H 2.303977 -0.181527 -4.873029
H 3.130409 1.338023 -5.304801
H 4.103841 -0.110929 -4.969225
H 4.225649 -0.754834 -1.985102
---------------------------------------
Statistical Thermodynamic Analysis
Temperature 298.150 Kelvin. Pressure 1.00000 Atm.
---------------------------------------
SCF = -2073.32343335 | Predicted change in Energy=-1.097327D-08
Zero-point correction (ZPE) = -2072.95118135 0.372252
Internal Energy (U) = -2072.92015135 0.403282
Enthalpy (H) = -2072.91920735 0.404226
Gibbs Free Energy (G) = -2073.02391535 0.299518
---------------------------------------
Frequencies
8.9248 10.8554 16.4545
22.4930 22.9893 26.5770
30.3459 36.8684 39.8392
45.3150 46.7303 53.4722
84.3926 94.6238 101.5593
105.7467 110.7192 120.8667
123.9137 128.5427 139.1283
156.6997 159.6331 162.2292
186.6729 193.8420 198.5558
268.3671 271.3289 274.1280
274.9957 276.3399 280.1327
348.5767 349.6490 350.6938
383.3606 389.2552 628.5479
629.4506 630.4422 659.8695
660.9555 661.2341 665.8720
666.2651 666.6736 680.6302
687.3878 687.4394 687.8119
712.5259 736.4792 807.1466
809.1634 813.3022 828.8558
830.7907 831.5595 957.3076
977.5888 979.0633 980.0764
980.2192 991.9080 992.0735
992.3328 992.8285 1033.3786
1033.7519 1034.0830 1066.8355
1067.0404 1067.1206 1133.8544
1135.1808 1140.3181 1168.4114
1168.5121 1169.1266 1219.1797
1257.5853 1258.8274 1259.9775
1295.1059 1295.7854 1296.0446
1365.3567 1368.0293 1368.5152
1369.3012 1432.1850 1432.4321
1432.5634 1444.5236 1446.6494
1447.5550 1448.0083 1448.5611
1476.4077 1483.1054 1483.1638
1483.3603 1495.0477 1495.3952
1495.5685 1543.9956 1544.6036
1549.2404 1645.4329 1645.6364
1646.0432 2842.7040 3053.0850
3054.2319 3054.3623 3054.3761
3115.1052 3115.3158 3115.3374
3141.8659 3153.3709 3153.5728
3153.9579 3172.9632 3284.4562
3287.4392 3288.9729 3291.4062
3296.6410 3296.8772 3628.6340
3629.1670 3629.6429 3774.8641

**$[Zn(Im)_3(CH_3SO_2H)]^{2+}$ Complexed with $H_2O$**

---------------------------------------
Gaussian 16: ES64L-G16RevB.01 20-Dec-2017
---------------------------------------
### B3LYP/GEN PSEUDO=READ gfprint gfinput scf=(direct,tight,maxcycle=300
,xqc) OPT freq=noraman
17-Jul-2020\0\\# B3LYP/GEN PSEUDO=READ gfprint gfinput scf=(direct,tig
ht,maxcycle=300,xqc) OPT freq=noraman\\zn_3im_1meso2h_ctr_h2o.xyz\\2,1
---------------------------------------
Full point group C1 NOp 1
Stoichiometry C13H24N6O3SZn(2+) Framework group C1[X(C13H24N6O3SZn)]
---------------------------------------
Num atoms: 48
Charge = 2 Multiplicity = 1
---------------------------------------
SCF = -1689.01310743 | Predicted change in Energy=-1.280941D-08

Optimization completed.
Maximum Force 0.000020 0.000450 YES
RMS Force 0.000004 0.000300 YES
Maximum Displacement 0.001641 0.001800 YES
RMS Displacement 0.000438 0.001200 YES

---------------------------------------
Atom Coordinates (in Angstroms)
Type X Y Z
---------------------------------------
Zn -0.086765 0.107393 0.205993
N -1.922407 -0.236134 -0.520508
C -2.219917 -1.257216 -1.419032
C -3.550883 -1.212962 -1.746079
C -3.060599 0.422910 -0.301887
N 0.949475 -1.547685 0.665126
C 1.876515 -2.190634 -0.149883
C 2.327241 -3.329191 0.469281
C 0.836226 -2.287341 1.766247
N 1.010659 1.406279 -0.859188
C 2.392655 1.568423 -0.815965
C 2.770542 2.558745 -1.686495
C 0.554585 2.288396 -1.747004
H -3.191049 1.266186 0.362861
H -1.469335 -1.951925 -1.764406
H -0.477171 2.435239 -2.029028
H 3.017452 0.971725 -0.168880
H 0.197149 -2.079852 2.611447
H 2.157335 -1.798593 -1.115838
C 3.309235 -4.374865 0.056222
H 2.842341 -5.364391 0.005405
H 3.708772 -4.141777 -0.932663
H 4.151189 -4.432044 0.754533
C -4.391816 -2.055413 -2.646874
H -4.837487 -1.460450 -3.451348
H -3.781229 -2.835374 -3.105843
H -5.202926 -2.543835 -2.096097
C 4.102452 3.134769 -2.036198
H 4.330482 2.999733 -3.099000
H 4.145745 4.206224 -1.812876
H 4.886360 2.639988 -1.459524
H -5.015221 0.171800 -1.029303
N -4.051573 -0.140510 -1.023115
H 1.498960 3.726111 -2.951722
N 1.583533 2.991980 -2.258585
H 1.747390 -4.075971 2.388592
N 1.649808 -3.359577 1.678832
H -3.421724 2.435963 2.990300
O -2.692920 2.352746 2.358766
H -2.264475 3.226971 2.285366
O -0.375189 1.043921 2.096928
H -1.318839 1.305487 2.307634
C 0.445935 2.360115 4.268635
H 1.001083 3.211786 4.672868
H -0.606648 2.432233 4.551094
H 0.891305 1.422764 4.609404
S 0.606227 2.466281 2.461416
O -0.202374 3.646675 2.027990
---------------------------------------
Statistical Thermodynamic Analysis
Temperature 298.150 Kelvin. Pressure 1.00000 Atm.
---------------------------------------
SCF = -1689.01310743 | Predicted change in Energy=-1.280941D-08
Zero-point correction (ZPE) = -1688.62645343 0.386654
Internal Energy (U) = -1688.59614643 0.416961
Enthalpy (H) = -1688.59520243 0.417905
Gibbs Free Energy (G) = -1688.69387543 0.319232
---------------------------------------
Frequencies
12.1353 18.6295 24.5101
27.3757 29.6155 32.7594
35.2204 43.6346 52.2493
69.6129 94.0046 100.1080
101.2193 110.2014 114.3775
123.1626 127.9762 130.4864
144.7182 149.8454 159.4552
165.6159 185.3944 187.8101
199.6598 242.9952 263.8595
267.8079 273.6430 274.8551
278.9451 340.3174 346.2384
348.3431 350.1048 351.6276
389.9899 414.0692 448.1399
580.8365 600.1218 627.6094
629.2990 633.7039 660.6367
661.0052 663.3687 665.7567
666.1153 666.7884 668.8299
686.0453 687.0345 687.6126
805.1994 811.2414 815.3739
826.3193 832.2041 864.8399
898.7290 954.4925 970.1107
977.7685 979.5755 980.0394
991.6938 992.1639 993.0672
1032.7291 1033.8786 1035.8695
1066.5473 1066.6605 1067.0847
1127.2332 1133.4380 1135.0787
1141.7487 1167.8690 1169.2581
1171.1728 1257.9394 1258.4964
1265.5426 1295.0698 1295.3231
1300.7034 1341.9048 1356.5683
1367.2452 1369.7064 1370.5383
1432.1610 1432.4140 1432.5899
1442.3982 1446.9871 1447.1303
1448.4999 1452.5763 1483.1120
1483.2858 1483.3686 1495.2632
1495.3818 1495.6451 1544.2916
1546.5005 1550.6643 1620.1072
1644.4621 1645.9981 1646.6963
3054.1509 3054.2614 3054.3027
3061.4172 3114.9113 3115.2362
3115.2707 3153.1499 3153.4295
3153.5289 3164.4038 3180.6448
3203.0837 3267.8759 3285.8025
3287.8507 3288.1242 3294.1367
3294.7245 3628.7403 3629.0779
3630.5405 3674.7781 3856.4924

**$[Zn(Im)_3(H_2O)]^{2+}$ Complexed with $CH_3SO_2H$**
---------------------------------------
Gaussian 16: ES64L-G16RevB.01 20-Dec-2017
---------------------------------------
### B3LYP/GEN PSEUDO=READ gfprint gfinput scf=(direct,tight,maxcycle=300 ,xqc) OPT freq=noraman
17-Jul-2020\0\\# B3LYP/GEN PSEUDO=READ gfprint gfinput scf=(direct,tig ht,maxcycle=300,xqc) OPT freq=noraman\\zn_3im_1h2o_ctr_meso2h.xyz\\2,1
---------------------------------------

Full point group C1 NOp 1
Stoichiometry C13H24N6O3SZn(2+) Framework group C1[X(C13H24N6O3SZn)]
---------------------------------------
Num atoms: 48
Charge = 2 Multiplicity = 1
---------------------------------------
SCF = -1689.02109991 | Predicted change in Energy=-2.354933D-08

Optimization completed.

| | | | |
|---|---|---|---|
| Maximum Force | 0.000022 | 0.000450 | YES |
| RMS Force | 0.000005 | 0.000300 | YES |
| Maximum Displacement | 0.001372 | 0.001800 | YES |
| RMS Displacement | 0.000327 | 0.001200 | YES |

---------------------------------------

| Atom Type | Coordinates (in Angstroms) X | Y | Z |
|---|---|---|---|
| Zn | -0.276961 | -0.033805 | -0.244728 |
| N | 0.647553 | -1.807497 | -0.101281 |
| C | 1.013738 | -2.436045 | 1.083629 |
| C | 1.644444 | -3.621776 | 0.803765 |
| C | 1.051117 | -2.599527 | -1.092026 |
| N | -2.265827 | -0.137554 | -0.591657 |
| C | -3.140394 | 0.938675 | -0.710095 |
| C | -4.414800 | 0.480115 | -0.931911 |
| C | -3.000676 | -1.237916 | -0.737454 |
| N | 0.107792 | 1.228723 | 1.250678 |
| C | -0.764777 | 1.531925 | 2.293389 |
| C | -0.146321 | 2.372971 | 3.181716 |
| C | 1.245536 | 1.876217 | 1.500868 |
| H | 0.930593 | -2.410107 | -2.148099 |
| H | 0.808062 | -1.997807 | 2.048416 |
| H | 2.130769 | 1.861829 | 0.878321 |
| H | -1.766172 | 1.131271 | 2.335581 |
| H | -2.641395 | -2.255399 | -0.695698 |
| H | -2.803610 | 1.960768 | -0.620873 |
| C | -5.715502 | 1.185152 | -1.130835 |
| H | -6.151353 | 0.953161 | -2.108592 |
| H | -6.443154 | 0.907871 | -0.360489 |
| H | -5.566401 | 2.265362 | -1.077986 |
| C | 2.235227 | -4.675714 | 1.680433 |
| H | 1.750301 | -5.645490 | 1.524935 |
| H | 3.307505 | -4.797748 | 1.492898 |
| H | 2.107762 | -4.401906 | 2.729535 |
| C | -0.605965 | 3.002971 | 4.454728 |
| H | -0.586068 | 4.096382 | 4.391232 |
| H | -1.631567 | 2.698783 | 4.672872 |
| H | 0.020796 | 2.700881 | 5.300792 |
| H | 2.046818 | -4.451120 | -1.126907 |
| N | 1.651150 | -3.694985 | -0.581308 |
| H | 1.846029 | 3.144703 | 3.061452 |
| N | 1.120210 | 2.570539 | 2.650402 |
| H | -5.049972 | -1.557798 | -1.079385 |
| N | -4.290770 | -0.900916 | -0.943097 |
| H | 3.060840 | 1.021000 | -3.997993 |
| O | 0.330111 | 0.638171 | -2.138588 |
| H | -0.328798 | 1.012304 | -2.739864 |
| H | 1.210052 | 0.981743 | -2.417901 |
| O | 2.725534 | 1.575979 | -3.271944 |
| C | 3.994409 | 3.621803 | -2.239047 |
| H | 4.696088 | 3.992620 | -1.487058 |
| H | 4.332698 | 3.901318 | -3.239721 |
| H | 2.984866 | 3.991342 | -2.045933 |
| S | 3.988661 | 1.813054 | -2.110579 |
| O | 3.326324 | 1.482574 | -0.807095 |

---------------------------------------
Statistical Thermodynamic Analysis
Temperature 298.150 Kelvin. Pressure 1.00000 Atm.
---------------------------------------
SCF = -1689.02109991 | Predicted change in Energy=-2.354933D-08
Zero-point correction (ZPE) = -1688.63487791 0.386222
Internal Energy (U) = -1688.60413291 0.416967
Enthalpy (H) = -1688.60318891 0.417911
Gibbs Free Energy (G) = -1688.70421991 0.31688
---------------------------------------
Frequencies
12.1309 17.0962 21.7292
23.4735 26.8692 32.2802
33.3336 38.3494 41.3600
54.7899 69.0587 80.4836
93.1802 101.7843 105.8490
112.4741 124.1396 127.0248
132.6536 149.2486 163.2110
179.3636 181.9947 194.7733
201.9662 256.2043 267.1987
271.9640 275.4944 276.2349
277.4367 299.3201 320.8697
339.9456 348.4104 349.1416
351.4524 370.0346 424.1875
599.7391 619.7174 628.5490
630.0993 631.4994 660.4421
661.9659 663.4839 665.5270
666.4212 667.0276 676.9650
684.9123 687.2120 687.3821
736.4213 808.2429 810.3025
817.5376 830.0758 831.1261
893.0944 956.0321 969.7558
976.6421 978.8556 979.8960
991.8548 992.2532 992.7787
1033.0348 1033.3658 1035.8262
1066.8639 1067.0720 1067.1291
1107.9250 1132.5743 1136.3181
1141.2778 1149.4294 1165.9927
1166.9600 1170.8647 1256.4155
1257.5319 1267.9300 1294.1309
1295.4675 1300.1397 1353.3251
1367.9708 1370.8119 1371.4269
1432.0748 1432.2394 1432.7219
1441.8489 1446.3552 1446.8973
1449.3350 1453.9464 1483.3377
1483.5864 1483.7046 1495.2438
1495.4948 1495.7360 1544.0643
1544.1770 1549.9460 1643.8103
1645.4272 1647.5138 1658.5867
3053.0500 3053.8737 3054.1812
3062.3817 3113.1248 3114.4422
3115.0025 3151.9482 3153.2447
3153.3192 3169.3141 3178.4910
3260.9749 3283.7250 3288.6320
3288.7521 3292.2948 3294.5619

3489.7145 3629.1581 3631.6977
3634.7292 3758.9483 3848.3140

**[Zn(Im)₃(CH₃SO₂H)]²⁺ Complexed with HOCl**

---------------------------------------
Gaussian 16: ES64L-G16RevB.01 20-Dec-2017
---------------------------------------
### B3LYP/GEN PSEUDO=READ gfprint gfinput scf=(direct,tight,maxcycle=300 ,xqc) OPT freq=noraman
DU\17-Jul-2020\0\\# B3LYP/GEN PSEUDO=READ gfprint gfinput scf=(direct, tight,maxcycle=300,xqc) OPT freq=noraman\\zn_3im_1meso2h_ctr_hocl.xyz\
---------------------------------------
Full point group C1 NOp 1
Stoichiometry C13H23ClN6O3SZn(2+) Framework group C1[X(C13H23ClN6O3SZn)]
---------------------------------------
Num atoms: 48
Charge = 2 Multiplicity = 1
---------------------------------------
SCF = -2148.51893145 | Predicted change in Energy=-9.268154D-09

Optimization completed.

| | | | |
|---|---|---|---|
| Maximum Force | 0.000017 | 0.000450 | YES |
| RMS Force | 0.000002 | 0.000300 | YES |
| Maximum Displacement | 0.001508 | 0.001800 | YES |
| RMS Displacement | 0.000283 | 0.001200 | YES |

---------------------------------------

| Atom Type | Coordinates (in Angstroms) X | Y | Z |
|---|---|---|---|
| Zn | -0.458577 | -0.147351 | 0.279071 |
| O | 4.157488 | 0.245475 | -1.345023 |
| N | 0.194280 | 1.677401 | -0.185811 |
| C | -0.605754 | 2.807806 | -0.339325 |
| C | 0.175203 | 3.883943 | -0.673009 |
| C | 1.449089 | 2.061281 | -0.423879 |
| N | 0.380944 | -1.782179 | -0.479646 |
| C | -0.312726 | -2.952888 | -0.780110 |
| C | 0.541966 | -3.864096 | -1.342728 |
| C | 1.645052 | -1.978082 | -0.859002 |
| H | 2.463356 | -1.277349 | -0.769456 |
| H | -1.367121 | -3.058308 | -0.576275 |
| H | 2.332395 | 1.438497 | -0.395738 |
| H | -1.676028 | 2.771656 | -0.204176 |
| C | -0.150910 | 5.313733 | -0.952076 |
| H | 0.139640 | 5.599129 | -1.968917 |
| H | -1.225609 | 5.477944 | -0.852513 |
| H | 0.358164 | 5.985292 | -0.252317 |
| C | 0.338999 | -5.254792 | -1.845710 |
| H | 0.578832 | -5.333867 | -2.911579 |
| H | 0.963213 | -5.972697 | -1.302734 |
| H | -0.703563 | -5.550799 | -1.714249 |
| H | 2.631605 | -3.609686 | -1.732093 |
| N | 1.768500 | -3.215390 | -1.377705 |
| H | 2.299695 | 3.909970 | -0.933698 |
| N | 1.465986 | 3.376636 | -0.717794 |
| H | -2.808376 | -0.847937 | 2.435070 |
| C | -4.637980 | -0.277549 | -0.214748 |
| C | -3.355145 | -0.073744 | -0.657304 |
| N | -2.459092 | -0.284241 | 0.386943 |
| C | -3.188201 | -0.613476 | 1.451947 |
| N | -4.496077 | -0.616861 | 1.122363 |
| H | 4.588834 | 0.363874 | -2.209724 |
| H | -5.255625 | -0.834764 | 1.756575 |
| H | -3.025581 | 0.201410 | -1.648117 |
| C | -5.961179 | -0.190930 | -0.900209 |
| H | -6.487501 | -1.151241 | -0.876989 |
| H | -6.605903 | 0.561027 | -0.432587 |
| H | -5.821854 | 0.089774 | -1.945962 |
| Cl | 5.483246 | 0.180226 | -0.234026 |
| O | 0.011148 | -0.211555 | 2.434841 |
| H | 0.022877 | 0.698297 | 2.787276 |
| C | 1.806788 | -0.292581 | 4.423711 |
| H | 2.736093 | -0.721173 | 4.810726 |
| H | 1.916599 | 0.792816 | 4.341323 |
| H | 0.969006 | -0.580023 | 5.063621 |
| S | 1.574908 | -1.007329 | 2.767275 |
| O | 2.563698 | -0.349702 | 1.869587 |

---------------------------------------
Statistical Thermodynamic Analysis
Temperature 298.150 Kelvin. Pressure 1.00000 Atm.
---------------------------------------
SCF = -2148.51893145 | Predicted change in Energy=-9.268154D-09
Zero-point correction (ZPE) = -2148.14322645 0.375705
Internal Energy (U) = -2148.11077645 0.408155
Enthalpy (H) = -2148.10983145 0.4091
Gibbs Free Energy (G) = -2148.21662445 0.302307
---------------------------------------
Frequencies
8.3253 14.9292 21.7350
23.0681 27.9887 30.7705
31.8960 36.9162 46.2852
47.0636 55.0074 59.4424
67.1225 71.0858 94.1534
95.4706 104.8693 112.4966
114.1739 124.3877 130.0963
132.0552 148.4254 155.7270
163.6017 186.2526 197.8081
198.7702 268.8846 271.0220
274.7259 276.8287 278.6295
310.8950 318.7969 347.3373
349.2377 352.6345 363.5974
395.1764 515.1734 563.1788
628.8584 631.6209 632.7902
659.8674 661.9522 662.9057
663.9064 665.9510 666.0576
667.1446 688.2893 689.9278
691.2651 724.6439 808.8494
815.4453 818.6386 832.9756
851.6929 860.2465 943.0977
967.1451 977.7814 981.1050
981.8363 991.5599 991.7863
992.2447 1033.3015 1034.3549
1035.1437 1066.8397 1067.1611
1067.2493 1125.2666 1132.7962
1134.5840 1141.4796 1164.3608
1167.7118 1171.6955 1175.4685
1252.7039 1256.1629 1262.2125

1267.1582 1296.1238 1296.6821
1298.2280 1351.0904 1364.8601
1365.4971 1369.4981 1432.2456
1432.8079 1433.0263 1438.9713
1446.3599 1449.4457 1451.1229
1455.4822 1483.0749 1483.3588
1483.4120 1495.0532 1495.2169
1495.5415 1543.7201 1545.7310
1552.2971 1644.3407 1647.3155
1649.1042 3053.7362 3053.7659
3053.8988 3054.4225 3114.3481
3114.5629 3115.3754 3152.9494
3152.9884 3153.9253 3156.2151
3173.4932 3276.3580 3282.2214
3284.4593 3289.4099 3292.7114
3295.0273 3628.2999 3631.1606
3631.9391 3710.2803 3752.9997

**$[Zn(Im)_3(HOCl)]^{2+}$ Complexed with $CH_3SO_2H$**

---------------------------------------
Gaussian 16: ES64L-G16RevB.01 20-Dec-2017
---------------------------------------
### B3LYP/GEN PSEUDO=READ gfprint gfinput scf=(direct,tight,maxcycle=300 ,xqc) OPT freq=noraman
DU\17-Jul-2020\0\\# B3LYP/GEN PSEUDO=READ gfprint gfinput scf=(direct, tight,maxcycle=300,xqc) OPT freq=noraman\\zn_3im_1hocl_ctr_meso2h.xyz\
---------------------------------------
Full point group C1 NOp 1
Stoichiometry C13H23ClN6O3SZn(2+) Framework group C1[X(C13H23ClN6O3SZn)]
---------------------------------------
Num atoms: 48
Charge = 2 Multiplicity = 1
---------------------------------------
SCF = -2148.53158744 | Predicted change in Energy=-2.221120D-08

Optimization completed.

| | | | |
|---|---|---|---|
| Maximum Force | 0.000028 | 0.000450 | YES |
| RMS Force | 0.000004 | 0.000300 | YES |
| Maximum Displacement | 0.001255 | 0.001800 | YES |
| RMS Displacement | 0.000305 | 0.001200 | YES |

---------------------------------------
Atom Coordinates (in Angstroms)

| Type | X | Y | Z |
|---|---|---|---|
| Zn | -0.373434 | -0.076873 | 0.112947 |
| N | -2.233356 | -0.862373 | 0.000102 |
| C | -3.284861 | -0.633919 | 0.884514 |
| C | -4.391521 | -1.339605 | 0.483077 |
| C | -2.695770 | -1.698526 | -0.926857 |
| N | -0.314956 | 1.905674 | -0.010498 |
| C | 0.820463 | 2.706503 | -0.098066 |
| C | 0.447397 | 4.025930 | -0.126828 |
| C | -1.363761 | 2.727732 | 0.011817 |
| N | 0.689193 | -1.054480 | 1.485618 |
| C | 1.986691 | -0.805625 | 1.922591 |
| C | 2.277784 | -1.625163 | 2.983157 |
| C | 0.198363 | -2.014324 | 2.270073 |
| H | -2.141396 | -2.086561 | -1.768093 |
| H | -3.180314 | 0.008870 | 1.746183 |
| H | -0.788383 | -2.447709 | 2.206837 |
| H | 2.618750 | -0.083975 | 1.428237 |
| H | -2.402708 | 2.439051 | 0.065932 |
| H | 1.810608 | 2.279288 | -0.159836 |
| C | 1.238476 | 5.288928 | -0.213270 |
| H | 0.977120 | 5.866539 | -1.106562 |
| H | 1.074198 | 5.925405 | 0.662997 |
| H | 2.303986 | 5.056611 | -0.262868 |
| C | -5.767576 | -1.457376 | 1.049843 |
| H | -6.523879 | -1.104929 | 0.340180 |
| H | -6.005748 | -2.493963 | 1.311591 |
| H | -5.851454 | -0.854801 | 1.956375 |
| C | 3.498679 | -1.775959 | 3.829139 |
| H | 3.907745 | -2.790455 | 3.769968 |
| H | 4.270728 | -1.080164 | 3.494932 |
| H | 3.285621 | -1.557453 | 4.881184 |
| H | -4.561898 | -2.616907 | -1.225289 |
| N | -3.983922 | -2.001608 | -0.664915 |
| H | 1.000510 | -3.086793 | 3.887169 |
| N | 1.127035 | -2.376581 | 3.176045 |
| H | -1.539740 | 4.819095 | -0.054569 |
| N | -0.938081 | 4.004220 | -0.055049 |
| H | 2.915023 | -0.721099 | -3.700528 |
| O | 0.362924 | -0.744440 | -1.820062 |
| H | 1.332133 | -0.835735 | -2.054099 |
| Cl | -0.327985 | 0.020670 | -3.207826 |
| O | 2.907308 | -0.909889 | -2.744597 |
| H | 5.893657 | -0.728635 | -0.744567 |
| S | 4.035199 | 0.140862 | -1.937192 |
| O | 3.272150 | 0.598307 | -0.730810 |
| C | 5.127762 | -1.191092 | -1.373258 |
| H | 4.536298 | -1.911839 | -0.804739 |
| H | 5.589441 | -1.658632 | -2.246432 |

---------------------------------------
Statistical Thermodynamic Analysis
Temperature 298.150 Kelvin. Pressure 1.00000 Atm.
---------------------------------------
SCF = -2148.53158744 | Predicted change in Energy=-2.221120D-08
Zero-point correction (ZPE) = -2148.15482344 0.376764
Internal Energy (U) = -2148.12322044 0.408367
Enthalpy (H) = -2148.12227644 0.409311
Gibbs Free Energy (G) = -2148.22515844 0.306429
---------------------------------------
Frequencies
15.2713 18.1914 22.9387
26.4122 29.9279 32.7517
35.6414 38.8409 44.1268
52.0828 64.2785 77.3777
86.5781 99.9373 104.4727
105.5330 114.5019 117.3779
125.6655 128.4508 135.3522
152.3429 159.8057 170.9260
180.0649 194.3529 205.7356
256.1460 270.1692 273.5933
279.0141 287.3078 291.6761
317.8725 333.4645 347.6807
349.7088 355.3998 377.3602
465.9737 611.3896 626.5920

628.0027 631.4225 660.3970
663.1324 664.1779 665.5482
667.9847 668.6523 678.1983
683.9974 685.0001 688.7190
727.3837 786.2522 806.1561
810.9408 816.2147 835.2931
845.8845 869.7419 957.4757
970.5248 976.9533 979.0042
979.6083 991.0807 991.8154
991.9488 1032.8770 1033.5768
1035.4717 1066.6704 1067.1806
1067.7098 1100.9265 1133.0508
1141.4818 1146.2431 1148.8747
1168.4726 1171.8140 1173.7415
1255.8649 1256.2424 1258.9085
1294.7744 1305.0166 1310.0097
1352.6560 1367.0041 1367.7785
1370.1454 1431.9707 1432.1860
1432.6520 1435.2001 1441.1249
1447.5567 1447.8512 1448.4176
1452.7529 1483.1769 1484.0010
1484.4418 1495.3600 1496.1162
1496.5768 1543.8590 1544.8319
1549.7180 1643.5371 1646.4411
1647.9806 3052.9999 3053.2742
3054.2392 3061.9627 3113.0290
3113.4569 3115.0784 3152.3154
3153.0333 3153.3037 3167.9888
3180.1710 3215.0508 3279.9350
3281.2021 3292.8503 3293.0529
3296.4125 3297.0653 3628.0323
3631.0613 3631.7963 3746.7665

**Single Point Energies in M06/SDD(Zn)/6-31+G(d,p)**

| Structure | E (Hartrees) |
|---|---|
| $[Zn(Im)_3(CH_3S)]^+$ Complexed with $H_2O$ | -1537.6651 |
| $[Zn(Im)_3(OH)]^+$ Complexed with $CH_3SH$ | -1537.6309 |
| $[Zn(Im)_3(CH_3S)]^+$ Complexed with HOCl | -1997.1519 |
| $[Zn(Im)_3(OCl)]^+$ Complexed with $CH_3SH$ | -1997.1319 |
| $[Zn(Im)_3(CH_3SO)]^+$ Complexed with $H_2O$ | -1612.8203 |
| $[Zn(Im)_3(OH)]^+$ Complexed with $CH_3SOH$ | -1612.8143 |
| $[Zn(Im)_3(CH_3SO)]^+$ Complexed with HOCl | -2072.3114 |
| $[Zn(Im)_3(OCl)]^+$ Complexed with $CH_3SOH$ | -2072.3168 |
| $[Zn(Im)_3(CH_3SO_2)]^+$ Complexed with $H_2O$ | -1688.0236 |
| $[Zn(Im)_3(OH)]^+$ Complexed with $CH_3SO_2H$ | -1688.0065 |
| $[Zn(Im)_3(CH_3SO_2)]^+$ Complexed with HOCl | -2147.5102 |
| $[Zn(Im)_3(OCl)]^+$ Complexed with $CH_3SO_2H$ | -2147.4917 |
| $[Zn(Im)_3(CH_3SH)]^{2+}$ Complexed with $H_2O$ | -1537.9511 |
| $[Zn(Im)_3(H_2O)]^{2+}$ Complexed with $CH_3SH$ | -1537.9465 |
| $[Zn(Im)_3(CH_3SH)]^{2+}$ Complexed with HOCl | -1997.4295 |
| $[Zn(Im)_3(HOCl)]^{2+}$ Complexed with $CH_3SH$ | -1997.4292 |
| $[Zn(Im)_3(CH_3SOH)]^{2+}$ Complexed with $H_2O$ | -1613.1302 |
| $[Zn(Im)_3(H_2O)]^{2+}$ Complexed with $CH_3SOH$ | -1613.1271 |
| $[Zn(Im)_3(CH_3SOH)]^{2+}$ Complexed with HOCl | -2072.6091 |
| $[Zn(Im)_3(HOCl)]^{2+}$ Complexed with $CH_3SOH$ | -2072.6084 |
| $[Zn(Im)_3(CH_3SO_2H)]^{2+}$ Complexed with $H_2O$ | -1688.3073 |
| $[Zn(Im)_3(H_2O)]^{2+}$ Complexed with $CH_3SO_2H$ | -1688.3117 |
| $[Zn(Im)_3(CH_3SO_2H)]^{2+}$ Complexed with HOCl | -2147.7806 |
| $[Zn(Im)_3(HOCl)]^{2+}$ Complexed with $CH_3SO_2H$ | -2147.7939 |
